# The brittle star genome illuminates the genetic basis of animal appendage regeneration

**DOI:** 10.1101/2023.10.30.564762

**Authors:** Elise Parey, Olga Ortega-Martinez, Jérôme Delroisse, Laura Piovani, Anna Czarkwiani, David Dylus, Srishti Arya, Samuel Dupont, Michael Thorndyke, Tomas Larsson, Kerstin Johannesson, Katherine M. Buckley, Pedro Martinez, Paola Oliveri, Ferdinand Marlétaz

**Author notes:** MRC London Institute of Medical Sciences, Imperial College London, London, UK. Technische Universität Dresden, Center for Regenerative Therapies Dresden (CRTD), Dresden, Germany. deceased (Michael Thorndyke).

## Abstract

Species within nearly all extant animal lineages are capable of regenerating body parts. However, it remains unclear whether the gene expression programme controlling regeneration is evolutionarily conserved. Brittle stars are a species-rich class of echinoderms with outstanding regenerative abilities, but investigations into the genetic bases of regeneration in this group have been hindered by the limited genomic resources. Here, we report a chromosome-scale genome assembly for the brittle star *Amphiura filiformis.* We show that the brittle star genome is the most rearranged amongst echinoderms sequenced to date, featuring a reorganised Hox cluster reminiscent of the rearrangements observed in sea urchins. In addition, we performed an extensive profiling of gene expression during brittle star adult arm regeneration and identified sequential waves of gene expression governing wound healing, proliferation and differentiation. We conducted comparative transcriptomic analyses with other invertebrate and vertebrate models for appendage regeneration and uncovered hundreds of genes with conserved expression dynamics, particularly during the proliferative phase of regeneration. Our findings emphasise the crucial importance of echinoderms to detect long-range expression conservation between vertebrates and classical invertebrate regeneration model systems.

## Introduction

Brittle stars are by far the most speciose class of echinoderms; over 2,600 extant species occupy benthic marine habitats globally (Stöhr et al. 2012; O’Hara et al. 2019). However, they remain poorly-documented from a genomic standpoint, despite their broad interest to diverse fields including marine (paleo)ecology, biodiversity monitoring, developmental biology and regenerative biology (Vistisen and Vismann 1997; Vopel et al. 2003; Dupont and Thorndyke 2007; Mosher and Watling 2009; Thuy et al. 2012; Delroisse et al. 2017; Dylus et al. 2018; O’Hara et al. 2019).

The echinoderm phylum encompasses five classes with a well-resolved phylogeny (Figure 1A; (Cannon et al. 2014; O’Hara et al. 2014; Telford et al. 2014; Mongiardino Koch et al. 2022)): brittle stars (Ophiuroidea), sea stars (Asteroidea), sea urchins (Echinoidea), sea cucumbers (Holothuroidea) and sea lilies/feather stars (Crinoidea). Genomics in this phylum began with the pioneering effort to sequence the genome of the purple sea urchin (*Strongylocentrotus purpuratus*) (Sea Urchin Genome Sequencing Consortium et al. 2006). Analysis of this genome provided broad insights into the evolution of diverse traits and biological processes, including for instance biomineralization, sensory capabilities and immune recognition (Livingston et al. 2006; Raible et al. 2006; Rast et al. 2006). In recent years, the taxonomic sampling of echinoderm genomes has steadily expanded (Hall et al. 2017; Zhang et al. 2017; Davidson et al. 2020; Lawniczak et al. 2021; Davidson et al. 2022; Chen et al. 2023; Marlétaz et al. 2023). This growing wealth of genomic resources in the context of the remarkable diversity of echinoderm body plans, life histories and developmental strategies, provides a unique framework to understand the evolution of novel traits. However, given the deep evolutionary divergence of the five echinoderm classes (480-500 Mya), the lack of robust genomic resources for the brittle stars represents a problematic knowledge gap.

**Figure 1:**
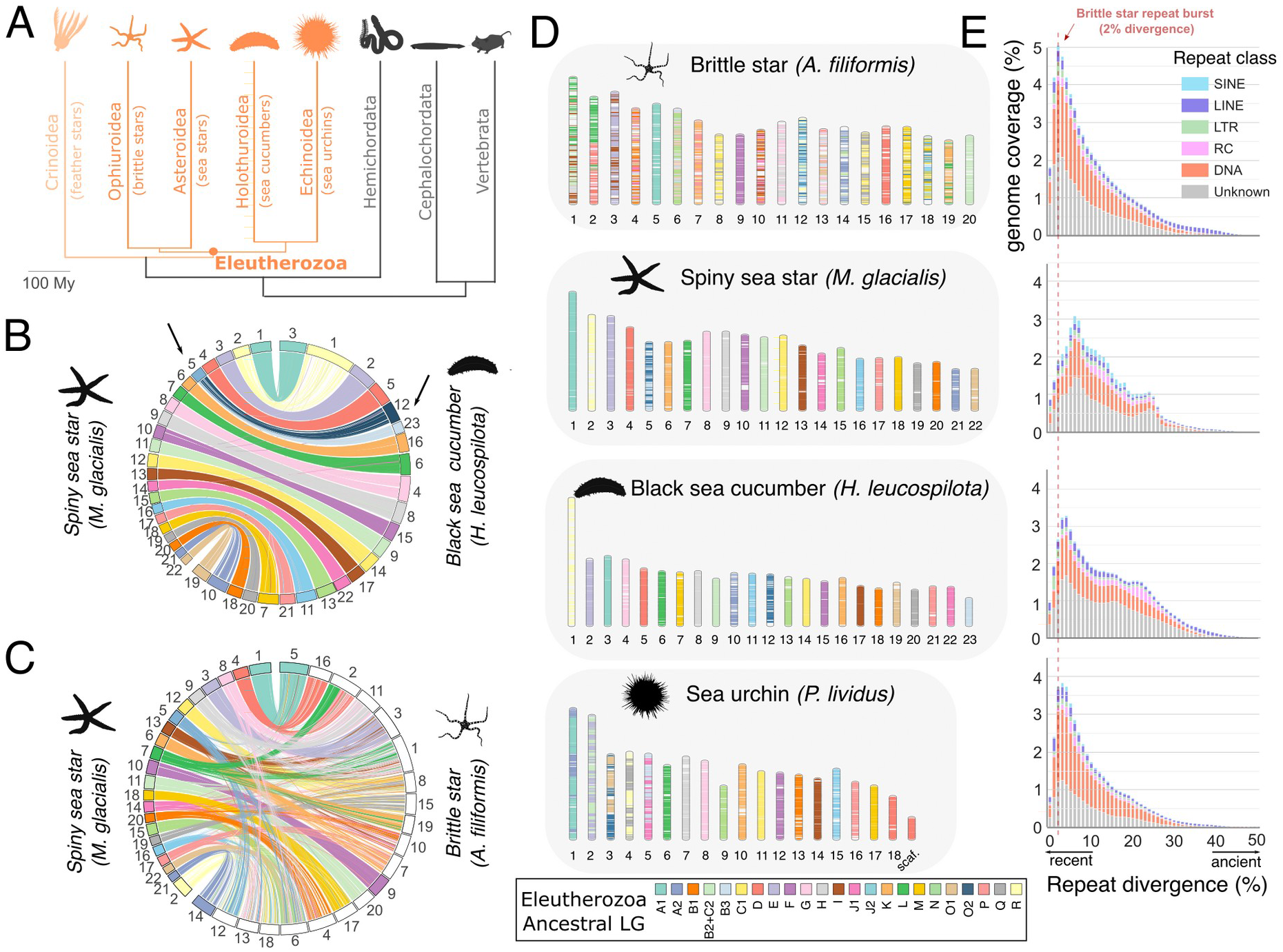
Chromosome evolution in echinoderms. **A.** Phylogenetic relationships of the five echinoderm classes (orange), with the position of the Eleutherozoa ancestor highlighted, and hemichordates and chordates as outgroups. Classes with available chromosome-scale genome assembly are shown in dark orange. Divergence times amongst echinoderms and with hemichordates were extracted from (Mongiardino Koch et al. 2022), divergence with chordates from TimeTree (Kumar et al. 2017). **B.** Synteny comparison between the 22 chromosomes of spiny sea star and the 23 chromosomes of the black sea cucumber. The single macrosyntenic rearrangement between the two genomes is indicated with arrows. **C.** Synteny comparison between the 22 chromosomes of spiny sea star and the 20 chromosomes of brittle star. The three brittle star chromosomes with a 1-to-1 relationship with sea star chromosomes are shown with a colour matching its orthologous counterpart in spiny sea star. **D.** Chromosome evolution in Eleutherozoa. We named the ancestral Eleutherozoa linkage groups (ELG) using established naming conventions proposed for the 24 bilaterian ancestral linkage groups defined in (Simakov et al. 2022; Marlétaz et al. 2023). B2+C2 corresponds to a fusion of bilaterian B2 and C2 present in the Eleutherozoa ancestor. **E.** Repeat landscapes for the brittle star and the three selected echinoderm genomes, with the y-axis representing the genomic coverage and x-axis CpG-corrected Kimura divergence to the repeat consensus. Species are presented in the same order as in **D**. The dashed red line indicates the repeat burst in the brittle star.

Adult echinoderms share a characteristic pentameral symmetry, which represents the most derived body plan amongst Bilateria (Smith 2008). Early analyses of sea urchin genomes unveiled local reorganisations within the Hox cluster, prompting speculation that they were associated with the evolution of this unique body plan (Lowe and Wray 1997; Cameron et al. 2006; Mooi and David 2008; David and Mooi 2014). However, the subsequent discovery of an intact Hox cluster in the crown-of-thorns sea star revealed that these rearrangements were not instrumental in the establishment of the pentameral symmetry (Baughman et al. 2014; Byrne et al. 2016). These observations showcase the need to examine a more comprehensive sample of echinoderm whole genomes to accurately identify echinoderm-specific chromosomal rearrangements and subsequently investigate their functional significance.

Echinoderms exhibit extensive regenerative abilities. Species from each of the five classes are capable of varying levels of regeneration, including (larval) whole-body regeneration, appendage or organ regeneration (Medina-Feliciano and García-Arrarás 2021). Although species within nearly all major animal groups exhibit some regenerative capacity, it is not clear whether this trait is ancestral or independently acquired through convergent evolution (Bely and Nyberg 2010; Lai and Aboobaker 2018; Srivastava 2021). A comparative analysis of whole-body regeneration across a sea star larva, planarian worm and hydra has suggested that broadly-conserved molecular pathways may mediate regeneration (Cary et al. 2019). However, given the diversity of regenerative modes, additional comparative analyses of regenerating organisms are needed to fully understand the evolution of this complex process (Lai and Aboobaker 2018; Srivastava 2021). In particular, gene expression dynamics during regeneration have not been explicitly compared between invertebrates and vertebrates, partly because of the lack of gene expression profiling across comparable regenerating structures and of difficulties in identifying orthologs among distant model systems. Echinoderms are more closely related to vertebrates than other classical invertebrate models of regeneration, hence providing a unique phylogenetic perspective. Despite recent studies in sea stars and sea cucumbers (Fumagalli et al. 2018; Cary et al. 2019; Quispe-Parra et al. 2021), echinoderms remain largely underrepresented in transcriptomic assays of regeneration (Dupont and Thorndyke 2007; Goldman and Poss 2020; Bideau et al. 2021).

One highly regenerative echinoderm species is the brittle star *Amphiura filiformis,* likely related to their sediment dwelling lifestyle where extended arms are often severed by predators. In this species, fully differentiated arms regrow in a few weeks following amputation and over 90% of individuals sampled in the wild display signs of arm regeneration (Duineveld and Van Noort 1986; Sköld and Rosenberg 1996). Consequently, *A. filiformis* is emerging as a powerful model for animal appendage regeneration, with a well-established morphological staging system (Dupont and Thorndyke 2006; Czarkwiani et al. 2013; Hu et al. 2014; Purushothaman et al. 2015; Czarkwiani et al. 2016; Piovani et al. 2021; Czarkwiani et al. 2022). Here, we report a chromosome-scale genome assembly for the brittle star *A. filiformis* and leverage this unique resource to investigate the complex history of karyotypes, Hox cluster and gene family evolution across echinoderms. We find that *A. filiformis* displays the most rearranged echinoderm genome sequenced to date. Moreover, we reveal that *A. filiformis* extensive regenerative capacities correlate with significant expansions of genes involved in wound healing. Finally, we generate extensive transcriptomic data from regenerating brittle star arms, which we analyse in a comparative framework with previously generated datasets from the crustacean *Parhyale hawaiensis* (Sinigaglia et al. 2022) and the axolotl *Ambystoma mexicanum* (Stewart et al. 2013), to illuminate common genetic mechanisms of animal appendage regeneration.

## Results

### The *Amphiura filiformis* chromosome-scale assembly is a key genomic resource

To address the lack of high-quality genome for the brittle stars (**Supp. Note 1**), we sequenced and assembled the genome of the brittle star *Amphiura filiformis* using high-coverage long nanopore reads assisted with proximity ligation data for scaffolding (**Methods**). The haploid assembly spans 1.57 Gb and contains 20 chromosome-size scaffolds (>60 Mb) that account for 93.5% of the assembly length (**Figure S1,** N50: 68.8 Mb). This is consistent with cytogenetic studies in two other brittle star species that identified 21-chromosome karyotypes (Colombera and Venier 1976; Saotome and Komatsu 2002). We annotated a total of 30,267 protein-coding genes (92.7% complete BUSCO score, **Methods, Table S1**, **Table S2**), which is in line with the predicted gene complements of other echinoderms (Hall et al. 2017; Li et al. 2020; Davidson et al. 2022; Chen et al. 2023; Marlétaz et al. 2023). In addition, we generated manually-curated lists for *A. filiformis* genes associated with immunity, stemness and neuronal function as well as transcription factors and genes involved in 19 major signalling pathways (**Table S2**, **Methods**). These lists allow for genome-wide interrogation of *A. filiformis* genes as a complement to gene ontology-based approaches. In summary, the *A. filiformis* genome represents the first high-quality and chromosome-scale genome assembly for the brittle star class, and fills an important knowledge gap in the echinoderm genomics landscape.

### The brittle star exhibits the highest inter-chromosomal rearrangement rate amongst sequenced echinoderms

Chromosome evolution in echinoderms has primarily been investigated through the lens of sea urchin genomes, which have globally preserved the ancestral bilaterian chromosomes that were previously reconstructed based on comparisons between chordates and molluscs (Simakov et al. 2020; Simakov et al. 2022; Marlétaz et al. 2023). However, sea urchins also underwent several chromosomal fusions whose origin cannot be established without examining more echinoderm genomes. To pinpoint the timing of these fusions within the context of echinoderm evolution, and to evaluate the conservation of the ancestral chromosomal units across echinoderm lineages, we took advantage of chromosome-scale genomes released for sea stars, sea cucumbers and sea urchins (Lawniczak et al. 2021; Chen et al. 2023; Marlétaz et al. 2023). Using these genomes in combination with the brittle star genome and selected outgroups, we reconstructed the linkage groups present in their ancestor (Eleutherozoa linkage groups, or ELGs, **Figure 1A**).

Synteny comparisons between the spiny sea star (*Marthasterias glacialis*) and the black sea cucumber (*Holothuria leucospilota*) (Lawniczak et al. 2021; Chen et al. 2023) reveal that only one inter-chromosomal macrosyntenic rearrangement occurred in the 500 million years (My) of independent evolution between these two genomes (**Figure 1B, Methods**). In striking contrast, the *A. filiformis* brittle star genome is extensively rearranged: only three chromosomes have a direct one-to-one orthology relationship with spiny sea star chromosomes (**Figure 1C**, Fisher’s exact test adjusted p-value<10^-5^). We reconstructed the ancestral ELGs based on near-perfect conservation of macrosynteny between the spiny sea star and black sea cucumber and using amphioxus (*Branchiostoma floridae*) and sea scallop (*Pecten maximus*) as outgroups to disentangle derived and ancestral chromosomal arrangements (**Figure S2**). We predicted that 23 ELGs were present in the Eleutherozoan ancestor (**Figure 1D**). The ELGs descend from the 24 bilaterian linkage groups (BLGs) (Simakov et al. 2022) through the fusion of the BLGs B2 and C2. Among echinoderms, the black sea cucumber maintained the 23 ancestral ELGs, a single chromosomal fusion took place in the spiny sea star lineage (corresponding to an inter-chromosomal rearrangement rate of 0.002 event / My), five fusions occurred in the sea urchin *Paracentrotus lividus* (0.01 event / My) and 26 inter-chromosomal rearrangements shaped the brittle star *A. filiformis* karyotype (0.052 event / My, the highest amongst sequenced echinoderms; **Figure S3**). These results indicate that, among Eleutherozoa, sea cucumbers and sea stars show the strongest conservation of ancestral bilaterian linkage groups, whereas the brittle star genome is highly reshuffled relative to the Bilaterian ancestor. Examination of additional sea stars and sea urchins genomes suggest that these trends might broadly extend to species within their respective classes ((Davidson et al. 2023; Liu et al. 2023; Marlétaz et al. 2023); **Figure S3**), but, given the limited sampling, this should be re-examined as more chromosome-scale genome assemblies become available.

One potential driver of genomic rearrangements is transposable elements and repetitive sequences, which serve as substrates for non-allelic homologous recombination (George and Alani 2012; Balachandran et al. 2022). Transposable elements have also been implicated in driving changes in genome size (Canapa et al. 2015). Among the four echinoderm genomes analysed, we find that repetitive elements coverage correlates as expected with genome size but not with rates of rearrangements. Repeat coverage is highest in the highly-rearranged brittle star genome (1.57 Gb, repeat coverage 59.3%) and slowly-evolving black sea cucumber *H. leucospilota* (1.31 Gb, 56.0%) compared to the sea urchin *P. lividus* (927 Mb, 49.2%) and spiny sea star *M. glacialis* (521 Mb, 47.6%). Analysis of sequence divergence reveals that repetitive elements accumulated more gradually in the slowly-evolving sea star and sea cucumber genomes, compared to both the sea urchin and the brittle star which display recent bursts of repeat activity (**Figure 1E**). Specifically, the brittle star genome is marked by a burst of repeat activity 10-15 Mya, consisting mostly of DNA transposons (peak of repeats with 2% divergence to consensus, **Methods**). We thus speculate that the evolutionary history of *A. filiformis* includes at least one period of genomic instability (Belyayev 2014). Together, these data highlight contrasting trends of chromosome evolution across echinoderm classes, and indicate that *A. filiformis* is the most rearranged echinoderm genome among those sequenced to date.

### The brittle star Hox cluster is marked by small-scale genomic rearrangements

The organisation of the Hox and ParaHox gene clusters has been documented in each class of echinoderms with the exception of brittle stars (Cameron et al. 2006; Annunziata et al. 2013; Baughman et al. 2014; Zhang et al. 2017; Li et al. 2020). To further explore the enigmatic evolution of these developmental homeobox gene clusters in echinoderms (Byrne et al. 2016), we investigated the structure of the *A. filiformis* Hox and ParaHox clusters. Strikingly, the *A. filiformis* Hox and ParaHox clusters both exhibit genomic rearrangements (**Figure 2, Figure S4, Methods**): anterior Hox genes (*Hox1*, *Hox2* and *Hox3*) are inverted within the 3’ end of the cluster and a transposition/inversion event occurred that displaced *Hox8* between *Hox9/10* and *Hox11/13a.* Five repeat families are significantly expanded within the brittle star Hox cluster, of which one is significantly associated with breakpoint locations and may have contributed to the Hox1-3 inversion through non-homologous repair (SINE/tRNA-Deu-L2, BH-corrected permutation-based p-value <0.05, **Figure 2B**). Four out of five of expanded repeats have an inferred divergence of 18-22% to their consensus, suggesting they were active approximately 100 Mya (**Methods**). While brittle star Hox reorganisation is convergent and distinct from the one observed in sea urchins, in both cases the orientation of *Hox1*, *Hox2* and *Hox3* is reversed relative to the other genes of the cluster, and, in both lineages, one of the breakpoints is located near *Hox4* (**Figure 2C**). Moreover, mirroring the evolution of the sea urchins ParaHox genes (**Figure 2D**), the brittle star ParaHox cluster also underwent disruptions, such that *Gsx* was tandemly duplicated to generate two paralogs (protein identity: 74%) located a long distance (>5 Mb) from *Xlox*-*Cdx*. Whereas *Xlox*-*Cdx* maintained close linkage in the brittle star, all three members of the ParaHox cluster are dispersed over their chromosome in sea urchins (Annunziata et al. 2013).

**Figure 2:**
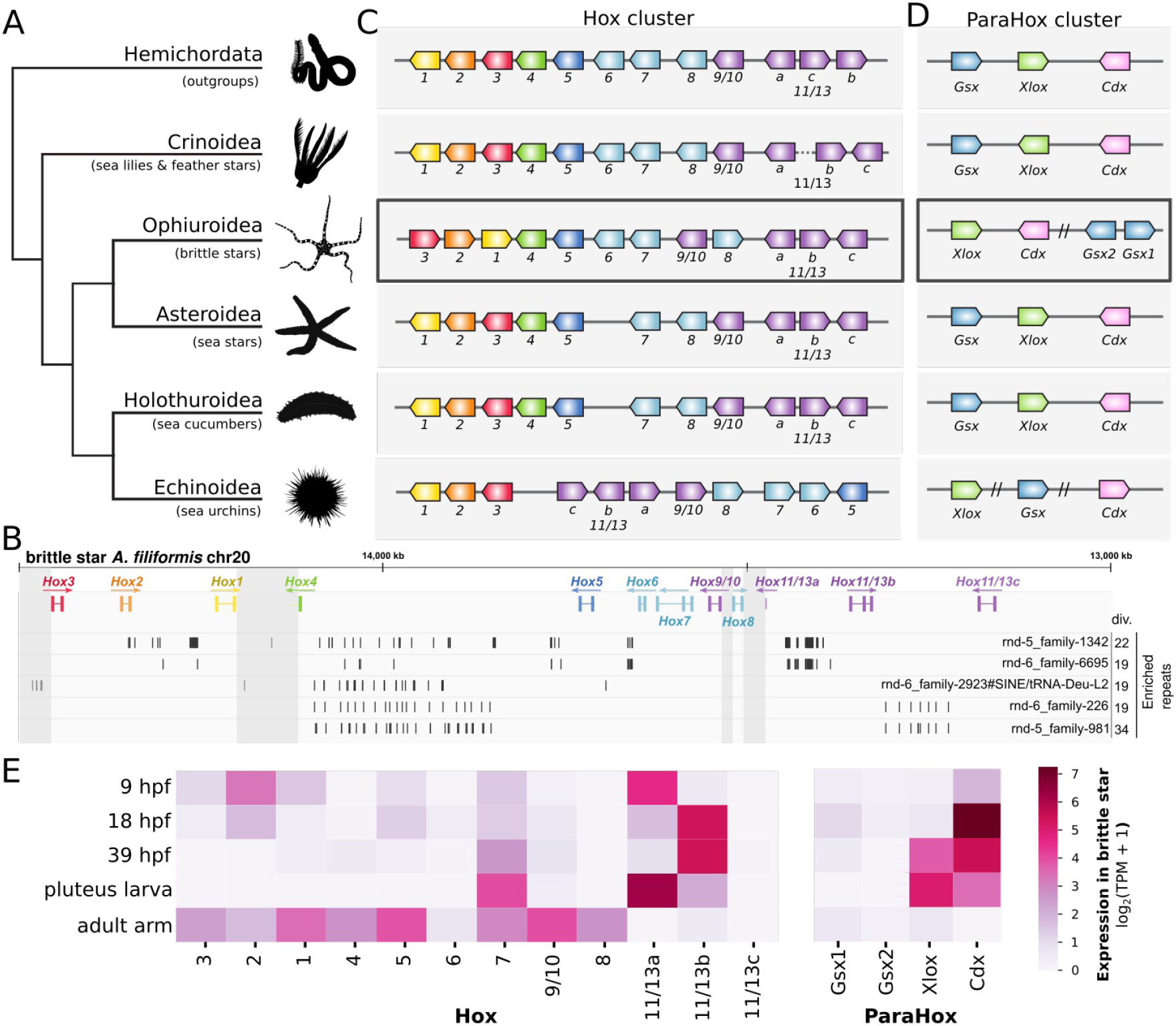
Hox and ParaHox clusters organisation across echinoderms. **A.** Phylogenetic relationships between the five classes of echinoderms, with hemichordates as the outgroup. **B.** Genomic organisation of the brittle star *A. filiformis* Hox cluster. Significantly expanded repeats at the Hox cluster location are represented in their respective tracks below Hox genes, with the average sequence divergence to consensus indicated (div., %). Divergence to consensus is a proxy for repeat age, where higher divergence indicates older repeat insertions. Vertical grey rectangles indicate breakpoint locations. **C.** Schematic representation of Hox cluster organisation across echinoderms and outgroups, based on organisation reported in *S. kowalevskii* and *P. flava* (Freeman et al. 2012) for Hemichordata, feather star *A. japonica* (Li et al. 2020) for Crinoidea, brittle star *A. filiformis* for Ophiuroidea, crown-of-thorns sea star *A. planci* (Baughman et al. 2014; Hall et al. 2017) for Asteroidea, Japanese sea cucumber *A. japonicus* (Kikuchi et al. 2015; Zhang et al. 2017) for Holothuroidea and purple sea urchin *S. purpuratus* (Cameron et al. 2006) for Echinoidea. **D.** ParaHox gene cluster organisation, based on the same genomes as in B. Double slashes indicate non-consecutive genes, all separated by distances > 5 Mb on the same chromosome or scaffold. **E.** Expression of Hox and ParaHox genes throughout 4 brittle star developmental time points and in the adult arm (hpf: hours post-fertilisation). Expression data from (Delroisse et al. 2014; Delroisse et al. 2015; Dylus et al. 2016) was normalised across samples using the TMM method (Robinson et al. 2010) on the full set of brittle star genes, and shown as log_2_(TPM + 1).

Hox expression throughout echinoderm embryogenesis, larval stages and metamorphosis remain largely enigmatic, such that spatio-temporal expression does not follow classical Hox collinearity rules (Arenas-Mena et al. 1998; Byrne et al. 2016). We investigated Hox and ParaHox gene expression in the brittle star using previously published datasets throughout four developmental time points (Delroisse et al. 2014; Delroisse et al. 2015; Dylus et al. 2016) (**Figure 2E, Table S1, Methods**). As in sea urchins (Arenas-Mena et al. 1998), *Hox1* and *Hox3-6* are expressed at very low levels in the brittle star embryos and pluteus larvae (normalised TPM < 2), whereas *Hox7*, *Hox11/13a* and *Hox11/13b* are highly expressed. One notable difference between the expression of Hox genes in the two lineages is seen in *Hox2*. In the brittle star, *Hox2* is expressed early in embryogenesis, with maximal expression at 9 hours post-fertilisation (developmental stage). In contrast, sea urchins *Hox2* is not expressed during early development (Arenas-Mena et al. 1998; Tu et al. 2014). Furthermore, the brittle star ParaHox genes *Xlox* and *Cdx* are each expressed during early development whereas the anterior *Gsx1 and Gsx2* genes are not (**Figure 2E**), matching the expression patterns observed in sea stars (Annunziata et al. 2013). In contrast, dispersion of the ParaHox cluster in sea urchins is associated with the distinct temporal activation of *Gsx*, *Xlox* and *Cdx* during embryogenesis (Arnone et al. 2006).

These results highlight intriguing parallels in the reorganisation of developmental gene clusters and their expression patterns between brittle stars and sea urchins. Limited data are available on Hox gene expression in other echinoderm classes, but investigations in crinoids and sea cucumbers suggest that, even in species with an intact Hox cluster, the anterior genes (Hox1-6) exhibit low or no expression in early embryonic stages (Hara et al. 2006; Kikuchi et al. 2015; Li et al. 2018), and that *Hox7* and *Hox11/13b* may play an important role in embryogenesis (Annunziata et al. 2014). We therefore speculate that the relaxation of expression constraints on Hox genes during echinoderm embryogenesis may have allowed for the rearranged Hox cluster architectures seen in the sea urchins and brittle star lineages.

### Tandem duplications of key genes contribute to brittle star larval skeleton and bioluminescence

Tandem gene duplications and subsequent asymmetric divergence are widespread in the evolution of animal genomes and have been linked to the evolution of species-specific traits (Holland et al. 2017). In echinoderms, two specific gene families represent relevant examples of lineage-specific evolution through tandem duplications: *phb*/*pmar1* (larval skeleton) and luciferases (bioluminescence) (Dylus et al. 2016; Delroisse et al. 2017; Marlétaz et al. 2023). Pmar1 is the most upstream zygotic factor of the regulatory network controlling the specification of skeletogenic cells in sea urchins (Oliveri et al. 2003; Oliveri et al. 2008). Among Eleutherozoa, only sea urchins and brittle stars develop an elaborated larval skeleton. In sea urchins, the *pmar1* gene originated through repeated lineage-specific duplications of an ancient *phb* paired-class homeobox gene. Duplications of the *pmar1* gene have been pinpointed as important drivers for the establishment of this sea urchin-specific regulatory programme, which culminates in the formation of the larval skeleton (Dylus et al. 2016; Yamazaki et al. 2020; Marlétaz et al. 2023). In the brittle star *A. filiformis*, we identify a similar expansion of *phb* paralogs (totaling 13 *phb* genes). Phylogenetic analysis confirms that these *phb* paralogs are distant homologs of the sea urchins *pmar1*, but indicates that they are distinct from the previously described brittle star *pplx* gene (Dylus et al. 2016) (**Figure S5A**). Moreover, expression of the *A. filiformis phb* genes occur largely during early development (**Figure S5E**), as described for the *pmar1* sea urchin gene. This suggests that the convergent evolution of brittle stars and sea urchins larval skeleton may have been driven by independent duplications of *phb* genes.

Brittle stars stand out by their ability to emit light (Mallefet 2009). In *A. filiformis*, bioluminescence is mediated by a specific type of luciferase which is homologous to the well-characterised luciferase of the soft coral *Renilla reniformis* (Delroisse et al. 2017). Within the *A. filiformis* genome, we identified nine luciferase-like gene copies: seven are organised in two clusters of tandem duplicates, two are isolated copies. This corroborates and extends the previously-inferred repertoire (Delroisse et al. 2017). We find that luciferase-like genes have duplicated not only in the brittle star but also in all echinoderm lineages with the exception of sea stars (**Figure S5**). These results confirm previous propositions that echinoderms harbour multiple copies of luciferase-like genes, which likely encode diverse functions across bioluminescent and non-bioluminescent species (Delroisse et al. 2017).

### Expanded gene families in echinoderms are enriched in regeneration-related processes

To comprehensively assess the functional significance of gene complement evolution, we inferred gene family expansion and contraction events within echinoderms (**Figure 3A, Methods**). In contrast with other deuterostome lineages, which exhibit either extensive gene losses (Seo et al. 2001) or duplications (Putnam et al. 2008), we found that echinoderms harbour relatively stable gene complements, with only ∼7.6% of gene families showing significant expansions or contractions in their ancestral lineage and throughout their evolution (790 of 10,367 tested families). Within the brittle star, genes in these families are enriched in GO terms that are also found in the expanded and contracted families of other echinoderm classes (**Figure 3B, Table S3, Methods**). This includes several enriched GO terms linked to immune-related processes (e.g. “response to other organisms”, “leukocyte migration”, “cell recognition”), which encompass genes known to display elevated gene birth and death rates in other animal lineages, such as Toll-Like Receptors (Nei et al. 1997; Leulier and Lemaitre 2008; Saco et al. 2023). Some GO term enrichments may reflect specific aspects of echinoderm biology. For instance, recurrent duplications of “regeneration-related” genes may underlie the remarkable regenerative capacity of many echinoderm species. Notably, in *A. filiformis,* members of these expanded gene families (**Figure 3C**) are expressed during arm regeneration (**Figure S6**). Additionally, genes within four expanded families (*plasminogen*, *carboxypeptidase B*, *coagulation factor* and *ficolin*) directly regulate coagulation and/or clotting in vertebrates (Pryzdial et al. 2022), but may play a broader role in immune defence in echinoderms (Hanington and Zhang 2011; Loof et al. 2011). Moreover, the *ficolin* gene has also been implicated in the early stages of *A. filiformis* arm regeneration (Ferrario et al. 2018; Arenas Gómez et al. 2020). Duplications within the brittle star may thus have contributed to the evolution of a rapid and efficient wound closure process that is prerequisite to regeneration (Suárez-Álvarez et al. 2016; Ferrario et al. 2018). Finally, genes involved in keratan sulfate metabolism are over-represented in both expanded and contracted gene families in the brittle star (**Figure 3B**). Increased sulfated glycosaminoglycans production is required for proper arm regeneration in *A. filiformis* (Ramachandra et al. 2017). The numerous duplications and losses of these genes suggest that the evolution of brittle star efficient regeneration may have been accompanied by a specialisation of the glycosaminoglycan metabolism.

**Figure 3:**
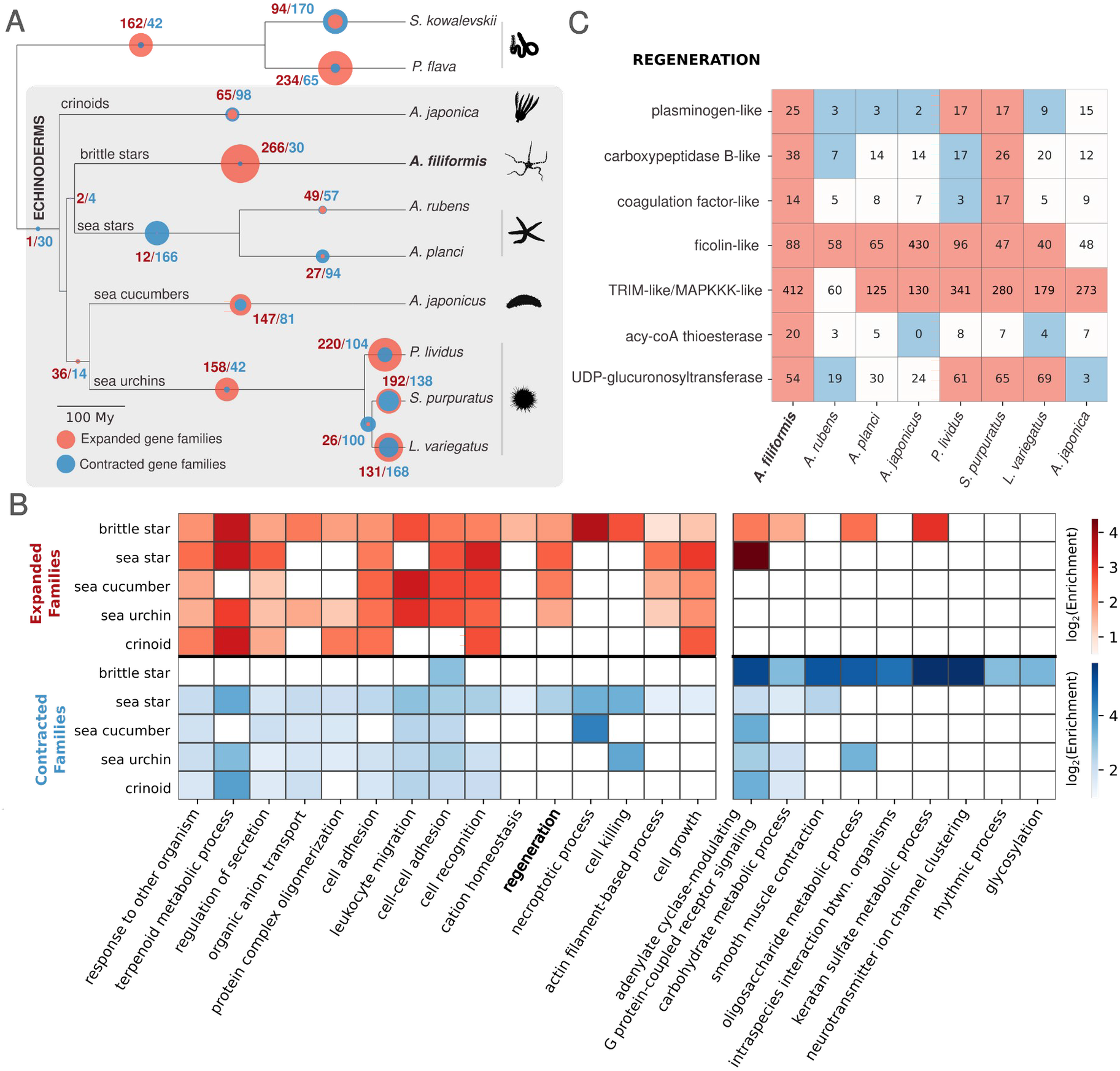
Gene family evolution in echinoderms. **A.** Number of significantly expanded (red) and contracted (blue) gene families throughout echinoderm evolution, from a total of 10,367 tested gene families (**Methods**). **B.** Gene ontology functional enrichment tests (Biological Process) for expanded and contracted families in the different echinoderm classes. We selected the top 15 representative GO terms enriched in the expanded brittle star gene families and 10 in contracted families (**Methods**). In the heatmap, colours indicated GO terms significantly enriched in expanded or contracted families in other echinoderm classes (FDR < 0.05). **C.** Gene copy number variation across echinoderms for regeneration gene families with significant expansion in *A. filiformis* (>1 brittle star gene in the family annotated with the GO term ‘regeneration’). Gene families were named according to the *S. purpuratus* gene name. Red and blue colours denote significantly expanded and contracted families, respectively.

### Gene expression during brittle star arm regeneration recapitulates major regeneration phases

To gain insight into the transcriptional programmes that underlie brittle star arm regeneration, we profiled gene expression in seven representative regeneration stages following amputation and one non-regenerating control (**Methods**). Using soft-clustering, we classified genes into nine major temporal clusters (A1-A9; **Figure 4A**, **Figure S6, Methods**). Functional enrichment analysis of genes within the co-expression clusters revealed three distinct phases of arm regeneration: (1) wound healing, (2) proliferation and (3) tissue differentiation. These are consistent with morphological timelines of regeneration in the brittle star and other animal systems (Czarkwiani et al. 2016; Bideau et al. 2021; Srivastava 2021).

**Figure 4:**
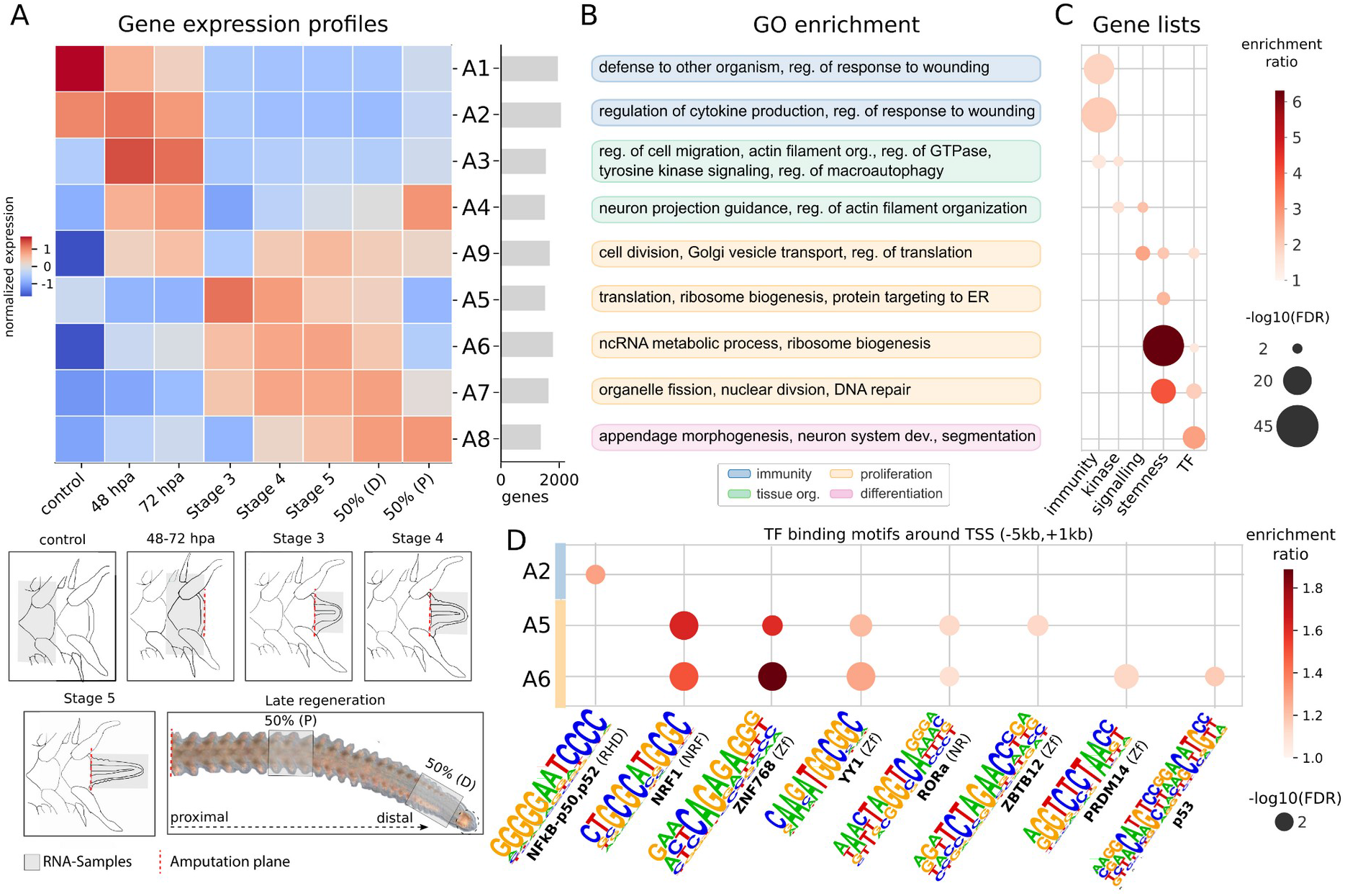
Gene expression during brittle star arm regeneration. **A.** Soft-clustering of gene expression profiles throughout regeneration time points, yielding 9 main temporal co-expression clusters (A1-A9, **Methods,** see also **Figure S6**). Co-expression clusters are temporally ordered (from top to bottom) on the basis of their first expression time point. Barplots on the right indicate the number of genes assigned to each cluster. The RNA-sampling procedure for each stage is illustrated below. Early stages are sampled at 48 and 72 hpa (hours post-amputation). Subsequent stages are defined by morphological landmarks: Stage 3 corresponds to the presence of the radial nerve (∼6 days post-amputation, dpa), Stage 4 is the appearance of the first regenerated metameric units (∼8 dpa), Stage 5 corresponds to advanced arm extension and differentiation onset (∼9 dpa), 50% stages correspond to when 50% of the regenerated arm has differentiated (∼2-3 weeks post-amputation) sampled at the distal (D, less differentiated) and proximal (P, more differentiated) ends (Dupont and Thorndyke 2006; Czarkwiani et al. 2016). **B.** Gene ontology enrichment for each co-expression cluster (**Methods**, see **Figure S7** and **Table S4** for exhaustive GO results). **C.** Curated gene lists enrichment for each co-expression cluster (hypergeometric test, Benjamini-Hochberg adjusted p-values <0.05, **Methods, Table S2**). **D.** TF binding motifs enriched around the TSS (5kb upstream to 1kb downstream) of genes from co-expression clusters (hypergeometric test adjusted p-value <0.05, **Methods**).

Early regeneration is marked by the expression of genes involved in wound response, including immunity/wound healing (clusters A1-A2), and cell migration/tissue protection (clusters A3-A4), which are enriched in immune and kinase genes, respectively (**Figure 4B, 4C**). Notably, the regions surrounding transcription start sites (TSS) of genes within cluster A2 are enriched for transcription factor (TF) binding motifs of NF-κB, a broadly conserved regulator of immune response (**Figure 4D**).

Wound healing is followed by cell proliferation (clusters A9 and A5, A6, A7), as indicated by the overrepresentation of stemness genes and genes involved in cell proliferation, cell division and enhanced translational activity. Accordingly, binding motifs associated with several proliferation-related TF are enriched around the TSS of genes from clusters A5 and A6. This includes NRF1 and p53, which have been implicated in vertebrates in regulating (stem) cell survival and proliferation (Cui et al. 2021; Ayaz et al. 2022), PRDM14 and YY1, which regulate pluripotency (Kawaguchi et al. 2019; Dong et al. 2022), and RORa, which controls inflammation by down-regulating targets of NF-κB (Oh et al. 2019) and may thus play a role in the transition from wound response to proliferation (**Figure 4C, 4D**). We also find enrichment of binding motifs corresponding to zinc-finger transcription factors that are involved in cell proliferation and pluripotency (Villot et al. 2021; Han et al. 2023). Cluster A9 encompasses genes expressed as early as 48 hours post-amputation (hpa) and active throughout regeneration, including translational regulators, cell division and vesicle transport genes (**Figure 4B**), as well as genes involved in signalling pathways that promote cell proliferation (VEGF, Akt, Insulin-like and Jak-STAT pathways, **Figure 4C, Figure S6**) (Xu et al. 2012; Huat et al. 2014; Apte et al. 2019; Herrera and Bach 2019). These data suggest that the signalling cascades that initiate cell proliferation are induced very early during brittle star regeneration (cluster A9); they are activated during the wound response phase and exhibit amplified expression during the peak of cell proliferation (Stage 5; **Figure 4A**). The early onset of proliferation (around 48 hpa) is consistent with previous observations of cell proliferation and expression quantification of selected marker genes (Czarkwiani et al. 2016; Czarkwiani et al. 2022).

Finally, late regeneration is characterised by the expression of genes involved in differentiation, patterning and appendage morphogenesis, with a significant over-representation of transcription factors (cluster A8, **Figure 4B, 4C**). This cluster includes two T-box TFs that are important for axis specification in echinoderms (*tbx3-1* and *tbx3-2*) and two TFs with key roles in neurogenesis (*ngn1-like* and *hey1-like*) (Gross et al. 2003; Slota and McClay 2018; Slota et al. 2019).

Overall, these data provide a genome-wide picture of the molecular pathways at play throughout brittle star arm regeneration and highlight three waves of gene expression that successively mobilise genes involved in wound response, cell proliferation and tissue differentiation. These general phases have been described in many regenerating animals. Consequently, this dataset can be leveraged to assess the conservation of regeneration gene expression dynamics across species.

### Animal appendage regeneration is characterised by conserved gene expression during the cell proliferation phase

Several key genes and pathways have been repeatedly implicated in regeneration across animal lineages (Bideau et al. 2021; Srivastava 2021), However, direct comparisons of temporal expression gene profiles throughout regeneration remain limited. Here, we investigate the conservation of the animal appendage regeneration programmes through the lens of the brittle star *A. filiformis*.

Using a genomic phylostratigraphy approach (Barrera-Redondo et al. 2023), we found that, overall, brittle star arm regeneration is mediated by ancient genes (i.e. metazoan or older; **Figure 5A, Methods**). The exception is the initial wound-healing phase, which is enriched in genes that are specific to the brittle star lineage (**Figure 5A**). The observation that brittle star regeneration is mostly driven by ancient genes prompted us to investigate whether these genes are similarly involved in appendage regeneration across animals, and whether they are deployed in the same temporal order. As an invertebrate deuterostome, brittle stars enable phylogenetic comparisons among vertebrate and ecdysozoa appendage regeneration models. We thus compared gene expression dynamics during appendage regeneration in *A. filiformis* with comparable datasets from the axolotl (*Ambystoma mexicanum*) (Stewart et al. 2013) and the crustacean Parhyale (*Parhyale hawaiensis*) (Sinigaglia et al. 2022). For this analysis, we defined nine major co-expression clusters during axolotl limb regeneration (Ax1-Ax9, **Figure S8, Table S5**) that effectively recapitulate the three regeneration phases (wound healing, proliferation and differentiation), and used existing Parhyale clustering (Sinigaglia et al. 2022).

**Figure 5:**
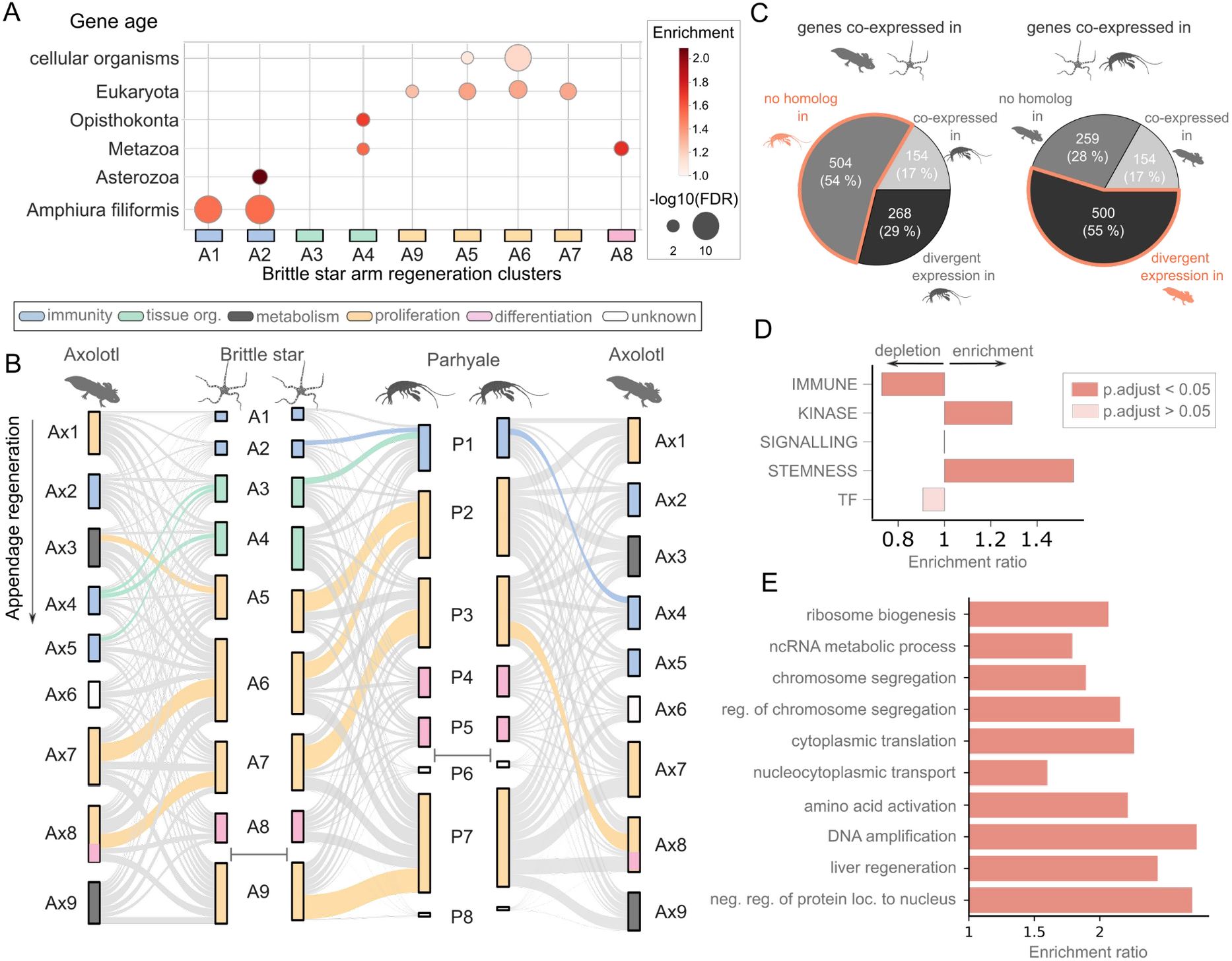
Gene expression throughout appendage regeneration across animals. **A**. Gene age enrichments for brittle star arm regeneration clusters (hypergeometric test, Benjamini-Hochberg adjusted p-values <0.05). Clusters are ordered by the time of expression onset. **B.** Comparison of co-expressed gene clusters deployed during appendage regeneration in axolotl, brittle star and Parhyale (left to right: axolotl vs brittle star, brittle star vs Parhyale, Parhyale vs axolotl). Clusters in Parhyale (clusters P1 to P8) correspond to the clustering reported in (Sinigaglia et al. 2022), but clusters were renamed to follow temporal activation and homogenise with respect to brittle star and axolotl clusters (**Methods**). Co-expression clusters in each species are shown in order of their temporal expression (from top to bottom), with the exception of brittle star cluster A9 and Parhyale clusters P6-P7-P8 that are expressed throughout several regeneration time points and shown at the bottom. Clusters are represented by vertical rectangles whose sizes are proportional to the number of homologous genes in the cluster, and coloured according to enriched GO terms (**Methods**, Figure 4, **Figure S8**, see **A** for legend). Links between clusters of the two compared species indicate cluster membership of homologous genes, with coloured links indicating significant overlaps (permutation-based p-values with Benjamini-Hochberg correction <0.05, **Methods**). **C.** The majority of genes identified as co-expressed in the brittle star - Parhyale and brittle star - axolotl comparisons are not recovered in the direct Parhyale - axolotl comparison. The majority of genes co-expressed in the axolotl and brittle star have no identified homologs in Parhyale (54%, left pie chart). Genes co-expressed in Parhyale and the brittle star have a divergent expression in the axolotl, i.e. they are not found in matched co-expression clusters (55%, right pie chart). **D.** Gene lists enrichment and depletion tests performed for the set of brittle star genes with conserved temporal expression during animal regeneration (**Methods**). **E.** Gene ontology enrichment tests, as in **D.**

Pairwise comparisons and permutation tests reveal that many of the co-expression clusters employ homologous genes during appendage regeneration across the three species (**Figure 5B**, **Methods**). Among the nine co-expression clusters that mediate brittle star regeneration, five consist of genes that are also co-expressed during axolotl regeneration (926 genes), six clusters overlap with Parhyale (913 genes), and four clusters are consistent across the three species (154 genes) (**Figure 5B, 5C, Table S6**). Expression comparisons between the more phylogenetically distant axolotl and Parhyale identify only two conserved co-expressed gene clusters (370 axolotl genes); this direct comparison is thus considerably less informative than comparisons that include the brittle star. Most genes with conserved expression patterns in the brittle star/axolotl comparison lack identifiable homologs in Parhyale, whereas genes with a conserved expression in the brittle star/Parhyale comparison exhibit a different expression pattern in the axolotl (**Figure 5C**). This underscores the relevance of using the brittle star to bridge comparisons across established regeneration models.

The broadly-conserved co-expression clusters largely consist of genes expressed during the proliferative phase, and, to a lesser extent, the initial wound healing phase. In contrast, the genes that comprise clusters corresponding to tissue differentiation are distinct in each species, which is consistent with the fact that the regenerating appendages are not homologous across species. Strikingly, the conserved co-expression clusters are deployed in a consistent temporal sequence in each species (**Figure 5C**). Specifically, the only identified heterochrony concerns the matching of the axolotl cluster Ax3 (peak at 0-3 hpa) with brittle star cluster A5 (peak at 6 dpa, **Figure 5C**, **Figure 4A, Figure S8**). Previous work suggested that similar co-expression gene modules are deployed during regeneration and development, but are activated according to distinct temporal sequences (Sinigaglia et al. 2022). To investigate if this extends across species, we compared gene expression profiles during regeneration and development from the brittle star and Parhyale, notwithstanding the fact that the indirect development of brittle stars does not allow a direct comparison (**Figure S9**). Results indicate that the order in which co-expressed gene modules are activated is as expected more conserved within regeneration and within developmental datasets across species than between regeneration and development in individual species (**Figure S9**). Together, these results broaden previous observations of distinct expression dynamics during development and regeneration, and document conserved gene expression modules recruited for animal appendage regeneration (**Table S6**).

We further investigated the functions of brittle star genes with similar temporal expression profiles during regeneration in Parhyale and/or axolotl. Using a carefully selected background that accounts for homology-detection and functional biases of different clusters (**Methods**), we found that, within the set of genes that exhibit conserved expression in regeneration, there is a significant over-representation of kinase and stemness genes and an under-representation of immune genes (Gene lists enrichment tests, **Figure 4D, Figure S9**). Moreover, these genes are enriched in general biological processes related to cell proliferation, such as translation, chromosome segregation, DNA replication and intracellular transport (Gene Ontology enrichment tests, **Figure 4E**). The temporal expression patterns of genes that encode transcription factors are neither significantly more nor less conserved across species than is expected at random. Among the conservative set of 154 genes with conserved expression profiles across the three species, only two transcription factors emerge (**Table S6**): *Id2-like*, which activates regeneration-induced proliferation in mice (Kiyokawa et al. 2021) and *Wdhd1-like*, which regulates DNA replication (Zhou and Chen 2021). We thus propose that *Id2* and *Wdhd1* may play a conserved role during animal regeneration. In addition, while several TF binding motifs found in the vicinity of brittle star co-expressed genes are also over-represented near Parhyale and axolotl co-expressed genes, only YY1 and NRF1 are present in corresponding co-expression clusters (Ax7-A6, **Figure S8**), suggesting a possible conserved role for these transcription factors in regulating cell proliferation during regeneration in these distantly related organisms.

Finally, we find that two temporally-matched gene expression clusters in brittle star and Parhyale regeneration include key genes involved in repressing transposable elements (i.e. *Risc-like* [A2 - P1] and *Ago2-like* [A9 - P7]; **Table S6**). It has been proposed that transposon repression is important for proceeding from the immune response phase to regeneration (Angileri et al. 2022), by preserving genome integrity for cell proliferation and differentiation. In line with this hypothesis, we found a higher transcriptional activity of brittle star repetitive elements in the initial wound-response regeneration phase compared to the proliferative phase (**Figure S11, Methods**). Repetitive sequence divergence analysis indicates that these repetitive elements are significantly younger and are more frequently present in intergenic regions than those expressed during cell proliferation, suggesting a higher potential to effectively be active transposable elements.

### Explant experiments reveal expression differences between non-regenerative and regenerative responses in the brittle star

To define what differentiates regeneration from non-regenerative wound healing at the molecular level, we performed explant experiments in which the arm is first amputated from the body (proximal cut) and subsequently amputated a second time at the distal end (**Figure 6A**). As in whole animals, explanted brittle star arms regenerate from the distal tip, whereas the proximal end undergoes a non-regenerative wound healing response. To identify genes specifically involved in regeneration, we sampled distal, medial and proximal explant segments for RNA-seq experiments at 3 and 5 days post-amputation (dpa) when morphological differences start to become apparent (3 to 4 replicates each, for a total of 20 samples, **Figure 6A, Table S1**).

**Figure 6:**
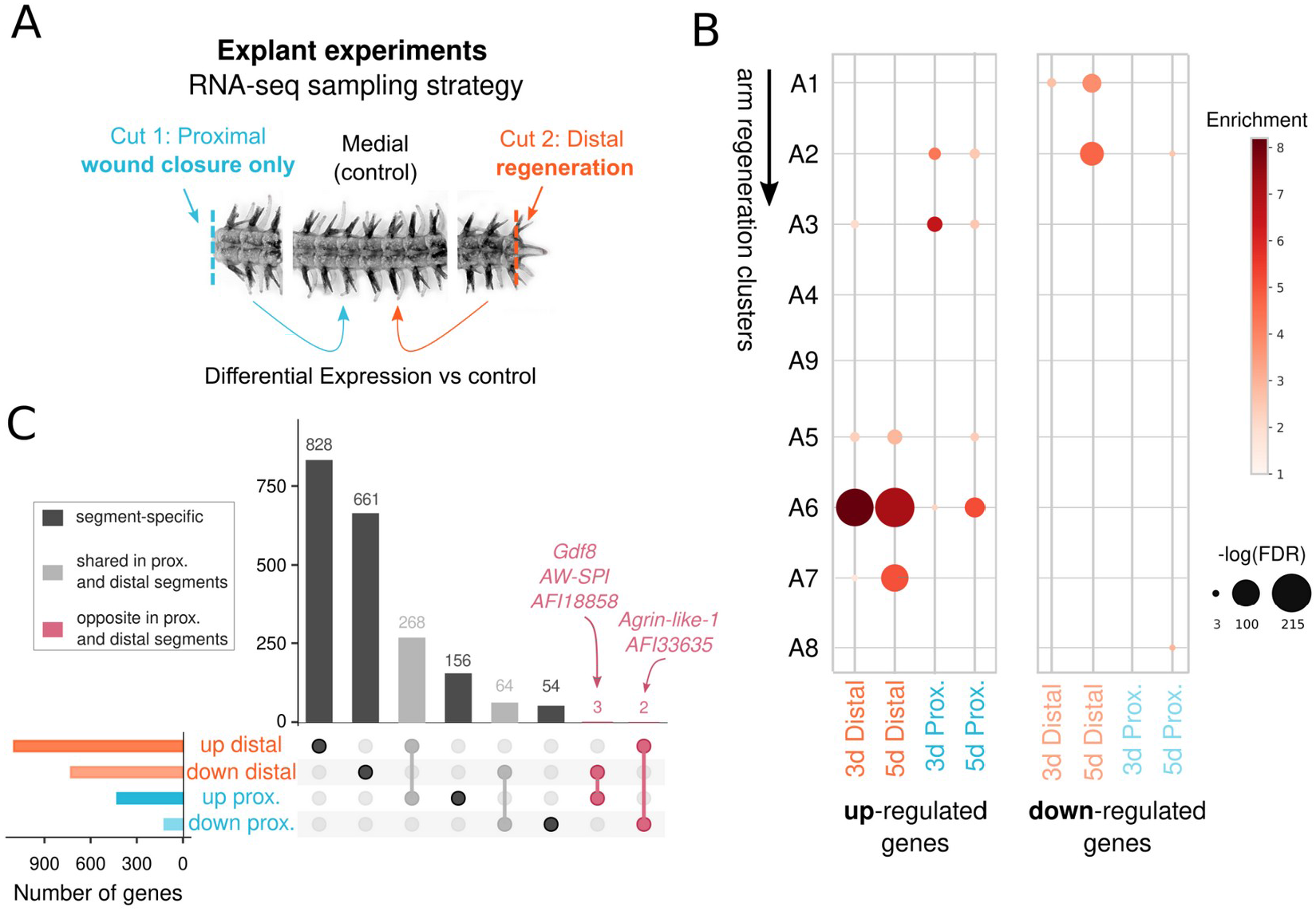
Comparison of gene expression during wound closure and regeneration in brittle star explant experiments. **A.** Experimental setup. Brittle star arms are amputated at the proximal (cut 1) and distal (cut 2) ends. Proximal, distal and medial (control) segments are sampled for RNA-seq at 3 and 5 days post-amputation (dpa). We identify differentially expressed genes (DEGs) in proximal (wound closure only, not followed by regeneration) segments and distal (regenerative) segments, compared to control medial segments. **B.** Comparison of DEGs from explant experiments with brittle star arm regeneration time-course clusters (see Figure 4, hypergeometric enrichment test, BH-corrected p-values <0.05). **C.** Overlap between DEGs genes in distal and proximal segments. Bars in the UpSet are coloured to highlight (i) segment-specific DEGs, for DEGs unique to distal or proximal segments, (ii) shared prox. and distal segments, for DEGs shared between proximal and distal, and (iii) opposite prox. and distal segments, for DEGs up-regulated in proximal and down-regulated in distal (or vice-versa).

We tested for differential expression of genes at the distal and proximal end compared to control medial segments (**Methods**, **Figure 6A**). As expected, up-regulated distal genes correspond to genes expressed during the proliferative phase of the brittle star arm regeneration time-series, whereas up-regulated genes in proximal segments correspond to early response/wound closure genes (**Figure 6B**). We identified more differentially expressed genes (DEGs) in the distal regenerating samples than in proximal non-regenerating samples (distal: 595 and 828 up-regulated genes at 3 and 5 dpa respectively, 238 and 562 down; proximal: 148 and 373 up, 27 and 97 down; **Figure 6C**). Most genes differentially expressed in proximal segments are also differentially expressed in distal segments (61% of the proximal DEGs are shared with distal), whereas distal genes are largely distal-specific (82% of the distal DEGs are not shared with proximal) (**Figure 6C**). This is consistent with the expected expression patterns, as wound closure is an integral part of regeneration. Altogether, we identify hundreds of differentially expressed candidate genes, which document the genetic commonalities and differences between non-regenerative wound closure and regeneration (**Table S2**).

Strikingly, five genes display drastically opposite expression patterns in the wound healing and regenerating segments (**Figure 6C**) and thus are likely to contribute to distinct post-wounding outcomes. *Agrin-like-1* and *AFI33635* are significantly down-regulated during wound healing but up-regulated in regeneration (**Figure 6C**). Agrin proteins are critical for neuromuscular junction development in vertebrate embryogenesis (Hoch 1999). *AFI33635* is an uncharacterized brittle star gene with thyroglobulin and methyltransferase domains, putatively involved in regulating protease activity (Novinec et al. 2006). Conversely, the three genes *AW-SPI, AFI18858 and Gdf8* are significantly up-regulated during wound healing but down-regulated in regeneration (**Figure 6C**). *AW-SPI* is an antistasin/WAP-like serine protease inhibitor, with a possible role in immune defence (Yan et al. 2016). *AFI18858* is a brittle star gene with a zf-Bbox domain, and is a member of the expanded TRIM-like gene family, broadly involved in immune responses (**Figure 3C**). Interestingly, the myostatin gene *Gdf8* is a member of the TGF-beta signalling pathway that inhibits skeletal muscle growth and regeneration in mice (McCroskery et al. 2005; Elkasrawy and Hamrick 2010). Our findings suggest that repression of *Gdf8* may similarly enable muscle regeneration in brittle stars. In summary, these five candidate genes might be tightly linked with the transition from wound healing to regeneration-induced cell proliferation, and some may have a conserved function in the brittle star and vertebrates (*Agrin* and *Gdf8*).

## Discussion

The chromosome-scale genome of the brittle star *Amphiura filiformis* represents a critical resource for the fields of evolutionary genomics, marine ecology and regenerative biology. We leveraged this novel genome to gain fundamental insights into echinoderm chromosome evolution, Hox cluster organisation and regenerative processes across animals. Analyses of chromosome evolution have been key to understanding the basic principles of genome evolution, particularly those linked to the emergence of metazoan and bilaterian clades (Simakov et al. 2020; Simakov et al. 2022). Whereas previous studies of chromosome evolution in echinoderms were limited to sea urchins (Simakov et al. 2020; Marlétaz et al. 2023), our analyses revealed that the genomes of sea cucumbers and sea stars display even fewer rearrangements of the bilaterian ancestral chromosomal units than sea urchins. Interestingly, sea cucumbers have the lowest rate of inter-chromosomal rearrangements, yet the most derived echinoderm body plan (Rahman et al. 2019), which highlights the uncoupling of global genomic rearrangements from morphological evolution. We showed that the ‘Eleutherozoa Linkage Groups’ descend from a single fusion of ancestral bilaterian linkages (B2+C2). Chromosome-scale crinoid and hemichordate genomes will reveal whether this fusion is ancestral to Ambulacraria (the clade encompassing echinoderms and hemichordates). Critically, the fusion has not been observed in the genome of *Xenoturbella bocki*, a member of the Xenacoelomorpha lineage whose phylogenetic position is controversial, and thus cannot be used to support their proposed grouping with Ambulacraria (Philippe et al. 2019; Schiffer et al. 2022). In contrast with its sea star sister-group, the *A. filiformis* genome is highly rearranged: our analyses identified 26 inter-chromosomal rearrangements since the Eleutherozoa ancestor. A more comprehensive sampling of brittle star genomes will provide additional context toward establishing the precise timeline of chromosomal rearrangements and investigate the relative contributions of repeat expansion, chromatin architecture and population genetics dynamics to the rapid karyotype evolution in this group.

On a more local scale, we identified convergent rearrangements in the Hox clusters of sea urchins and the brittle star, which could be hallmarks of relaxed regulatory constraints within echinoderms, perhaps resulting from the temporal decoupling of embryo and adult patterning. Specifically, Hox genes, and in particular anterior Hox, show limited expression during echinoderm embryogenesis and are mostly expressed in adults, suggesting that they are mostly used to pattern adult structures (Arenas-Mena et al. 1998; Arenas-Mena et al. 2000; Hara et al. 2006; Kikuchi et al. 2015; Li et al. 2018). In this context, we speculate that anterior and central/posterior Hox genes may belong to distinct chromatin compartments in echinoderms. Small-scale rearrangements may have occurred through elevated physical contacts at compartments boundaries (i.e. around *Hox4*), and eventually become fixed due to relaxed selection constraints on Hox expression. Further study of chromatin conformation and regulatory footprint in rearranged and non-rearranged echinoderms will make it possible to test this hypothesis. In addition, we revealed significant expansions of transposable elements in the brittle star Hox cluster ∼100 Mya. In the event that Hox cluster rearrangements co-occurred with the activation of repeats, distantly related brittle star species (O’Hara et al. 2014) may exhibit distinct Hox organisations.

The brittle star genome furthermore enables genetic characterization of the animal appendage regeneration process, and remarkably allows the detection of long-range conservation of the gene expression programme that regulates regeneration. In particular, incorporating the brittle star within a comparative transcriptomics framework extensively increased our ability to detect conserved co-expression modules between vertebrates (e.g. axolotl) and arthropods (e.g. Parhyale). We revealed that the proliferative phase of regeneration displays the highest expression conservation across these animals, suggesting that regeneration deploys an ancient, evolutionarily conserved proliferation machinery. This observation ties in with the proposition that animal regeneration may recruit a homologous proliferating cell type (Lai and Aboobaker 2018; Srivastava 2021), a hypothesis that should be further explored with single-cell sequencing techniques and additional comparative analyses. The stronger conservation of gene expression during proliferation as opposed to the initial wound healing response is moreover consistent with the elevated turnover of immunity-related genes, broadly reported across animal lineages (Nei et al. 1997; Leulier and Lemaitre 2008; Saco et al. 2023) and which we also demonstrate here in echinoderms. Our results however contrast with the only previous study to have explicitly interrogated the conservation of animal regeneration gene expression programmes, which revealed a higher conservation of early response genes as opposed to the genes expressed during proliferation (Cary et al. 2019). These discrepancies might be due to asynchronous temporal sampling across species in the comparisons of (Cary et al. 2019), which is alleviated in our study through more comprehensive samplings of regeneration time points and explicit comparisons of temporal expression profiles. Alternatively, they could reflect genuine biological differences of (larval) whole body regeneration studied in (Cary et al. 2019) and the adult appendage regeneration we investigate here.

Finally, in the brittle star *A. filiformis*, we uncover significant expansions of gene families linked to regeneration-related processes and in particular of homologs of vertebrate coagulation regulator genes, suggesting them as relevant candidates for follow-up in-depth functional characterizations. We also propose a conserved role for *Gdf8* during regeneration, as it is repressed during regenerative proliferation in both brittle stars and mice (McCroskery et al. 2005; Elkasrawy and Hamrick 2010). Our findings emphasise the importance of echinoderms as a powerful model for regeneration owing to their unique regenerative capabilities and experimental amenability, but also to their phylogenetic position crucial for comparative analyses. The extensive genomic and transcriptomic resources we generated for the brittle star *Amphiura filiformis* thus represent a cornerstone to understand the evolutionary, molecular and genetic underpinnings of animal appendage regeneration, emergence of pentameral symmetry and remarkable diversity of morphologies and developmental strategies seen across echinoderm lineages.

## Data and code availability

Genome sequence and RNA-seq data have been deposited in NCBI SRA (Bioproject PRJNA1029566) and will be made publicly available upon publication. The code for the annotation workflow is publicly available on GitHub (https://github.com/eparey/AnnotateSnakeMake). Supplemental datasets to reproduce the results have been deposited in Zenodo (https://zenodo.org/doi/10.5281/zenodo.10036671, see Supplemental Material). These include the genome, gene and repeat annotations, processed gene expression tables and source data for the figures.

## Author contributions

OOM, TL, MT, SD and KJ initiated the genome sequencing project. OOM, PO, FM and EP designed the study. FM, PO generated sequencing data, and FM and EP assembled and annotated the final genome version. LP performed proximity ligation experiments. For transcriptomic analyses, OOM and TL designed the explant study and generated RNA-seq explant data, and AC collected regeneration time course samples. EP performed the computational analysis of synteny, gene family and comparative transcriptomics of regeneration with the support of FM. DD and SA contributed in data analysis and visualisation. OOM, JD, KMB, PM and PO curated the data and assisted with interpreting the results. EP wrote the manuscript with key contributions from FM, PM and KMB. All authors commented on the manuscript and approved the final version.

## Acknowledgements

This work was supported by the Centre for Marine Evolutionary Biology at the University of Gothenburg (http://www.cemeb.science.gu.se/) and the IMAGO project led by Anders Blomberg. EP is supported by a Newton International Fellowship from the Royal Society (NIF\R1\222125). FM is supported by a Royal Society University Research Fellowship (URF\ R1\191161), BBSRC (BB/V01109X/1), Leverhulme (RPG-2021-436) and a Japan Society for the Promotion of Science Kakenhi grant (JP 19K06620). OOM was financially supported by CEMEB through grants from Swedish research councils Formas and VR (217-2008-1719) and from a VR grant to KJ (253016979). JD is supported by an F.R.S.-FNRS research project (PDR, 40013965), previously held an F.R.S.-FNRS ‘Chargé de recherche’ fellowship (CR, 34761044), and also received financial support from an F.R.S.-FNRS research project (PDR, T.0169.20) and the Biosciences Research Institute of the University of Mons. PM visited the Department of Genetics, Evolution & Environment of UCL financed by a short-term scientific mission Grant of the EU COST Action MARISTEM (CA-16203). KMB is supported by the National Science Foundation (NSF Award 2131297). PO visits to the Kristineberg Marine Station (Sweden) were supported by the KVA fund SL2015-0048 from the Royal Swedish Academy of Sciences and the EU FP7 Research Infrastructures Initiative ASSEMBLE (227799). PO is supported by the BBSRC (BB/W017865/1).

We thank the staff at the Kristineberg Center for Marine Research and Innovation, especially Ursula Schwarz, for assistance during animal collection. We thank Albert Sabarí i Martí, Emre Onal, Larissa Henke and Wendy Hart for their assistance in conducting experiments. We acknowledge the Okinawa Institute of Science & Technology sequencing facility and its members for their support, especially Nana Arakaki and Mayumi Kawamitsu. Finally, we thank Jonathan Rast for helpful comments on the manuscript.

The authors would like to dedicate this manuscript to Michael Thorndyke and R. Andrew Cameron, who were both instrumental in the early stages of this project but, sadly, passed away before analyses were completed. Without their contributions, this work would not have been possible.

## Methods

### Animal sampling

Adult *A. filiformis* were collected at 25-40 m depth from sediment in the Güllmarsfjord in the vicinity of Kristineberg Marine Station, Sweden, using a Petersen mud grab. Individuals were separated from the sediment by rinsing them with seawater and then maintained in natural flowing seawater at 14°C. Sperm was collected from a single-individual by dissecting the gonads from the bursae.

### DNA extraction and sequencing

Sperm cells were concentrated by centrifugation, washed repeatedly, and subsequently embedded in 2% low melting agarose. Sperm cells were lysed in a solution of 1% SDS, 10mM Tris (pH 8) and 100mM EDTA and then resuspended in a solution of 0.2% N-laurylsarcosine, 2mM Tris (pH 9) and 0.13 mM EDTA. High molecular weight (HMW) DNA was released from the agarose blocks using β-agarase (NEB).

Long-read sequencing was performed on six Nanopore PromethIon flowcells (vR9.4.1). Several libraries were constructed using the Ligation Sequencing Kit (Nanopore LSK109) using DNA sheared to different size using a megaRuptor (Diagenode) to optimise yield and contiguity. Bases were called from raw signal with Guppy (model “dna_r9.4.1_450bps_hac_prom”, version 2.3.5). A total of 160.56 Gb nanopore reads was acquired (∼100x coverage). A library of 10x linked-reads was generated using the Chromium system (10x genomics) and sequencing on a Novaseq6000 SP lane in a 2 x 150 bp layout for a total of 246M reads (86 Gb). Genome size was estimated to 1.33 Gb with a heterozygosity of 3.22% by counting k-mer (k=31) in the short read data using jellyfish2 (Marçais and Kingsford 2011) and fitted through a four-peak model using Genomescope2 (Ranallo-Benavidez et al. 2020).

### Genome Assembly

We assembled Nanopore reads using flye (v2.9-b1768) (Kolmogorov et al. 2019) assuming a coverage of 30x and a genome size of 3 Gb to account for the high level of heterozygosity. We obtained a diploid assembly of 2.86 Gb (N50: ∼2.78 Mb) which was subsequently polished using Racon (v1.5.0) (Vaser et al. 2017) for two iterative rounds using the nanopore reads and then for another two rounds using the short-read illumina reads that were aligned to the assembly using minimap2 (v2.24-r1122) (Li 2018). Haplotypes were then removed from the assembly using purge_dups (v1.2.5) (Guan et al. 2020) with cutoffs visually adjusted from the coverage distribution on contigs. The resulting assembly had a total length of 1.57 Gb, with N50 and L50 of 3.2 Mb and 154 respectively, and 96.1% complete BUSCO score.

To scaffold this assembly, we built a Hi-C library using the Omni-C kit (Dovetail) from gonadal tissue. Chromatin was fixed using PFA and digested using a sequence-independent nuclease after re-ligation and biotinylation. A sequencing library was built from purified DNA and 225M reads sequenced on a Novaseq X (∼45x coverage). Hi-C reads were mapped to the polished haplopurged assembly using bwa mem (0.7.17-r1198-dirty) with options -5SP -T0 and alignment were further sanitised, sorted and duplications removed using pairtools (v1.0.2) (Open2C et al. 2023) with options ‘--walks-policy 5uniquè, ‘--max-inter-align-gap 30’ an a minimum MAPQ of 40. We used YAHS (v1.1a-r3) (Zhou et al. 2023) to scaffold the genomic contigs using the Hi-C read alignment as input. We obtained 20 main chromosome-scale scaffolds totalling 1.47 Gb corresponding to 93.5% of the total assembly length (**Figure S1**). The GC level of the final genomic sequence is 36.67 % and the N50 is 68.86 Mb.

### Repeat annotation

We used RepeatModeler 2.0.2 to build a *de novo* repeat library for the brittle star genome and then ran RepeatMasker 4.1.2-p using this library as input to soft-mask the genome and extract repeat location (Flynn et al. 2020). We used DeepTE (Yan et al. 2020) to classify repeats that could not be classified with the homology-based repeat classification implemented in RepeatModeler. We re-trained a DeepTE model to classify metazoan repeats into 5 classes, using a balanced dataset of 12,500 distinct repeats (2,500 repeats for each of the 5 classes) from different sources including repbase (Bao et al. 2015), Dfam (Hubley et al. 2016) and homology-based classifications of repeats from 17 echinoderms and 2 hemichordates genomes (Validation accuracy=0.98 at the class probability threshold p>=0.55, **Figure S1 B**). On a test set of 827 brittle star repeat families that were not included in the training set and where RepeatModeler homology-based predictions serve as ground truth, this re-trained DeepTE model has higher accuracy than the default metazoa model available in DeepTE (accuracy=0.81 vs 0.67, **Figure S1 C**). Divergence to consensus (kimura %) were computed and repeat landscapes plotted using the ‘calcDivergence.pl’ and ‘createRepeatLansdscape.pl’ scripts from RepeatMasker. The same methodology was applied to build repeat landscapes for *P. lividus*, *H. leucospilota* and *M. glacialis*. Repeat annotations are provided in **Dataset S1**. Repeat ages were estimated from divergence to consensus using a neutral substitution rate of 1.885 x 10^-9^ per base pair per year for *A. filiformis*, which was estimated with phyloFit from an alignment of 66,818 4-fold degenerate sites containing 17 echinoderm and 2 hemichordate genomes.

### RNA isolation, extraction and sequencing

#### Arm regeneration RNA-seq in brittle star (time course in whole animals)

*A. filiformis* individuals were obtained in the fjord close to the Kristineberg Center for Marine Research and Innovation, Sweden, at depths of 20-60 metres. Samples of different regenerating stages were obtained as previously described in (Czarkwiani et al. 2016) for early regeneration stages (48 hpa, 72 hpa, Stages 3, 4 and 5) and in (Dupont and Thorndyke 2006) for 50% differentiation index stages (50% P and 50% D). 30 regenerates from different individuals were used per stage. Dissection for RNA-sampling was performed as follows (**Figure 4A**): (i) for the non-regenerating control, we dissected one mature arm segment, (ii) for 48 and 72 hpa samples, we dissected the last segment at the amputation site, (iii) for stages 3 to 5 we dissected the regenerative tissues, and finally (iv), for 50% regenerates, we sampled several segments of proximal and distal tissues, excluding the differentiated distal cap structure. The collected regenerates were lysed in 10 volumes of RLT (Qiagen), and total RNA extracted using RNAeasy micro RNA kit (Qiagen). RNA concentration and integrity was measured using Bioanalyzer (Agilent). Library preparation and paired-end sequencing was conducted by Novogene.

#### Arm regeneration RNA-seq in brittle star severed arm experiments (explant)

We collected around 3,500 brittle stars with a 5-7 mm disc diameter. While animals were sedated in 3.5% w/w MgCl_2_ in artificial seawater, two arms from each organism were amputated by pressing a scalpel blade into the intervertebral autotomy plane. We first sectioned the arms 0.5 cm from the disc (amputation 1, **Figure 6A**) and then sectioned them again at the distal end (amputation 2, **Figure 6A**). We thus produce explants (i.e. severed brittle star arms) of 1cm in length with wound sites at the proximal and distal ends. 43 groups of 150-200 explants were cultured in flow through aquaria at 16°C. Explants were sampled at 3 and 5 days post-amputation (dpa), sedated in 3.5% w/w MgCl_2_ in artificial seawater for 15 minutes and then dissected into three sections: proximal, medial and distal (**Figure 6A**). Each explant section was flash frozen in liquid nitrogen and collected in batches of 150-200 pieces. Each batch was individually homogenised with glass pistils and RNA was extracted with the RiboPure kit (Applied Biosystems), following manufacturer’s protocol. RNA concentrations were measured using a QuBit 2.0 RNA fluorometric assay (Thermo Fisher Scientific, Waltham, MA, USA) and RNA integrity was checked using 0.5% (w/v) agarose-MOPS-formaldehyde de-naturating gel electrophoresis.

Complementary DNA (cDNA) libraries were prepared using the Illumina TruSeq v2 mRNA sample prep kit (Illumina, San Diego, CA, USA), following a standard protocol. Briefly, mRNA was isolated with poly-A selection, followed by cDNA synthesis, Illumina standard index adapter ligation and a brief PCR reaction. Concentrations of the cDNA libraries were measured using a QuBit DNA High-sensitivity assay (Thermo Fisher Scientific, Waltham, MA, USA) and fragment length distributions were assessed using an Agilent TapeStation with a D1000 tape (Agilent, Santa Clara, CA, USA). cDNA libraries were multiplexed by equimolar pooling (5 or 6 samples/pool), and were then sent to the Swedish National Genomics Infrastructure’s SNP & SEQ platform in Uppsala for Illumina HiSeq 2500 sequencing (8 lanes; 126 bp Paired-End sequencing; Illumina, San Diego, CA, USA).

### Gene annotation

We annotated the brittle star genome using a custom pipeline leveraging three types of evidence: (i) assembled transcriptomes from 18 brittle star developmental and arm regeneration RNA-seq (including both publicly available (Delroisse et al. 2014; Purushothaman et al. 2015; Dylus et al. 2016) and newly generated datasets, **Table S1**), (ii) similarity to proteins from 27 selected metazoa and (iii) *ab initio* predictions. We first assembled a consensus transcriptome from all RNA-seq samples with mikado (Venturini et al. 2018), combining an alignment-free transcriptome assembled with Trinity (Grabherr et al. 2011) and mapped to the genome with gmap (Wu and Watanabe 2005) with an alignment-based transcriptome mapped to the genome with STAR (Dobin et al. 2013), assembled with Stringtie (Pertea et al. 2015) and merged with taco (Niknafs et al. 2017). Second, we selected best-scoring mikado transcripts (i.e. transcripts with identified start and stop codons by TransDecoder, at least 2 exons, over 50% of the predicted coding sequence covered by a swissprot (Boutet et al. 2007) blast hit and no overlap of the coding sequence with an annotated repeat) to train a gene prediction model with AUGUSTUS (Stanke and Waack 2003). Third, we obtained similarity-based gene predictions with Metaeuk, based on proteomes from a total of 27 metazoa, including 8 echinoderms and 2 hemichordates. Fourth, we performed *ab initio* gene prediction with AUGUSTUS (Stanke and Waack 2003), using the previously trained model and providing as hints the predicted exons by mikado and Metaeuk, and curated splice junctions defined by portcullis (Mapleson et al. 2018) on the STAR transcriptome. Fifth, we filtered out all predicted gene models with over 40% of exons overlapping annotated repeats and then ran PASA (Haas et al. 2003) on retained genes to finalise models and annotate UTRs. Finally, we further filtered out 3,465 poorly supported gene models (no PFAM domain, no swissprot blast hit and maximal expression < 2 TPM), to retain 30,267 annotated gene models in the final annotation (**Table S2**). The quality and completeness of the annotation is demonstrated by a score of 92.7% complete BUSCO (C:92.7 [S:86.2%, D:6.5%], F:5.0%, M:2.3%, n:954) (Simão et al. 2015) and a total of 4,974 unique PFAM domains (Finn et al. 2014), with 76% of genes (23,047) containing a PFAM domain. Genes were named by BLAST search against the swissprot database. The names of genes specifically investigated in this study (*hox*, *phb*, *luciferase)* were further manually curated to reflect their evolutionary history. This genome annotation pipeline is implemented as a Snakemake workflow (Köster and Rahmann 2012) and is publicly available on Github (https://github.com/eparey/AnnotateSnakeMake). Annotation files are provided in **Dataset S1**.

### Synteny comparisons and reconstruction of Eleutherozoa ancestral linkage groups

For the sea urchin *P. lividus* and the black sea cucumber *H. leucospilota*, we used previously reported gene annotations (Chen et al. 2023; Marlétaz et al. 2023). We generated a draft homology-based annotation for the spiny sea star *M. glacialis* (Lawniczak et al. 2021) with MetaEuk (Levy Karin et al. 2020) using proteins of the sea urchin *S. purpuratus* (Spur_5.0, available in Ensembl Metazoa v56, (Sea Urchin Genome Sequencing Consortium et al. 2006)), the crown-of-thorns sea star *A. planci* (OKI_Apl_1.0, available in Ensembl Metazoa v56 (Baughman et al. 2014)) and the octopus sea star *P. borealis* (Lee et al. 2022). One-to-one orthologous genes were identified by reciprocal best blast hit between pairs of compared genomes, using diamond (Buchfink et al. 2021). We used Circos version 0.69.8 and circos-tools 0.23 (Krzywinski et al. 2009) to plot synteny comparisons, with the bundlelinks tool to group together neighbouring genes (with a maximum gap of 50 genes) and filter out bundles with fewer than 3 links. We then used the orderchr tool to order chromosomes so as to minimise link crossings. The ancestral Eleutherozoa linkage groups were reconstructed on the basis of synteny comparisons between the spiny sea star *M. glacialis* and the black sea cucumber *H. leucospilota*, and with the amphioxus *B. floridae* and the scallop *P. maximus* genomes as well as previously defined bilaterian linkage groups (BLGs) to untangle derived from ancestral chromosomal arrangements (**Figure S2**). Specifically, only one macro-syntenic rearrangement occurred between the spiny sea star and the black sea cucumber: (a) spiny sea star chr5 maps to both sea cucumber chr12 and chr23. Comparisons with outgroups and ancestral BLGs revealed that (a) corresponds to a derived fusion in the spiny sea star and that the black sea cucumber retained the ancestral state. Using this reconstruction, we annotated genes from matched orthologous chromosomes between sea stars and sea cucumbers with respect to their ancestral ELGs of origins and propagated annotations to orthologous genes in *P. lividus*, *A. filiformis* and other available chromosome-scale echinoderm genomes. Karyotypes were drawn with RIdeograms (Hao et al. 2020): we painted genes on extant chromosomes using the ancestral chromosome colour when a significant number of genes were inferred to descend from an ancestral chromosome (p<10^-5^, Fisher exact tests corrected for multiple testing with the Benjamini-Hochberg procedure). Oxford grid plots of ELGs distribution in *P. lividus*, *A. filiformis and other s*equenced echinoderms (**Figure S3**) were plotted using the same statistical thresholds. ELG-related data files are provided in **Dataset S2**.

### Hox and ParaHox genes identification

We first compiled a dataset of curated full length HOX protein sequences from *S. purpuratus* and HOX homeodomains from *B. floridae* and *S. kowaleskii* to search for homologous Hox genes in the brittle star. A comprehensive list of candidate Hox genes in brittle star was then constructed using two approaches: (i) a diamond blastp (Buchfink et al. 2021) of the curated Hox dataset against brittle star predicted proteins (ii) a miniprot (Li 2023) alignment of *S. purpuratus* HOX protein sequence against the brittle star genome sequence, to recover Hox genes potentially missed by the automatic annotation process. The same approach was used to identify Hox genes in *M. glacialis*. Finally, we built a molecular phylogenetic tree (**Figure S4**) with RAxML-NG (Kozlov et al. 2019) using the LG+G4+F model and 5 distinct starting parsimony trees, to reconstruct the evolutionary history of Hox genes from *S. purpuratus*, *B. floridae*, *S. kowaleskii, A. filiformis, M. glacialis* and three additional echinoderms with curated Hox genes in Echinobase (Arshinoff et al. 2022) *(L. variegatus, P. miniata and A. planci).* We extracted ParaHox sequences and location in *A. japonica* from Ensembl Metazoa v56 (Yates et al. 2022), in *S. purpuratus* from Echinobase, *A. planci* from Ensembl Metazoa and similar approaches as for the Hox to identify ParaHox genes in *A. filiformis* and *M. glacialis*. Hox and ParaHox data files are provided in **Dataset S3**.

### Gene families expansion and contraction

We used broccoli (Derelle et al. 2020) to group proteins of 28 selected Metazoa, 10 of which were Ambulacraria, into gene families. Gene families were predicted to have originated in the last common ancestor of the species with a gene in the family. Out of the complete set of broccoli gene families, 10,367 originated before the last common ancestor of Ambulacraria (echinoderms and hemichordates outgroups). We used CAFE5 (Mendes et al. 2020) on the set of 10,367 families to identify significantly expanded and contracted gene families on each branch of the Ambulacraria phylogeny. Briefly, CAFE fits a birth-death model on a dated phylogeny from the gene count data, and tests for significant expansions/contractions on specific branches. To obtain a dated Ambulacraria phylogeny, we: (i) extracted 192 1-to-1 orthologs in Ambulacraria from broccoli gene families, (ii) built multiple sequence alignments for each orthologous group using MAFFT v7.475, (iii) concatenated alignments across orthology groups and reconstructed a Maximum Likelihood phylogeny with RAxML-NG v.1.1 (LG+G4+F model with 10 parsimony starting trees), (iv) filtered out columns of the alignment with over 15% gaps (47,520 retained sites) and (v) ran PhyloBayes v4.1b (Lartillot et al. 2009) to obtain a time-calibrated tree, with the RAxML reconstructed tree as constrained topology and selected fossil calibrations extracted from the literature ((Benton et al. 2009; Mongiardino Koch et al. 2022). The chain was run for 4,166 samples and 3,500 were retained after burn-in to estimate the posterior distributions for node ages. We next ran CAFE in 2 steps: we estimated the lambda and alpha parameters of the 2-categories CAFE GAMMA model excluding the 128 gene families with the largest copy number differential and then ran CAFE on all families with these parameters fixed to test for significant contractions and expansions (p-values <0.05). Fossil calibrations, dated species tree, gene families and CAFE output files are provided in **Dataset S4.**

### Gene lists curation

We generated lists of immune genes in *A. filiformis* (**Table S2**) using a combination of PFAM domain annotation and lists of previously curated immune genes in the sea urchin *Strongylocentrotus purpuratus* (Sea Urchin Genome Sequencing Consortium et al. 2006). Specifically, we first selected *A. filiformis* genes based on their PFAM domains (e.g. TIR, IL17, Mif) and completed this list by searching for homologs (using the set of broccoli gene families) with the immune genes of the sea urchin *Strongylocentrotus purpuratus*. For the list of kinase genes, we also identified through homologies with curated kinase genes from the sea urchin genome (Sea Urchin Genome Sequencing Consortium et al. 2006). We generated a list of TFs based on the presence of DNA-binding PFAM domains. For the stemness genes, we identified putative homologues of the 180 “stemness-like” genes established by (Alié et al. 2015), that is, genes that are shared between three stem cell populations: poriferan (*Ephydatia fluviatilis*) archeocytes, *Hydra vulgaris* interstitial stem cells, and planarian (*Schmidtea mediterranea*) neoblasts. Specifically, we used the human cognates of all those genes as queries for BLAST searches (McGinnis and Madden 2004) against *A. filiformis* genes. We emphasise that, while these genes are expressed in stem cell populations, they may not all be stemness regulators. For genes involved in neuronal function, we first compiled a list of neurogenic and glial markers, TFs, cell signalling genes involved in embryonic, homeostatic, and regenerative neurogenesis in vertebrates (rodents, humans, and Xenopus) and invertebrates (*Caenorhabditis elegans* and *Drosophila melanogaster*). We identified putative homologues of “neuronal” genes in *A. filiformis* using a reciprocal blast approach. We generated gene lists for 19 signalling pathways, in two steps: (i) manual curation of main members of selected pathways, (ii) identification of their gene ID in the *S. purpuratus* genome via echinobase gene searches (Arshinoff et al. 2022) and (iii) identification of *S. purpuratus* orthologs in *A. filiformis* using gene trees built with Generax (Morel et al. 2020) for each of our broccoli (Derelle et al. 2020) gene family. Finally, the repertoire of luciferase-like genes was identified through reciprocal BLAST searches using the reference *Renilla* luciferase (GenBank: AAA29804) as initial query (Delroisse et al. 2017).

### Gene ontology enrichment tests

We used eggnog-mapper (Cantalapiedra et al. 2021) to automatically annotate *A. filiformis* and *P. lividus* genes with Gene Ontology terms from the Biological Process domain. The GO annotations were then transferred to the level of gene families. Specifically, for each family, we propagated all GO annotations associated with any *P. lividus* or *A. filiformis* genes as the complete set of GO annotations for this family. Hypergeometric tests for functional enrichments were then conducted with the enricher function from the ClusterProfiler R package (Wu et al. 2021), with custom foreground and background GO annotation sets. For functional enrichment tests on expanded/contracted gene families (**Figure 3**), tests were conducted at the level of gene families with expanded or contracted families as foreground and all gene families as background, as described above. For functional enrichment tests on regeneration co-expression clusters (**Figure 4**), tests were conducted at the level of brittle star genes, using genes of a given cluster as foreground and genes of all clusters as background. We used FDR < 0.05 as significance threshold. Enrichment results were summarised with REVIGO (Supek et al. 2011), we selected top ontology terms based on the REVIGO “dispensability” score to make a representative overview of the diversity of enriched GO terms.

### Clustering of the arm regeneration expression series

Gene expression was quantified for all samples using the alignment-free method kallisto (Bray et al. 2016). We normalised TPM values across samples using the TMM method as implemented in edgeR (Robinson et al. 2010; Robinson and Oshlack 2010) and used MFuzz (Kumar and E Futschik 2007) to perform soft-clustering of genes on the basis of their standardised expression profiles across samples. We used the minimum centroid distance method to select the optimal number of clusters (n=19, **Figure S6**). In the main text, we filtered the obtained clusters to retain clusters with > 1 enriched GO term and elevated expression in >1 regenerating sample (**Figure 4, Figure S6**). Normalised gene expression tables are provided in **Dataset S5**.

### Transcription factor binding motif enrichment tests

We used HOMER v4.11 (Heinz et al. 2010), to test for enriched transcription factor binding motifs in the proximal regulatory domains (TSS + 5 kb upstream, +1 kb downstream) of genes of each regeneration cluster. We ran the findMotifsGenome.pl script from the HOMER suite, with –h to perform hypergeometric tests, contrasting proximal regulatory domains of genes from one expression cluster as foreground with proximal regulatory domains from genes of all clusters as background.

### Axolotl limb regeneration RNA-seq time course

Raw RNA-seq data for 12 limb regeneration time points from (Stewart et al. 2013) were downloaded from https://www.axolomics.org/?q=node/2. We used Trim Galore (https://github.com/FelixKrueger/TrimGalore) with default parameters to trim and quality filter raw sequencing reads via the Cutadapt tool (Martin 2011). Gene expression was quantified with kallisto (Bray et al. 2016) using the set of annotated axolotl transcripts from the latest *Ambystoma mexicanum* assembly version (AmexG_v6.0-DD, available from https://www.axolotl-omics.org/assemblies, (Schloissnig et al. 2021)). Similarly as for the brittle star regeneration series, we normalised TPM values across samples using the TMM method (Robinson et al. 2010; Robinson and Oshlack 2010) and used MFuzz (Kumar and E Futschik 2007) to cluster genes according to their expression profile (**Figure S7**). Gene ontology and TFBS motifs enrichment were performed as described for the brittle star. Normalised gene expression tables are provided in **Dataset S5**.

### Parhyale limb regeneration RNA-seq time course

Parhyale leg regeneration expression data were previously processed and clustered into 8 co-expression gene groups using the same approach as we used for brittle star data (Sinigaglia et al. 2022). As such, we directly used the clustering reported in (Sinigaglia et al. 2022), but renamed the clusters so that numbering follow temporal activation (P1 is R4 in the notation of Sinigaglia et al., P2 is R1, P3 is R8, P4 is R2, P5 is R6, P6 is R3, P7 is R5 and P8 is R7).

### Comparison of gene expression dynamics during appendage regeneration

We used broccoli (Derelle et al. 2020) to build homologous gene families encompassing genes of the brittle star *A. filiformis*, the axolotl *Ambystoma mexicanum* and *Parhyale hawaiensis,* as well as 8 echinoderms, 6 vertebrates, 7 ecdysozoans and 12 other animal genomes. We used these gene families to identify homologous genes and compare their expression profile during appendage regeneration in a pairwise manner across the three species. For each pairwise comparison, we retained all homologous gene families with > 1 gene and < 5 genes in each of the two compared species, resulting in a total of 5,203 homologous groups retained for the axolotl (8,810 homologous genes) - brittle star (6,813 homologous genes) comparison, 3,137 for the brittle star (4,196) - Parhyale (3,617) comparison and 2,299 for the axolotl (3,903) - Parhyale (2,628) comparison (see **Dataset S5**). We next computed permutation-based p-values to test for the overrepresentation of homologous genes across co-expression clusters of the two compared species. Specifically, we generated, for each pairwise comparison, 10,000 randomizations of the gene labels of species 2, keeping clusters and orthologous gene family size constant to build a null distribution of the number of expected homologous genes shared by two clusters at random. Empirical p-values were computed from the null distribution and corrected for multiple testing using the Benjamini-Hochberg procedure. To investigate functional annotation of genes displaying co-expression across regeneration models as opposed to genes from the same clusters that do not show co-expression across species, we conducted gene list and GO enrichment tests as described previously but using carefully selected background: we used as background all brittle star genes with a homolog in either Parhyale or axolotl (i.e. whose expression conservation could be tested) and in a cluster with identified co-expressed genes in either Parhyale or axolotl (to test for the specificity of genes of a given cluster that show conservation vs those of the same cluster that do not).

### Differential analysis of repeats transcriptional activity during brittle star arm regeneration

We tested for differentially expressed repetitive elements in early regeneration (immune phase: 48 hpa and 72 hpa samples) versus middle regeneration (proliferation: Stage 3, Stage 4, Stage 5 samples), using our time course brittle star arm regeneration RNA-seq data. We used a conservative approach to first filter out highly duplicated genes which could have been captured in the set of repetitive elements called by RepeatModeler/RepeatMasker. We used diamond blastx (Buchfink et al. 2021) to search for homologies between repeat consensus and proteins in the swissprot database (Boutet et al. 2007) and filtered out all ‘Unknown’ repeat families for which the consensus sequence had a strict match in swissprot (e-value cut-off 10 ^-10^) which did not correspond to transposon genes. We next use the SalmonTE pipeline (Jeong et al. 2018) with default parameters on the full set of filtered repeat consensus (n=4,695 repeat families), followed by differential analysis with DESeq2 (Love et al. 2014) on the estimated count values, to test for differential transcriptional activity of repeats in the immune versus proliferation regeneration phases. We retained as differentially expressed the repetitive elements with an absolute log_2_ fold change > 1, adjusted p-value < 0.001.

### Differential gene expression in brittle star arm explants

Gene expression was quantified for all samples using the alignment-free method Kallisto (Bray et al. 2016). Differential expression analyses were conducted with DESeq2 (Love et al. 2014) on count values, contrasting distal replicates against medial replicates and proximal replicates against medial replicates for each time point. All genes with an adjusted p-value < 0.05 and absolute log_2_ fold change > 1 were retained as differentially expressed. Gene expression tables are provided in **Dataset S5**.

## References

Alié A, Hayashi T, Sugimura I, Manuel M, Sugano W, Mano A, Satoh N, Agata K, Funayama N. 2015. The ancestral gene repertoire of animal stem cells. Proc. Natl. Acad. Sci. U. S. A. [Internet] 112:E7093–E7100. Available from: 10.1073/pnas.1514789112

Angileri KM, Bagia NA, Feschotte C. 2022. Transposon control as a checkpoint for tissue regeneration. Development [Internet] 149. Available from: 10.1242/dev.191957

Annunziata R, Martinez P, Arnone MI. 2013. Intact cluster and chordate-like expression of ParaHox genes in a sea star. BMC Biol. [Internet] 11:68. Available from: 10.1186/1741-7007-11-68

Annunziata R, Perillo M, Andrikou C, Cole AG, Martinez P, Arnone MI. 2014. Pattern and process during sea urchin gut morphogenesis: the regulatory landscape. Genesis [Internet] 52:251–268. Available from: 10.1002/dvg.22738

Apte RS, Chen DS, Ferrara N. 2019. VEGF in Signaling and Disease: Beyond Discovery and Development. Cell [Internet] 176:1248–1264. Available from: https://www.ncbi.nlm.nih.gov/pmc/articles/PMC6410740/

Arenas Gómez CM, Sabin KZ, Echeverri K. 2020. Wound healing across the animal kingdom: Crosstalk between the immune system and the extracellular matrix. Dev. Dyn. [Internet] 249:834– 846. Available from: 10.1002/dvdy.178

Arenas-Mena C, Cameron AR, Davidson EH. 2000. Spatial expression of Hox cluster genes in the ontogeny of a sea urchin. Development [Internet] 127:4631–4643. Available from: 10.1242/dev.127.21.4631

Arenas-Mena C, Martinez P, Cameron RA, Davidson EH. 1998. Expression of the *Hox* gene complex in the indirect development of a sea urchin. Proc. Natl. Acad. Sci. U. S. A. [Internet] 95:13062– 13067. Available from: 10.1073/pnas.95.22.13062

Arnone MI, Rizzo F, Annunciata R, Cameron RA, Peterson KJ, Martínez P. 2006. Genetic organization and embryonic expression of the ParaHox genes in the sea urchin S. purpuratus: insights into the relationship between clustering and colinearity. Dev. Biol. [Internet] 300:63–73. Available from: 10.1016/j.ydbio.2006.07.037

Arshinoff BI, Cary GA, Karimi K, Foley S, Agalakov S, Delgado F, Lotay VS, Ku CJ, Pells TJ, Beatman TR, et al. 2022. Echinobase: leveraging an extant model organism database to build a knowledgebase supporting research on the genomics and biology of echinoderms. Nucleic Acids Res. [Internet] 50:D970–D979. Available from: 10.1093/nar/gkab1005

Ayaz G, Yan H, Malik N, Huang J. 2022. An Updated View of the Roles of p53 in Embryonic Stem Cells. Stem Cells [Internet] 40:883–891. Available from: 10.1093/stmcls/sxac051

Balachandran P, Walawalkar IA, Flores JI, Dayton JN, Audano PA, Beck CR. 2022. Transposable element-mediated rearrangements are prevalent in human genomes. Nat. Commun. [Internet] 13:7115. Available from: 10.1038/s41467-022-34810-8

Bao W, Kojima KK, Kohany O. 2015. Repbase Update, a database of repetitive elements in eukaryotic genomes. Mob. DNA [Internet] 6:11. Available from: 10.1186/s13100-015-0041-9

Barrera-Redondo J, Lotharukpong JS, Drost H-G, Coelho SM. 2023. Uncovering gene-family founder events during major evolutionary transitions in animals, plants and fungi using GenEra. Genome Biol. [Internet] 24:54. Available from: 10.1186/s13059-023-02895-z

Baughman KW, McDougall C, Cummins SF, Hall M, Degnan BM, Satoh N, Shoguchi E. 2014. Genomic organization of Hox and ParaHox clusters in the echinoderm, Acanthaster planci. Genesis [Internet] 52:952–958. Available from: 10.1002/dvg.22840

Bely AE, Nyberg KG. 2010. Evolution of animal regeneration: re-emergence of a field. Trends Ecol. Evol. [Internet] 25:161–170. Available from: 10.1016/j.tree.2009.08.005

Belyayev A. 2014. Bursts of transposable elements as an evolutionary driving force. J. Evol. Biol. [Internet] 27:2573–2584. Available from: 10.1111/jeb.12513

Benton MJ, Donoghue PCJ, Asher RJ. 2009. Calibrating and constraining molecular clocks. In: The timetree of Life. London, England: Oxford University Press. p. 35–86. Available from: https://research-information.bris.ac.uk/en/publications/calibrating-and-constraining-molecular-clocks

Bideau L, Kerner P, Hui J, Vervoort M, Gazave E. 2021. Animal regeneration in the era of transcriptomics. Cell. Mol. Life Sci. [Internet] 78:3941–3956. Available from: 10.1007/s00018-021-03760-7

Boutet E, Lieberherr D, Tognolli M, Schneider M, Bairoch A. 2007. UniProtKB/Swiss-Prot. Methods Mol. Biol. [Internet] 406:89–112. Available from: 10.1007/978-1-59745-535-0_4

Bray NL, Pimentel H, Melsted P, Pachter L. 2016. Near-optimal probabilistic RNA-seq quantification. Nat. Biotechnol. [Internet] 34:525–527. Available from: 10.1038/nbt.3519

Buchfink B, Reuter K, Drost H-G. 2021. Sensitive protein alignments at tree-of-life scale using DIAMOND. Nat. Methods [Internet] 18:366–368. Available from: 10.1038/s41592-021-01101-x

Byrne M, Martinez P, Morris V. 2016. Evolution of a pentameral body plan was not linked to translocation of anterior Hox genes: the echinoderm HOX cluster revisited. Evol. Dev. [Internet] 18:137–143. Available from: 10.1111/ede.12172

Cameron RA, Rowen L, Nesbitt R, Bloom S, Rast JP, Berney K, Arenas-Mena C, Martinez P, Lucas S, Richardson PM, et al. 2006. Unusual gene order and organization of the sea urchin hox cluster. J. Exp. Zool. B Mol. Dev. Evol. [Internet] 306:45–58. Available from: 10.1002/jez.b.21070

Canapa A, Barucca M, Biscotti MA, Forconi M, Olmo E. 2015. Transposons, Genome Size, and Evolutionary Insights in Animals. Cytogenet. Genome Res. [Internet] 147:217–239. Available from: 10.1159/000444429

Cannon JT, Kocot KM, Waits DS, Weese DA, Swalla BJ, Santos SR, Halanych KM. 2014. Phylogenomic resolution of the hemichordate and echinoderm clade. Curr. Biol. [Internet] 24:2827–2832. Available from: 10.1016/j.cub.2014.10.016

Cantalapiedra CP, Hernández-Plaza A, Letunic I, Bork P, Huerta-Cepas J. 2021. eggNOG-mapper v2: Functional Annotation, Orthology Assignments, and Domain Prediction at the Metagenomic Scale. Mol. Biol. Evol. [Internet] 38:5825–5829. Available from: 10.1093/molbev/msab293

Cary GA, Wolff A, Zueva O, Pattinato J, Hinman VF. 2019. Analysis of sea star larval regeneration reveals conserved processes of whole-body regeneration across the metazoa. BMC Biol. [Internet] 17:16. Available from: 10.1186/s12915-019-0633-9

Chen T, Ren C, Wong N-K, Yan A, Sun C, Fan D, Luo P, Jiang X, Zhang L, Ruan Y, et al. 2023. The *Holothuria leucospilota* genome elucidates sacrificial organ expulsion and bioadhesive trap enriched with amyloid-patterned proteins. Proc. Natl. Acad. Sci. U. S. A. [Internet] 120:e2213512120. Available from: 10.1073/pnas.2213512120

Colombera D, Venier G. 1976. Chromosomes of echinoderms. Caryologia [Internet] 29:35–40. Available from: 10.1080/00087114.1976.10796647

Cui M, Atmanli A, Morales MG, Tan W, Chen K, Xiao X, Xu L, Liu N, Bassel-Duby R, Olson EN. 2021. Nrf1 promotes heart regeneration and repair by regulating proteostasis and redox balance. Nat. Commun. [Internet] 12:1–15. Available from: https://www.nature.com/articles/s41467-021-25653-w

Czarkwiani A, Dylus DV, Oliveri P. 2013. Expression of skeletogenic genes during arm regeneration in the brittle star Amphiura filiformis. Gene Expr. Patterns [Internet] 13:464–472. Available from: 10.1016/j.gep.2013.09.002

Czarkwiani A, Ferrario C, Dylus DV, Sugni M, Oliveri P. 2016. Skeletal regeneration in the brittle star Amphiura filiformis. Front. Zool. [Internet] 13:18. Available from: 10.1186/s12983-016-0149-x

Czarkwiani A, Taylor J, Oliveri P. 2022. Neurogenesis during Brittle Star Arm Regeneration Is Characterised by a Conserved Set of Key Developmental Genes. Biology [Internet] 11. Available from: 10.3390/biology11091360

David B, Mooi R. 2014. How Hox genes can shed light on the place of echinoderms among the deuterostomes. Evodevo [Internet] 5:22. Available from: 10.1186/2041-9139-5-22

Davidson PL, Guo H, Swart JS, Massri AJ, Edgar A, Wang L, Berrio A, Devens HR, Koop D, Cisternas P, et al. 2022. Recent reconfiguration of an ancient developmental gene regulatory network in Heliocidaris sea urchins. Nat Ecol Evol [Internet] 6:1907–1920. Available from: 10.1038/s41559-022-01906-9

Davidson PL, Guo H, Wang L, Berrio A, Zhang H, Chang Y, Soborowski AL, McClay DR, Fan G, Wray GA. 2020. Chromosomal-Level Genome Assembly of the Sea Urchin Lytechinus variegatus Substantially Improves Functional Genomic Analyses. Genome Biol. Evol. [Internet] 12:1080– 1086. Available from: 10.1093/gbe/evaa101

Davidson PL, Lessios HA, Wray GA, McMillan WO, Prada C. 2023. Near-Chromosomal-Level Genome Assembly of the Sea Urchin Echinometra lucunter, a Model for Speciation in the Sea. Genome Biol. Evol. [Internet] 15. Available from: 10.1093/gbe/evad093

Delroisse J, Ortega-Martinez O, Dupont S, Mallefet J, Flammang P. 2015. De novo transcriptome of the European brittle star Amphiura filiformis pluteus larvae. Mar. Genomics [Internet] 23:109–121. Available from: 10.1016/j.margen.2015.05.014

Delroisse J, Ullrich-Lüter E, Blaue S, Ortega-Martinez O, Eeckhaut I, Flammang P, Mallefet J. 2017. A puzzling homology: a brittle star using a putative cnidarian-type luciferase for bioluminescence. Open Biol. [Internet] 7. Available from: 10.1098/rsob.160300

Delroisse J, Ullrich-Lüter E, Ortega-Martinez O, Dupont S, Arnone M-I, Mallefet J, Flammang P. 2014. High opsin diversity in a non-visual infaunal brittle star. BMC Genomics [Internet] 15:1035. Available from: 10.1186/1471-2164-15-1035

Derelle R, Philippe H, Colbourne JK. 2020. Broccoli: Combining Phylogenetic and Network Analyses for Orthology Assignment. Mol. Biol. Evol. [Internet] 37:3389–3396. Available from: 10.1093/molbev/msaa159

Dobin A, Davis CA, Schlesinger F, Drenkow J, Zaleski C, Jha S, Batut P, Chaisson M, Gingeras TR. 2013. STAR: ultrafast universal RNA-seq aligner. Bioinformatics [Internet] 29:15–21. Available from: 10.1093/bioinformatics/bts635

Dong X, Guo R, Ji T, Zhang J, Xu J, Li Y, Sheng Y, Wang Y, Fang K, Wen Y, et al. 2022. YY1 safeguard multidimensional epigenetic landscape associated with extended pluripotency. Nucleic Acids Res. [Internet] 50:12019–12038. Available from: 10.1093/nar/gkac230

Duineveld GCA, Van Noort GJ. 1986. Observations on the population dynamics of amphiura filiformis (ophiuroidea: echinodermata) in the southern north sea and its exploitation by the dab, Limanda limanda. Neth. J. Sea Res. [Internet] 20:85–94. Available from: https://www.sciencedirect.com/science/article/pii/0077757986900645

Dupont S, Thorndyke M. 2007. Bridging the regeneration gap: insights from echinoderm models. Nat. Rev. Genet. [Internet] 8:320–320. Available from: https://www.nature.com/articles/nrg1923-c1

Dupont S, Thorndyke MC. 2006. Growth or differentiation? Adaptive regeneration in the brittlestar Amphiura filiformis. J. Exp. Biol. [Internet] 209:3873–3881. Available from: 10.1242/jeb.02445

Dylus DV, Czarkwiani A, Blowes LM, Elphick MR, Oliveri P. 2018. Developmental transcriptomics of the brittle star Amphiura filiformis reveals gene regulatory network rewiring in echinoderm larval skeleton evolution. Genome Biol. [Internet] 19:26. Available from: 10.1186/s13059-018-1402-8

Dylus DV, Czarkwiani A, Stångberg J, Ortega-Martinez O, Dupont S, Oliveri P. 2016. Large-scale gene expression study in the ophiuroid Amphiura filiformis provides insights into evolution of gene regulatory networks. Evodevo [Internet] 7:2. Available from: 10.1186/s13227-015-0039-x

Elkasrawy MN, Hamrick MW. 2010. Myostatin (GDF-8) as a key factor linking muscle mass and bone structure. J. Musculoskelet. Neuronal Interact. [Internet] 10:56–63. Available from: https://www.ncbi.nlm.nih.gov/pubmed/20190380

Ferrario C, Ben Khadra Y, Czarkwiani A, Zakrzewski A, Martinez P, Colombo G, Bonasoro F, Candia Carnevali MD, Oliveri P, Sugni M. 2018. Fundamental aspects of arm repair phase in two echinoderm models. Dev. Biol. [Internet] 433:297–309. Available from: 10.1016/j.ydbio.2017.09.035

Finn RD, Bateman A, Clements J, Coggill P, Eberhardt RY, Eddy SR, Heger A, Hetherington K, Holm L, Mistry J, et al. 2014. Pfam: the protein families database. Nucleic Acids Res. [Internet] 42:D222–D230. Available from: 10.1093/nar/gkt1223

Flynn JM, Hubley R, Goubert C, Rosen J, Clark AG, Feschotte C, Smit AF. 2020. RepeatModeler2 for automated genomic discovery of transposable element families. Proc. Natl. Acad. Sci. U. S. A. [Internet] 117:9451–9457. Available from: 10.1073/pnas.1921046117

Freeman R, Ikuta T, Wu M, Koyanagi R, Kawashima T, Tagawa K, Humphreys T, Fang G-C, Fujiyama A, Saiga H, et al. 2012. Identical genomic organization of two hemichordate hox clusters. Curr. Biol. [Internet] 22:2053–2058. Available from: 10.1016/j.cub.2012.08.052

Fumagalli MR, Zapperi S, La Porta CAM. 2018. Regeneration in distantly related species: common strategies and pathways. NPJ Syst Biol Appl [Internet] 4:5. Available from: 10.1038/s41540-017-0042-z

George CM, Alani E. 2012. Multiple cellular mechanisms prevent chromosomal rearrangements involving repetitive DNA. Crit. Rev. Biochem. Mol. Biol. [Internet] 47:297–313. Available from: 10.3109/10409238.2012.675644

Goldman JA, Poss KD. 2020. Gene regulatory programmes of tissue regeneration. Nat. Rev. Genet. [Internet] 21:511–525. Available from: 10.1038/s41576-020-0239-7

Grabherr MG, Haas BJ, Yassour M, Levin JZ, Thompson DA, Amit I, Adiconis X, Fan L, Raychowdhury R, Zeng Q, et al. 2011. Full-length transcriptome assembly from RNA-Seq data without a reference genome. Nat. Biotechnol. [Internet] 29:644–652. Available from: 10.1038/nbt.1883

Gross JM, Peterson RE, Wu S-Y, McClay DR. 2003. LvTbx2/3: a T-box family transcription factor involved in formation of the oral/aboral axis of the sea urchin embryo. Development [Internet] 130:1989–1999. Available from: 10.1242/dev.00409

Guan D, McCarthy SA, Wood J, Howe K, Wang Y, Durbin R. 2020. Identifying and removing haplotypic duplication in primary genome assemblies. Bioinformatics [Internet] 36:2896–2898. Available from: https://academic.oup.com/bioinformatics/article-pdf/36/9/2896/33180804/btaa025.pdf

Haas BJ, Delcher AL, Mount SM, Wortman JR, Smith RK Jr, Hannick LI, Maiti R, Ronning CM, Rusch DB, Town CD, et al. 2003. Improving the Arabidopsis genome annotation using maximal transcript alignment assemblies. Nucleic Acids Res. [Internet] 31:5654–5666. Available from: 10.1093/nar/gkg770

Hall MR, Kocot KM, Baughman KW, Fernandez-Valverde SL, Gauthier MEA, Hatleberg WL, Krishnan A, McDougall C, Motti CA, Shoguchi E, et al. 2017. The crown-of-thorns starfish genome as a guide for biocontrol of this coral reef pest. Nature [Internet] 544:231–234. Available from: 10.1038/nature22033

Han D, Liu G, Oh Y, Oh S, Yang S, Mandjikian L, Rani N, Almeida MC, Kosik KS, Jang J. 2023. ZBTB12 is a molecular barrier to dedifferentiation in human pluripotent stem cells. Nat. Commun. [Internet] 14:632. Available from: 10.1038/s41467-023-36178-9

Hanington PC, Zhang S-M. 2011. The primary role of fibrinogen-related proteins in invertebrates is defense, not coagulation. J. Innate Immun. [Internet] 3:17–27. Available from: 10.1159/000321882

Hao Z, Lv D, Ge Y, Shi J, Weijers D, Yu G, Chen J. 2020. RIdeogram: drawing SVG graphics to visualize and map genome-wide data on the idiograms. PeerJ Comput Sci [Internet] 6:e251. Available from: 10.7717/peerj-cs.251

Hara Y, Yamaguchi M, Akasaka K, Nakano H, Nonaka M, Amemiya S. 2006. Expression patterns of Hox genes in larvae of the sea lily Metacrinus rotundus. Dev. Genes Evol. [Internet] 216:797– 809. Available from: 10.1007/s00427-006-0108-1

Heinz S, Benner C, Spann N, Bertolino E, Lin YC, Laslo P, Cheng JX, Murre C, Singh H, Glass CK. 2010. Simple combinations of lineage-determining transcription factors prime cis-regulatory elements required for macrophage and B cell identities. Mol. Cell [Internet] 38:576–589. Available from: 10.1016/j.molcel.2010.05.004

Herrera SC, Bach EA. 2019. JAK/STAT signaling in stem cells and regeneration: from Drosophila to vertebrates. Development [Internet] 146. Available from: 10.1242/dev.167643

Hoch W. 1999. Formation of the neuromuscular junction. Agrin and its unusual receptors. Eur. J. Biochem. [Internet] 265:1–10. Available from: 10.1046/j.1432-1327.1999.00765.x

Holland PWH, Marlétaz F, Maeso I, Dunwell TL, Paps J. 2017. New genes from old: asymmetric divergence of gene duplicates and the evolution of development. Philos. Trans. R. Soc. Lond. B Biol. Sci. [Internet] 372. Available from: 10.1098/rstb.2015.0480

Huat TJ, Khan AA, Pati S, Mustafa Z, Abdullah JM, Jaafar H. 2014. IGF-1 enhances cell proliferation and survival during early differentiation of mesenchymal stem cells to neural progenitor-like cells. BMC Neurosci. [Internet] 15:91. Available from: 10.1186/1471-2202-15-91

Hubley R, Finn RD, Clements J, Eddy SR, Jones TA, Bao W, Smit AFA, Wheeler TJ. 2016. The Dfam database of repetitive DNA families. Nucleic Acids Res. [Internet] 44:D81–D89. Available from: 10.1093/nar/gkv1272

Hu MY, Casties I, Stumpp M, Ortega-Martinez O, Dupont S. 2014. Energy metabolism and regeneration are impaired by seawater acidification in the infaunal brittlestar Amphiura filiformis. J. Exp. Biol. [Internet] 217:2411–2421. Available from: 10.1242/jeb.100024

Jeong H-H, Yalamanchili HK, Guo C, Shulman JM, Liu Z. 2018. An ultra-fast and scalable quantification pipeline for transposable elements from next generation sequencing data. Pac. Symp. Biocomput. [Internet] 23:168–179. Available from: https://www.ncbi.nlm.nih.gov/pubmed/29218879

Kawaguchi M, Sugiyama K, Matsubara K, Lin C-Y, Kuraku S, Hashimoto S, Suwa Y, Yong LW, Takino K, Higashida S, et al. 2019. Co-option of the PRDM14-CBFA2T complex from motor neurons to pluripotent cells during vertebrate evolution. Development [Internet] 146. Available from: 10.1242/dev.168633

Kikuchi M, Omori A, Kurokawa D, Akasaka K. 2015. Patterning of anteroposterior body axis displayed in the expression of Hox genes in sea cucumber Apostichopus japonicus. Dev. Genes Evol. [Internet] 225:275–286. Available from: 10.1007/s00427-015-0510-7

Kiyokawa H, Yamaoka A, Matsuoka C, Tokuhara T, Abe T, Morimoto M. 2021. Airway basal stem cells reutilize the embryonic proliferation regulator, Tgfβ-Id2 axis, for tissue regeneration. Dev. Cell [Internet] 56:1917–1929.e9. Available from: 10.1016/j.devcel.2021.05.016

Kolmogorov M, Yuan J, Lin Y, Pevzner PA. 2019. Assembly of long, error-prone reads using repeat graphs. Nat. Biotechnol. [Internet] 37:540–546. Available from: 10.1038/s41587-019-0072-8

Köster J, Rahmann S. 2012. Snakemake--a scalable bioinformatics workflow engine. Bioinformatics [Internet] 28:2520–2522. Available from: 10.1093/bioinformatics/bts480

Kozlov AM, Darriba D, Flouri T, Morel B, Stamatakis A. 2019. RAxML-NG: a fast, scalable and user-friendly tool for maximum likelihood phylogenetic inference. Bioinformatics [Internet] 35:4453– 4455. Available from: 10.1093/bioinformatics/btz305

Krzywinski M, Schein J, Birol I, Connors J, Gascoyne R, Horsman D, Jones SJ, Marra MA. 2009. Circos: an information aesthetic for comparative genomics. Genome Res. [Internet] 19:1639– 1645. Available from: 10.1101/gr.092759.109

Kumar L, E Futschik M. 2007. Mfuzz: a software package for soft clustering of microarray data. Bioinformation [Internet] 2:5–7. Available from: 10.6026/97320630002005

Kumar S, Stecher G, Suleski M, Hedges SB. 2017. TimeTree: A Resource for Timelines, Timetrees, and Divergence Times. Mol. Biol. Evol. [Internet] 34:1812–1819. Available from: 10.1093/molbev/msx116

Lai AG, Aboobaker AA. 2018. EvoRegen in animals: Time to uncover deep conservation or convergence of adult stem cell evolution and regenerative processes. Dev. Biol. [Internet] 433:118–131. Available from: 10.1016/j.ydbio.2017.10.010

Lartillot N, Lepage T, Blanquart S. 2009. PhyloBayes 3: a Bayesian software package for phylogenetic reconstruction and molecular dating. Bioinformatics [Internet] 25:2286–2288. Available from: 10.1093/bioinformatics/btp368

Lawniczak MKN, Darwin Tree of Life Barcoding collective, Wellcome Sanger Institute Tree of Life programme, Wellcome Sanger Institute Scientific Operations: DNA Pipelines collective, Tree of Life Core Informatics collective, Darwin Tree of Life Consortium. 2021. The genome sequence of the spiny starfish, Marthasterias glacialis (Linnaeus, 1758). Wellcome Open Res [Internet] 6:295. Available from: 10.12688/wellcomeopenres.17344.1

Lee Y, Kim B, Jung J, Koh B, Jhang SY, Ban C, Chi W-J, Kim S, Yu J. 2022. Chromosome-level genome assembly of Plazaster borealis sheds light on the morphogenesis of multiarmed starfish and its regenerative capacity. Gigascience [Internet] 11. Available from: 10.1093/gigascience/giac063

Leulier F, Lemaitre B. 2008. Toll-like receptors--taking an evolutionary approach. Nat. Rev. Genet. [Internet] 9:165–178. Available from: 10.1038/nrg2303

Levy Karin E, Mirdita M, Söding J. 2020. MetaEuk-sensitive, high-throughput gene discovery, and annotation for large-scale eukaryotic metagenomics. Microbiome [Internet] 8:48. Available from: 10.1186/s40168-020-00808-x

Li H. 2018. Minimap2: pairwise alignment for nucleotide sequences. Bioinformatics [Internet] 34:3094–3100. Available from: 10.1093/bioinformatics/bty191

Li H. 2023. Protein-to-genome alignment with miniprot. Bioinformatics [Internet] 39:btad014. Available from: https://academic.oup.com/bioinformatics/advance-article/doi/10.1093/bioinformatics/btad014/6989621

Liu J, Zhou Y, Pu Y, Zhang H. 2023. A chromosome-level genome assembly of a deep-sea starfish (Zoroaster cf. ophiactis). Sci Data [Internet] 10:506. Available from: 10.1038/s41597-023-02397-4

Livingston BT, Killian CE, Wilt F, Cameron A, Landrum MJ, Ermolaeva O, Sapojnikov V, Maglott DR, Buchanan AM, Ettensohn CA. 2006. A genome-wide analysis of biomineralization-related proteins in the sea urchin Strongylocentrotus purpuratus. Dev. Biol. [Internet] 300:335–348. Available from: 10.1016/j.ydbio.2006.07.047

Li Y, Omori A, Flores RL, Satterfield S, Nguyen C, Ota T, Tsurugaya T, Ikuta T, Ikeo K, Kikuchi M, et al. 2020. Genomic insights of body plan transitions from bilateral to pentameral symmetry in Echinoderms. Commun Biol [Internet] 3:371. Available from: 10.1038/s42003-020-1091-1

Li Y, Wang R, Xun X, Wang J, Bao L, Thimmappa R, Ding J, Jiang J, Zhang L, Li T, et al. 2018. Sea cucumber genome provides insights into saponin biosynthesis and aestivation regulation. Cell Discov [Internet] 4:29. Available from: 10.1038/s41421-018-0030-5

Loof TG, Schmidt O, Herwald H, Theopold U. 2011. Coagulation systems of invertebrates and vertebrates and their roles in innate immunity: the same side of two coins? J. Innate Immun. [Internet] 3:34–40. Available from: 10.1159/000321641

Love MI, Huber W, Anders S. 2014. Moderated estimation of fold change and dispersion for RNA-seq data with DESeq2. Genome Biol. [Internet] 15:550. Available from: 10.1186/s13059-014-0550-8

Lowe CJ, Wray GA. 1997. Radical alterations in the roles of homeobox genes during echinoderm evolution. Nature [Internet] 389:718–721. Available from: 10.1038/39580

Mallefet J. 2009. Echinoderm bioluminescence: Where, how and why do so many ophiuroids glowIn: Meyer-Rochow VB, editor. Bioluminescence in focus: a collection of illuminating essays. Research Signpost. p. 67–83. Available from: https://dial.uclouvain.be/pr/boreal/object/boreal:130799

Mapleson D, Venturini L, Kaithakottil G, Swarbreck D. 2018. Efficient and accurate detection of splice junctions from RNA-seq with Portcullis. Gigascience [Internet] 7. Available from: 10.1093/gigascience/giy131

Marçais G, Kingsford C. 2011. A fast, lock-free approach for efficient parallel counting of occurrences of k-mers. Bioinformatics [Internet] 27:764–770. Available from: 10.1093/bioinformatics/btr011

Marlétaz F, Couloux A, Poulain J, Labadie K, Da Silva C, Mangenot S, Noel B, Poustka AJ, Dru P, Pegueroles C, et al. 2023. Analysis of the P. lividus sea urchin genome highlights contrasting trends of genomic and regulatory evolution in deuterostomes. Cell Genom [Internet] 3:100295. Available from: 10.1016/j.xgen.2023.100295

Martin M. 2011. Cutadapt removes adapter sequences from high-throughput sequencing reads. EMBnet.journal [Internet] 17:10–12. Available from: https://journal.embnet.org/index.php/embnetjournal/article/view/200

McCroskery S, Thomas M, Platt L, Hennebry A, Nishimura T, McLeay L, Sharma M, Kambadur R. 2005. Improved muscle healing through enhanced regeneration and reduced fibrosis in myostatin-null mice. J. Cell Sci. [Internet] 118:3531–3541. Available from: 10.1242/jcs.02482

McGinnis S, Madden TL. 2004. BLAST: at the core of a powerful and diverse set of sequence analysis tools. Nucleic Acids Res. [Internet] 32:W20–W25. Available from: 10.1093/nar/gkh435

Medina-Feliciano JG, García-Arrarás JE. 2021. Regeneration in Echinoderms: Molecular Advancements. Front Cell Dev Biol [Internet] 9:768641. Available from: 10.3389/fcell.2021.768641

Mendes FK, Vanderpool D, Fulton B, Hahn MW. 2020. CAFE 5 models variation in evolutionary rates among gene families. Bioinformatics [Internet] 36:5516–5518. Available from: https://academic.oup.com/bioinformatics/article/36/22-23/5516/6039105

Mongiardino Koch N, Thompson JR, Hiley AS, McCowin MF, Armstrong AF, Coppard SE, Aguilera F, Bronstein O, Kroh A, Mooi R, et al. 2022. Phylogenomic analyses of echinoid diversification prompt a re-evaluation of their fossil record. Elife [Internet] 11. Available from: 10.7554/eLife.72460

Mooi R, David B. 2008. Radial Symmetry, the Anterior/Posterior Axis, and Echinoderm Hox Genes. Annu. Rev. Ecol. Evol. Syst. [Internet] 39:43–62. Available from: 10.1146/annurev.ecolsys.39.110707.173521

Morel B, Kozlov AM, Stamatakis A, Szöllősi GJ. 2020. GeneRax: A Tool for Species-Tree-Aware Maximum Likelihood-Based Gene Family Tree Inference under Gene Duplication, Transfer, and Loss. Mol. Biol. Evol. [Internet] 37:2763–2774. Available from: 10.1093/molbev/msaa141

Mosher CV, Watling L. 2009. Partners for life: a brittle star and its octocoral host. Mar. Ecol. Prog. Ser. [Internet] 397:81–88. Available from: http://www.int-res.com/abstracts/meps/v397/p81-88/

Nei M, Gu X, Sitnikova T. 1997. Evolution by the birth-and-death process in multigene families of the vertebrate immune system. Proc. Natl. Acad. Sci. U. S. A. [Internet] 94:7799–7806. Available from: 10.1073/pnas.94.15.7799

Niknafs YS, Pandian B, Iyer HK, Chinnaiyan AM, Iyer MK. 2017. TACO produces robust multisample transcriptome assemblies from RNA-seq. Nat. Methods [Internet] 14:68–70. Available from: 10.1038/nmeth.4078

Novinec M, Kordis D, Turk V, Lenarcic B. 2006. Diversity and evolution of the thyroglobulin type-1 domain superfamily. Mol. Biol. Evol. [Internet] 23:744–755. Available from: 10.1093/molbev/msj082

O’Hara TD, Hugall AF, Thuy B, Moussalli A. 2014. Phylogenomic resolution of the class Ophiuroidea unlocks a global microfossil record. Curr. Biol. [Internet] 24:1874–1879. Available from: 10.1016/j.cub.2014.06.060

O’Hara TD, Hugall AF, Woolley SNC, Bribiesca-Contreras G, Bax NJ. 2019. Contrasting processes drive ophiuroid phylodiversity across shallow and deep seafloors. Nature [Internet] 565:636–639. Available from: 10.1038/s41586-019-0886-z

Oh SK, Kim D, Kim K, Boo K, Yu YS, Kim IS, Jeon Y, Im S-K, Lee S-H, Lee JM, et al. 2019. RORα is crucial for attenuated inflammatory response to maintain intestinal homeostasis. Proc. Natl. Acad. Sci. U. S. A. [Internet] 116:21140–21149. Available from: 10.1073/pnas.1907595116

Oliveri P, Davidson EH, McClay DR. 2003. Activation of pmar1 controls specification of micromeres in the sea urchin embryo. Dev. Biol. [Internet] 258:32–43. Available from: 10.1016/s0012-1606(03)00108-8

Oliveri P, Tu Q, Davidson EH. 2008. Global regulatory logic for specification of an embryonic cell lineage. Proc. Natl. Acad. Sci. U. S. A. [Internet] 105:5955–5962. Available from: 10.1073/pnas.0711220105

Open2C, Abdennur N, Fudenberg G, Flyamer IM, Galitsyna AA, Goloborodko A, Imakaev M, Venev SV. 2023. Pairtools: from sequencing data to chromosome contacts. bioRxiv [Internet]:2023.02.13.528389. Available from: https://www.biorxiv.org/content/10.1101/2023.02.13.528389v1

Pertea M, Pertea GM, Antonescu CM, Chang T-C, Mendell JT, Salzberg SL. 2015. StringTie enables improved reconstruction of a transcriptome from RNA-seq reads. Nat. Biotechnol. [Internet] 33:290–295. Available from: 10.1038/nbt.3122

Philippe H, Poustka AJ, Chiodin M, Hoff KJ, Dessimoz C, Tomiczek B, Schiffer PH, Müller S, Domman D, Horn M, et al. 2019. Mitigating Anticipated Effects of Systematic Errors Supports Sister-Group Relationship between Xenacoelomorpha and Ambulacraria. Curr. Biol. [Internet] 29:1818–1826.e6. Available from: 10.1016/j.cub.2019.04.009

Piovani L, Czarkwiani A, Ferrario C, Sugni M, Oliveri P. 2021. Ultrastructural and molecular analysis of the origin and differentiation of cells mediating brittle star skeletal regeneration. BMC Biol. [Internet] 19:9. Available from: 10.1186/s12915-020-00937-7

Pryzdial ELG, Leatherdale A, Conway EM. 2022. Coagulation and complement: Key innate defense participants in a seamless web. Front. Immunol. [Internet] 13:918775. Available from: 10.3389/fimmu.2022.918775

Purushothaman S, Saxena S, Meghah V, Swamy CVB, Ortega-Martinez O, Dupont S, Idris M. 2015. Transcriptomic and proteomic analyses of Amphiura filiformis arm tissue-undergoing regeneration. J. Proteomics [Internet] 112:113–124. Available from: 10.1016/j.jprot.2014.08.011

Putnam NH, Butts T, Ferrier DEK, Furlong RF, Hellsten U, Kawashima T, Robinson-Rechavi M, Shoguchi E, Terry A, Yu J-K, et al. 2008. The amphioxus genome and the evolution of the chordate karyotype. Nature [Internet] 453:1064–1071. Available from: 10.1038/nature06967

Quispe-Parra DJ, Medina-Feliciano JG, Cruz-González S, Ortiz-Zuazaga H, García-Arrarás JE. 2021. Transcriptomic analysis of early stages of intestinal regeneration in Holothuria glaberrima. Sci. Rep. [Internet] 11:346. Available from: 10.1038/s41598-020-79436-2

Rahman IA, Thompson JR, Briggs DEG, Siveter DJ, Siveter DJ, Sutton MD. 2019. A new ophiocistioid with soft-tissue preservation from the Silurian Herefordshire Lagerstätte, and the evolution of the holothurian body plan. Proc. Biol. Sci. [Internet] 286:20182792. Available from: 10.1098/rspb.2018.2792

Raible F, Tessmar-Raible K, Arboleda E, Kaller T, Bork P, Arendt D, Arnone MI. 2006. Opsins and clusters of sensory G-protein-coupled receptors in the sea urchin genome. Dev. Biol. [Internet] 300:461–475. Available from: 10.1016/j.ydbio.2006.08.070

Ramachandra R, Namburi RB, Dupont ST, Ortega-Martinez O, van Kuppevelt TH, Lindahl U, Spillmann D. 2017. A potential role for chondroitin sulfate/dermatan sulfate in arm regeneration in Amphiura filiformis. Glycobiology [Internet] 27:438–449. Available from: 10.1093/glycob/cwx010

Ranallo-Benavidez TR, Jaron KS, Schatz MC. 2020. GenomeScope 2.0 and Smudgeplot for reference-free profiling of polyploid genomes. Nat. Commun. [Internet] 11:1432. Available from: 10.1038/s41467-020-14998-3

Rast JP, Smith LC, Loza-Coll M, Hibino T, Litman GW. 2006. Genomic insights into the immune system of the sea urchin. Science [Internet] 314:952–956. Available from: 10.1126/science.1134301

Robinson MD, McCarthy DJ, Smyth GK. 2010. edgeR: a Bioconductor package for differential expression analysis of digital gene expression data. Bioinformatics [Internet] 26:139–140. Available from: 10.1093/bioinformatics/btp616

Robinson MD, Oshlack A. 2010. A scaling normalization method for differential expression analysis of RNA-seq data. Genome Biol. [Internet] 11:R25. Available from: 10.1186/gb-2010-11-3-r25

Saco A, Novoa B, Greco S, Gerdol M, Figueras A. 2023. Bivalves Present the Largest and Most Diversified Repertoire of Toll-Like Receptors in the Animal Kingdom, Suggesting Broad-Spectrum Pathogen Recognition in Marine Waters. Mol. Biol. Evol. [Internet] 40. Available from: 10.1093/molbev/msad133

Saotome K, Komatsu M. 2002. Chromosomes of Japanese starfishes. Zoolog. Sci. [Internet] 19:1095–1103. Available from: 10.2108/zsj.19.1095

Schiffer PH, Natsidis P, Leite DJ, Robertson H, Lapraz F, Marlétaz F, Fromm B, Baudry L, Simpson F, Høye E, et al. 2022. The slow evolving genome of the xenacoelomorph worm Xenoturbella bocki. bioRxiv [Internet]:2022.06.24.497508. Available from: https://www.biorxiv.org/content/10.1101/2022.06.24.497508v1.abstract

Schloissnig S, Kawaguchi A, Nowoshilow S, Falcon F, Otsuki L, Tardivo P, Timoshevskaya N, Keinath MC, Smith JJ, Voss SR, et al. 2021. The giant axolotl genome uncovers the evolution, scaling, and transcriptional control of complex gene loci. Proc. Natl. Acad. Sci. U. S. A. [Internet] 118. Available from: 10.1073/pnas.2017176118

Sea Urchin Genome Sequencing Consortium, Sodergren E, Weinstock GM, Davidson EH, Cameron RA, Gibbs RA, Angerer RC, Angerer LM, Arnone MI, Burgess DR, et al. 2006. The genome of the sea urchin Strongylocentrotus purpuratus. Science [Internet] 314:941–952. Available from: 10.1126/science.1133609

Seo HC, Kube M, Edvardsen RB, Jensen MF, Beck A, Spriet E, Gorsky G, Thompson EM, Lehrach H, Reinhardt R, et al. 2001. Miniature genome in the marine chordate Oikopleura dioica. Science [Internet] 294:2506. Available from: 10.1126/science.294.5551.2506

Simakov O, Bredeson J, Berkoff K, Marletaz F, Mitros T, Schultz DT, O’Connell BL, Dear P, Martinez DE, Steele RE, et al. 2022. Deeply conserved synteny and the evolution of metazoan chromosomes. Sci Adv [Internet] 8:eabi5884. Available from: 10.1126/sciadv.abi5884

Simakov O, Marlétaz F, Yue J-X, O’Connell B, Jenkins J, Brandt A, Calef R, Tung C-H, Huang T-K, Schmutz J, et al. 2020. Deeply conserved synteny resolves early events in vertebrate evolution. Nat Ecol Evol [Internet] 4:820–830. Available from: 10.1038/s41559-020-1156-z

Simão FA, Waterhouse RM, Ioannidis P, Kriventseva EV, Zdobnov EM. 2015. BUSCO: assessing genome assembly and annotation completeness with single-copy orthologs. Bioinformatics [Internet] 31:3210–3212. Available from: 10.1093/bioinformatics/btv351

Sinigaglia C, Almazán A, Lebel M, Sémon M, Gillet B, Hughes S, Edsinger E, Averof M, Paris M. 2022. Distinct gene expression dynamics in developing and regenerating crustacean limbs. Proc. Natl. Acad. Sci. U. S. A. [Internet] 119:e2119297119. Available from: 10.1073/pnas.2119297119

Sköld M, Rosenberg R. 1996. Arm regeneration frequency in eight species of ophiuroidea (Echinodermata) from European sea areas. J. Sea Res. [Internet] 35:353–362. Available from: https://www.sciencedirect.com/science/article/pii/S1385110196907625

Slota LA, McClay DR. 2018. Identification of neural transcription factors required for the differentiation of three neuronal subtypes in the sea urchin embryo. Dev. Biol. [Internet] 435:138–149. Available from: 10.1016/j.ydbio.2017.12.015

Slota LA, Miranda EM, McClay DR. 2019. Spatial and temporal patterns of gene expression during neurogenesis in the sea urchin Lytechinus variegatus. Evodevo [Internet] 10:2. Available from: 10.1186/s13227-019-0115-8

Smith AB. 2008. Deuterostomes in a twist: the origins of a radical new body plan. Evol. Dev. [Internet] 10:493–503. Available from: 10.1111/j.1525-142X.2008.00260.x

Srivastava M. 2021. Beyond Casual Resemblance: Rigorous Frameworks for Comparing Regeneration Across Species. Annu. Rev. Cell Dev. Biol. [Internet] 37:415–440. Available from: 10.1146/annurev-cellbio-120319-114716

Stanke M, Waack S. 2003. Gene prediction with a hidden Markov model and a new intron submodel. Bioinformatics [Internet] 19 Suppl 2:ii215–ii225. Available from: 10.1093/bioinformatics/btg1080

Stewart R, Rascón CA, Tian S, Nie J, Barry C, Chu L-F, Ardalani H, Wagner RJ, Probasco MD, Bolin JM, et al. 2013. Comparative RNA-seq analysis in the unsequenced axolotl: the oncogene burst highlights early gene expression in the blastema. PLoS Comput. Biol. [Internet] 9:e1002936. Available from: 10.1371/journal.pcbi.1002936

Stöhr S, O’Hara TD, Thuy B. 2012. Global diversity of brittle stars (Echinodermata: Ophiuroidea). PLoS One [Internet] 7:e31940. Available from: 10.1371/journal.pone.0031940

Suárez-Álvarez B, Liapis H, Anders H-J. 2016. Links between coagulation, inflammation, regeneration, and fibrosis in kidney pathology. Lab. Invest. [Internet] 96:378–390. Available from: 10.1038/labinvest.2015.164

Supek F, Bošnjak M, Škunca N, Šmuc T. 2011. REVIGO summarizes and visualizes long lists of gene ontology terms. PLoS One [Internet] 6:e21800. Available from: 10.1371/journal.pone.0021800

Telford MJ, Lowe CJ, Cameron CB, Ortega-Martinez O, Aronowicz J, Oliveri P, Copley RR. 2014. Phylogenomic analysis of echinoderm class relationships supports Asterozoa. Proc. Biol. Sci. [Internet] 281. Available from: 10.1098/rspb.2014.0479

Thuy B, Gale AS, Kroh A, Kucera M, Numberger-Thuy LD, Reich M, Stöhr S. 2012. Ancient origin of the modern deep-sea fauna. PLoS One [Internet] 7:e46913. Available from: 10.1371/journal.pone.0046913

Tu Q, Cameron RA, Davidson EH. 2014. Quantitative developmental transcriptomes of the sea urchin Strongylocentrotus purpuratus. Dev. Biol. [Internet] 385:160–167. Available from: 10.1016/j.ydbio.2013.11.019

Vaser R, Sović I, Nagarajan N, Šikić M. 2017. Fast and accurate de novo genome assembly from long uncorrected reads. Genome Res. [Internet] 27:737–746. Available from: 10.1101/gr.214270.116

Venturini L, Caim S, Kaithakottil GG, Mapleson DL, Swarbreck D. 2018. Leveraging multiple transcriptome assembly methods for improved gene structure annotation. Gigascience [Internet] 7. Available from: 10.1093/gigascience/giy093

Villot R, Poirier A, Devillers R, Kolnoguz A, Elowe S, Manem VSK, Joubert P, Mallette FA, Laplante M. 2021. ZNF768: controlling cellular senescence and proliferation with ten fingers. Mol Cell Oncol [Internet] 8:1985930. Available from: 10.1080/23723556.2021.1985930

Vistisen B, Vismann B. 1997. Tolerance to low oxygen and sulfide in Amphiura filiformis and Ophiura albida (Echinodermata: Ophiuroidea). Mar. Biol. [Internet] 128:241–246. Available from: 10.1007/s002270050088

Vopel K, Thistle D, Rosenberg R. 2003. Effect of the brittle starAmphiura filiformis(Amphiuridae, Echinodermata) on oxygen flux into the sediment. Limnol. Oceanogr. [Internet] 48:2034–2045. Available from: http://doi.wiley.com/10.4319/lo.2003.48.5.2034

Wu TD, Watanabe CK. 2005. GMAP: a genomic mapping and alignment program for mRNA and EST sequences. Bioinformatics [Internet] 21:1859–1875. Available from: 10.1093/bioinformatics/bti310

Wu T, Hu E, Xu S, Chen M, Guo P, Dai Z, Feng T, Zhou L, Tang W, Zhan L, et al. 2021. clusterProfiler 4.0: A universal enrichment tool for interpreting omics data. Innovation (Camb) [Internet] 2:100141. Available from: 10.1016/j.xinn.2021.100141

Xu N, Lao Y, Zhang Y, Gillespie DA. 2012. Akt: a double-edged sword in cell proliferation and genome stability. J. Oncol. [Internet] 2012:951724. Available from: 10.1155/2012/951724

Yamazaki A, Morino Y, Urata M, Yamaguchi M, Minokawa T, Furukawa R, Kondo M, Wada H. 2020. pmar1/phb homeobox genes and the evolution of the double-negative gate for endomesoderm specification in echinoderms. Development [Internet] 147. Available from: 10.1242/dev.182139

Yan A, Ren C, Chen T, Jiang X, Sun H, Hu C. 2016. Identification and functional characterization of a novel antistasin/WAP-like serine protease inhibitor from the tropical sea cucumber, Stichopus monotuberculatus. Fish Shellfish Immunol. [Internet] 59:203–212. Available from: 10.1016/j.fsi.2016.10.038

Yan H, Bombarely A, Li S. 2020. DeepTE: a computational method for de novo classification of transposons with convolutional neural network. Bioinformatics [Internet] 36:4269–4275. Available from: 10.1093/bioinformatics/btaa519

Yates AD, Allen J, Amode RM, Azov AG, Barba M, Becerra A, Bhai J, Campbell LI, Carbajo Martinez M, Chakiachvili M, et al. 2022. Ensembl Genomes 2022: an expanding genome resource for non-vertebrates. Nucleic Acids Res. [Internet] 50:D996–D1003. Available from: 10.1093/nar/gkab1007

Zhang X, Sun L, Yuan J, Sun Y, Gao Y, Zhang L, Li S, Dai H, Hamel J-F, Liu C, et al. 2017. The sea cucumber genome provides insights into morphological evolution and visceral regeneration. PLoS Biol. [Internet] 15:e2003790. Available from: 10.1371/journal.pbio.2003790

Zhou C, McCarthy SA, Durbin R. 2023. YaHS: yet another Hi-C scaffolding tool. Bioinformatics [Internet] 39. Available from: 10.1093/bioinformatics/btac808

Zhou Y, Chen JJ. 2021. STAT3 plays an important role in DNA replication by turning on WDHD1. Cell Biosci. [Internet] 11:10. Available from: 10.1186/s13578-020-00524-x

