## Supplemental Material for "The brittle star genome illuminates the genetic basis of animal appendage regeneration"

Parey et al. 2023

#### Supplemental Note

**Note S1**: Comparisons with previous efforts to sequence genomes from the brittle star group

#### Supplemental Tables

**Table S1:** Accessions for *A. filiformis* RNA-seq datasets. [Supplemental_TableS1.xlsx]
**Table S2:** *A. filiformis* gene models properties. [Supplemental_TableS2.xlsx]
**Table S3:** Enriched gene ontology terms in brittle star expanded and contracted gene families. [Supplemental_TableS3.xlsx]
**Table S4:** Enriched gene ontology terms in brittle star arm regeneration co-expression clusters. [Supplemental_TableS4.xlsx]
**Table S5:** Axolotl limb regeneration co-expression clusters and gene ontology enrichment. [Supplemental_TableS5.xlsx]
**Table S6:** Genes with conserved expression profiles during appendage regeneration in brittle star, axolotl and Parhyale, and associated gene ontology term enrichments. [Supplemental_TableS6.xlsx]

#### Supplemental Figures

**Figure S1:** Genome assembly and repeat annotation.
**Figure S2:** Reconstruction of the ancestral Euleterozoa linkage groups (ELG).
**Figure S3:** Inter-chromosomal macrosyntenic rearrangements since the Eleutherozoa divergence in the sea urchin *P. lividus* and brittle star *A. filiformis*.
**Figure S4:** Molecular phylogeny of echinoderm Hox genes.
**Figure S5:** Evolution of the *pmar1/phb* and luciferase genes by tandem duplications.
**Figure S6:** Clustering of the brittle star arm regeneration time course gene expression clustering.
**Figure S7:** Gene ontology enrichment results for brittle star arm regeneration co-expression clusters.
**Figure S8:** Clustering and functional enrichments for the axolotl and Parhyale limb regeneration gene expression time series.

**Figure S9:** Comparison of co-expression gene clusters in regeneration and development.
**Figure S10:** Gene lists enrichment for genes with a conserved expression profile during appendage regeneration.
**Figure S11:** Differential transcriptional activity of brittle star repetitive elements in immune and proliferative regeneration phases.

#### Supplemental Datasets

The following datasets have been deposited in a Zenodo repository (https://zenodo.org/doi/10.5281/zenodo.10036671):

**Dataset S1:** Gene and repeat annotation (including .fasta genome file, .bed, .gtf and .fasta files for *A. filiformis* genes and repeat annotations for the four genomes presented in Figure 1D). [Dataset_s1]

**Dataset S2:** Annotation of selected echinoderm genomes with respect to the predicted Eleutherozoa ancestral Linkage Groups (.bed files with genes annotated with their ancestral chromosome of origin and orthologous genes files). [Dataset_s2]

**Dataset S3:** Hox and ParaHox genes in A. filiformis (including Hox protein sequences and molecular phylogeny, and ParaHox sequences). [Dataset_s3]

**Dataset S4:** Echinoderm gene families, results of the expansion/contraction tests and alignments and trees for the luciferase and *pmar1/phb* gene families. [Dataset_s4]

**Dataset S5:** Raw counts and normalised gene expression tables for *A. filiformis* development, regeneration time course and explant experiments, the *A. mexicanum* limb regeneration time series and predicted homologs between *A. filiformis, A. mexicanum and P. hawaiensis.* [Dataset_s5]

### Supplemental Note

**Note S1: Comparisons with previous efforts to sequence genomes from the brittle star class Ophiuroidea.**

To date, and excluding our newly-generated high-quality *A. filiformis* genome, only three ophiuroid species have a draft genome assembly in the NCBI genome database: *Ophioderma brevispina*, *Ophionereis fasciata* and *Ophiothrix spiculata*. Only two of these genomes have been previously published: *Ophioderma brevispina* (Mashanov et al. 2022) and *Ophionereis fasciata (Long et al. 2016)*. Critically, none of these sub-chromosomal assemblies have been found sufficiently robust for inclusion in the authoritative Database Echinobase (Arshinoff et al. 2022). Producing high-quality genomic resources for the brittle star class has thus represented a significant challenge, probably due to their relatively high genome size compared to other echinoderms, elevated repeat content and high level of heterozygosity. Based on long-read nanopore reads and proximity ligation data, and refinement with illumina short reads, our *A. filiformis* genome greatly improves upon the previously published ophiuroid draft genomes. The *A. filiformis* assembly has markedly higher completeness (96.1% vs 30% assembly complete BUSCO) and contiguity (68.8 Mb vs 48.5 kb N50) as well as a more realistic number of predicted genes (30,267 vs 146,703) than *Ophioderma brevispina* (Mashanov et al. 2022). Similarly, the provided assembly statistics from the low-coverage *Ophionereis fasciata* genome *(Long et al. 2016)* (N50: 72.8 kb; gene numbers: 102,838) showcase the significant improvements of our *A. filiformis* genome over the state-of-the-art.

#### Supplemental Tables

**Table S1: Accessions for *A. filiformis* RNA-seq datasets.** [Supplemental_TableS1.xlsx].
This table lists all RNA-seq datasets used and/or generated in this study, along with the corresponding SRA accession numbers. The datasets used for the genome annotation are also indicated.

**Table S2: *A. filiformis* gene models properties.** [Supplemental_TableS2.xlsx]
This table presents the 30,267 predicted gene models, along with their properties, including: genomic location (chromosome or scaffold, start, end, strand columns), predicted gene name based on a blast with the swissprot database, predicted gene age, expression sample with maximal TPM (Max TPM, Max Sample) within the normalised "development and arm regeneration" dataset (**Table S1, Dataset S5**), cluster membership for the regeneration time course, differential expression in explant experiment, curated gene list membership, PFAM Domains, and a reduced set of associated Gene Ontology Terms. The second sheet contains details for the genes annotated to the 19 signalling pathways. The first 19 columns contain the 19 pathways, where a '1' in a given column represents membership to the pathway, additional columns contain the *A. filiformis* gene names and *S. purpuratus* orthologs.

**Table S3: Enriched gene ontology terms in brittle star expanded and contracted gene families.** [Supplemental_TableS3.xlsx]
Gene ontology enrichment results for each brittle star arm regeneration co-expression cluster, with columns as follows: GO term ID, GO term description, GO dispensability score, enrichment, adjusted p-value, corresponding gene families identifiers and brittle star genes.

**Table S4: Enriched gene ontology terms in brittle star arm regeneration co-expression clusters.** [Supplemental_TableS4.xlsx]
Gene ontology enrichment results for each brittle star arm regeneration co-expression cluster, with columns as follows: GO term ID, GO term description, fraction of genes in the cluster annotated with the GO term, fraction of background genes annotated with the GO term, p-value, adjusted p-value, q-value, corresponding genes and number of genes.

**Table S5: Axolotl limb regeneration co-expression clusters and gene ontology enrichment.** [Supplemental_TableS5.xlsx]
Gene ontology enrichment results for each axolotl limb regeneration co-expression cluster, as in Table S3. Axolotl gene ID corresponds to the AmexG_v6.0-DD assembly version.

**Table S6: Genes with conserved expression profiles during appendage regeneration in brittle star, axolotl and Parhyale, and enriched ontology terms.** [Supplemental_TableS6.xlsx]

List of genes co-expressed during regeneration in brittle star, axolotl and Parhyale, with columns as follows: species comparison in which the genes are conserved, co-expression clusters in these species, gene family ID, gene IDs and gene names, and curated gene list membership. The second sheet presents the full results of the Gene Ontology enrichment tests.

### Supplemental Figures

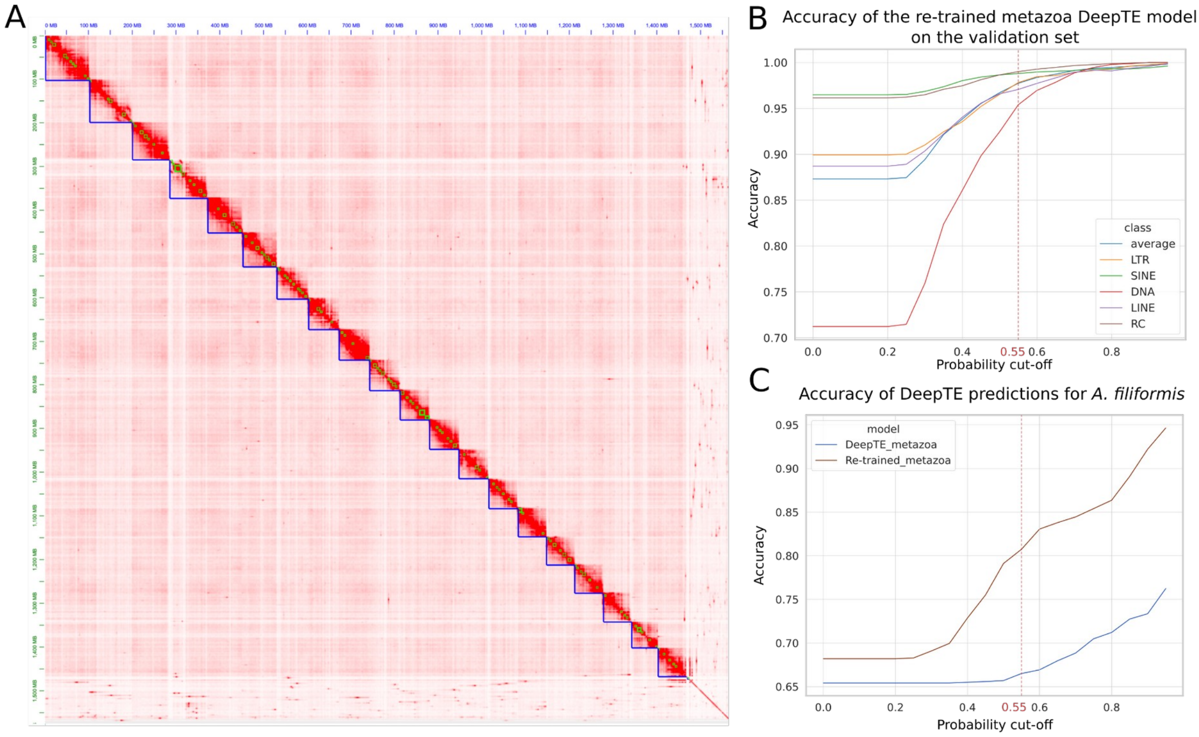


**Figure S1: Genome assembly and repeat classification. A.** Hi-C contact map showing the density of interactions between binned genomic regions in the proximity ligation data. The high contact regions are consistent with a 20 chromosome *A. filiformis* karyotype. **B.** Validation accuracy of a new DeepTE model (Yan et al. 2020), trained to classify repeats into 5 main classes: LTR, SINE, DNA, LINE and Rolling Circle (RC). The vertical dotted line corresponds to the calibrated 0.55 threshold that we used on the DeepTE scores to classify repetitive elements. **C.** Accuracy of the newly-trained and the default Metazoa DeepTE models on the test set of *A. filiformis* repeats. The accuracy of the new model is superior to the default model and can classify repeats into 5 as opposed to 3 classes (repeats of ClassI, ClassII and ClassIII).

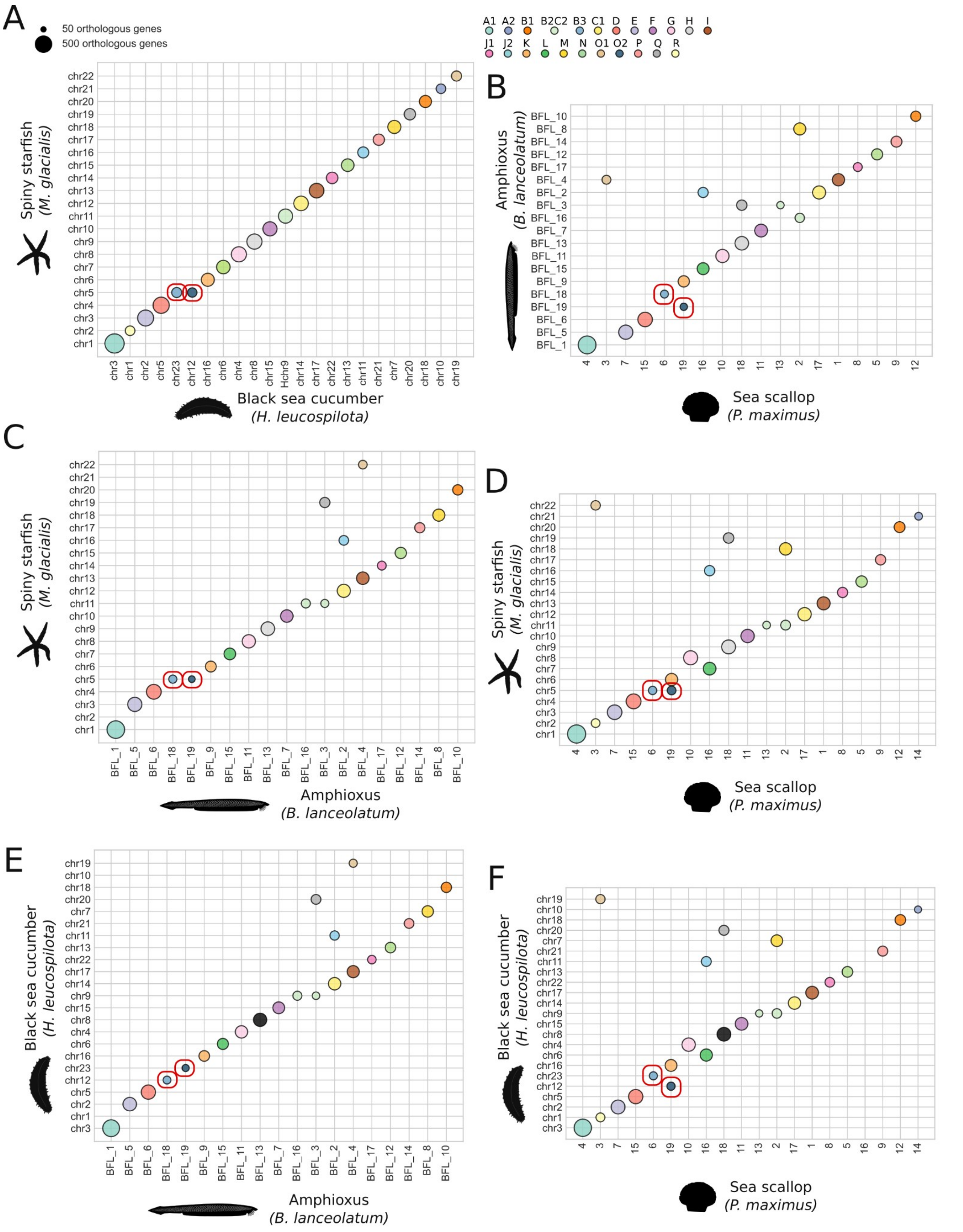


**Figure S2: Reconstruction of the ancestral Euleterozoa linkage groups (ELG). A.** Synteny comparison between spiny starfish and black sea cucumber reveals one macrosyntenic rearrangement (red boxes). ELGs colours are indicated at the top and correspond to colours on Figure 1. Pairwise synteny comparisons with Amphioxus and Sea Scallop are similarly displayed on **B.**, **C.**, **D.**, **E.** and **F**, with red boxes highlighting that B3, and O2 are all on distinct chromosomes in Amphioxus and Sea Scallop, thus confirming that the starfish B3-O2 fusion is starfish-specific derived rearrangement.

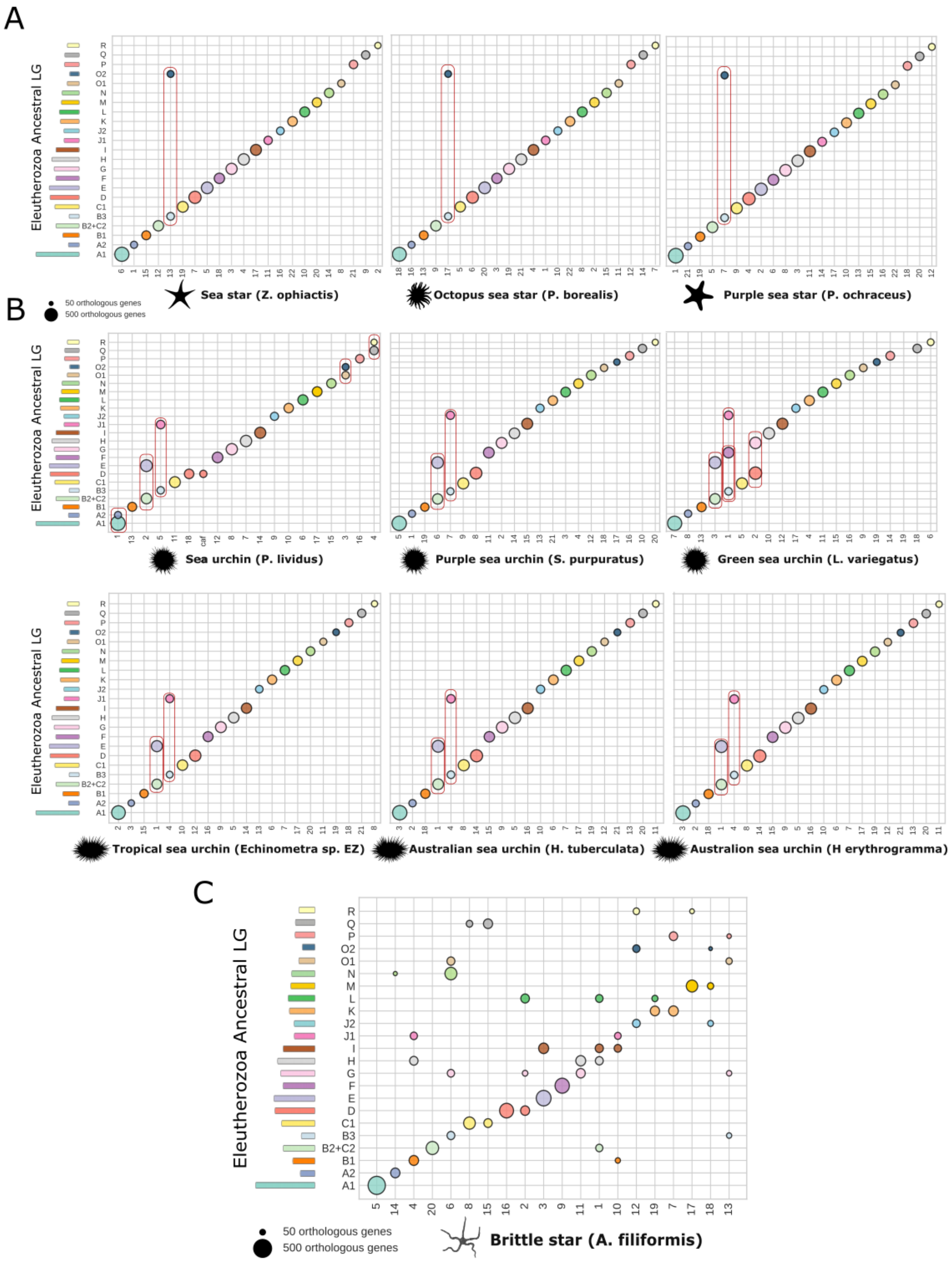
A
B
C
Eleutherozoa Ancestral LG
R
Q
P
O2
O1
N
M
L
K
J2
J1
I
H
G
F
E
D
C1
B3
B2+C2
B1
A2
A1
Sea star (Z. ophiactis)
Octopus sea star (P. borealis)
Purple sea star (P. ochraceus)
50 orthologous genes
500 orthologous genes
Sea urchin (P. lividus)
Purple sea urchin (S. purpuratus)
Green sea urchin (L. variegatus)
Tropical sea urchin (Echinometra sp. EZ)
Australian sea urchin (H. tuberculata)
Australion sea urchin (H erythrogramma)
Brittle star (A. filiformis)

**Figure S3: Inter-chromosomal macrosyntenic rearrangements since the Eleutherozoa ancestor in sequenced echinoderms. A.** Synteny comparison between ELGs and available chromosome-scale sea star genomes (Schiebelhut et al. 2018; Lee et al. 2022; Liu et al. 2023). All examined sea star genomes are marked by the single B3+O2 fusion. **B.** Synteny comparison between ELGs and available chromosome-scale sea urchin genomes (Sea Urchin Genome Sequencing Consortium et al. 2006; Davidson et al. 2020; Davidson et al. 2022; Ketchum et al. 2022; Marlétaz et al. 2023). All examined sea urchin genomes are marked by the (B2+C2)+E and B3+J1 fusion. *L. variegatus* underwent the additional (B3+J1)+J2 fusion and D+G. *P. lividus* underwent the additional A1+A2, O1+O2 and Q+R fusions (note that an additional fission of ELG D may have occurred if the large unplaced scaffold noted "Scaf." is not an assembly artefact.) **C.** Synteny comparison between ELGs and brittle star chromosomes reveals a total of 26 macrosyntenic inter-chromosomal rearrangements, in the most parsimonious scenario involving fusion, fission and translocation events. The rearrangements can be inferred from the oxford grid plot: 1 [fusion + mixing + fission] of 3 ELGs = 3 inter-chromosomal rearrangements (B3-G-O1), 3 [fusion + mixing + fission] of 2 ELGs = 6 inter-chromosomal rearrangements (B1-J1, C1-Q, J2-O2) and 17 translocations (A2-N, B1J1-H, B3GO1-N, D-G, DG-L, E-I, G-H, B2+C2-H, B2+C2H-I, B2+C2HI-L, B1J1-I, J2O2-R, K-L, K-P, M-R, J2O2-M, B3GO1-P).

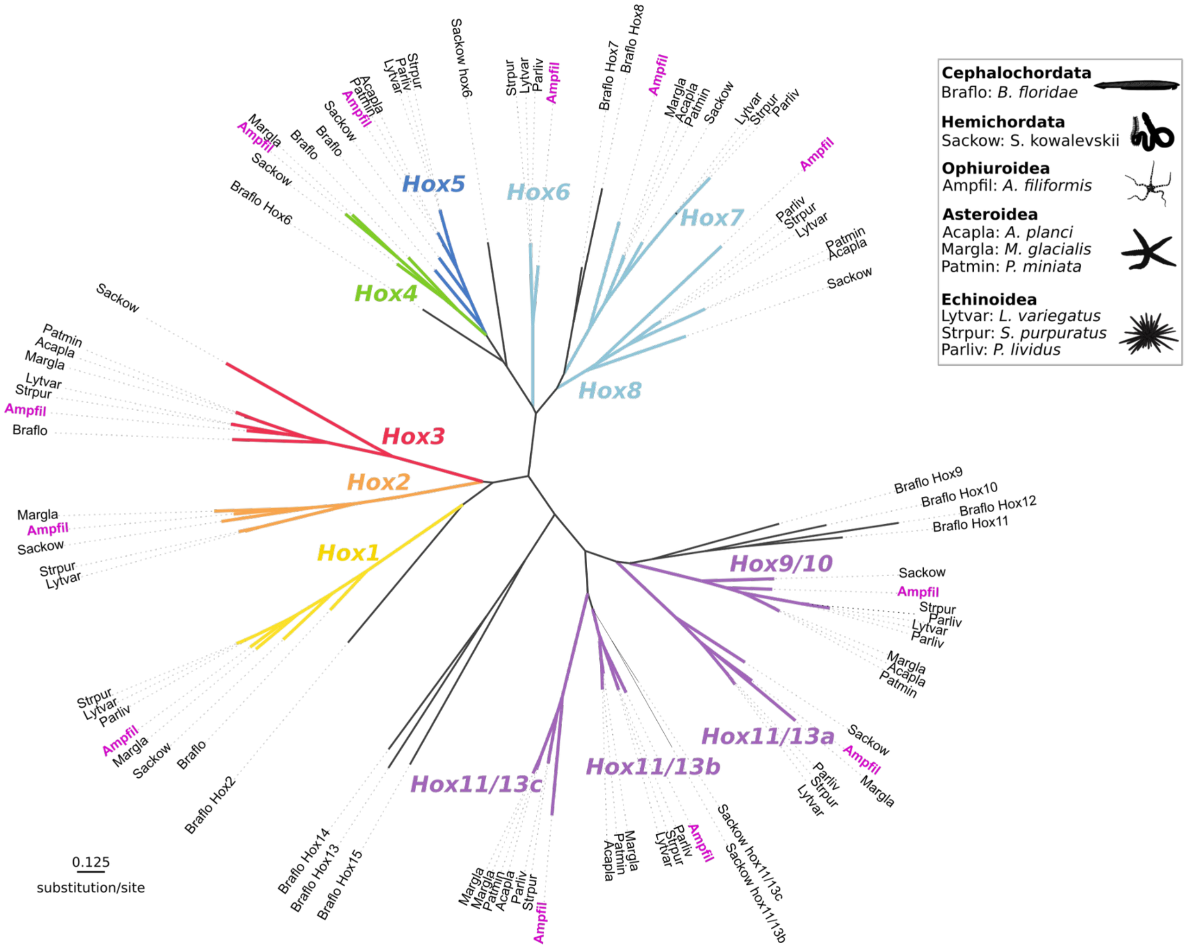


**Figure S4: Molecular phylogeny of echinoderm Hox genes.** The phylogenetic tree is shown as an unrooted tree, with clades of Hox genes indicated with the same colours as in **Figure 2**. The phylogenetic position of each identified Hox gene in the brittle star ("Ampfil") is highlighted in pink.

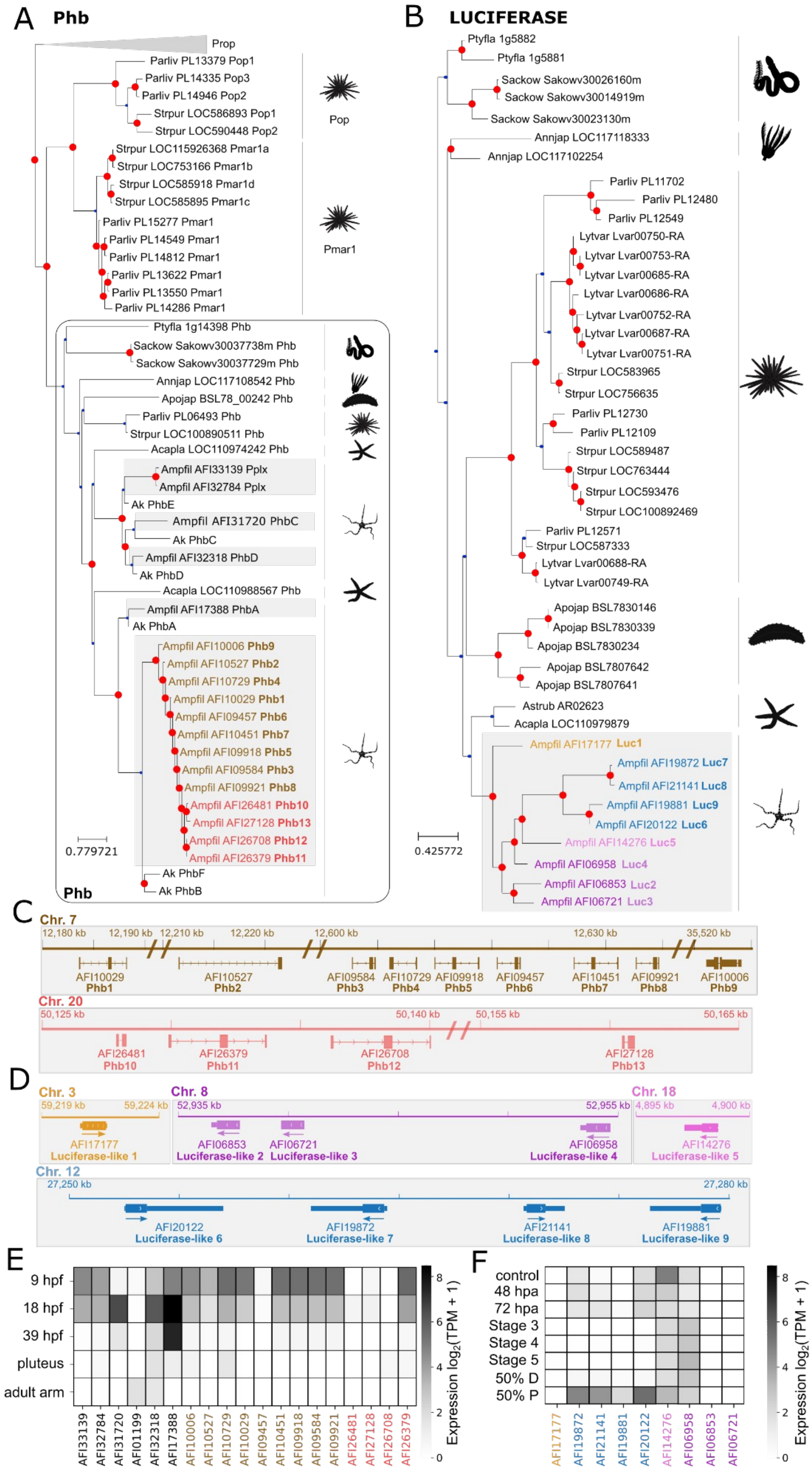


**Figure S5: Evolution of the *Pmar1/Phb* and *luciferase-like* genes by tandem duplications. A.** Molecular phylogeny of the Pmar1/Phb genes in echinoderms. The tree was reconstructed with RAxML-NG (Kozlov et al. 2019) (10 starting parsimony trees, 1000 bootstraps, LG+G4+F model), lowly supported nodes (bootstrap < 60) were subsequently corrected with Treerecs (Comte et al. 2020) to maximise the parsimony of duplications and losses. Species are indicated by abbreviations (Ptyfla = *P. flava*, Sackow = *S. kowalevskii*, Annjap = *A. japonica*, Parliv = *P. lividus*, Strpur = *S. purpuratus*, Apojap = *A. japonicus*, Acapla = *A. planci*, Ak = *A. kochii*, Ampfil = *A. filiformis*). Inferred duplication nodes are shown in red. *pmar1/phb* full gene sequences were identified based on (Yamazaki et al. 2020; Marlétaz et al. 2023) (see **Dataset S4**). **B.** Phylogeny of luciferase genes in echinoderms, as in **A**. Luciferase-like genes were identified based on sequences from (Delroisse et al. 2017) (**Methods,** see also **Dataset S4** for the sequences of the nine luciferase-like genes and two additional luciferase-like pseudogenes). **C.** Genomic location of tandem-duplicated *A. filiformis phb* genes. **D.** Genomic location of tandem-duplicated *A. filiformis* luciferase genes. **E.** *Phb* expression throughout 4 brittle star developmental time points and in the adult arm, showing the early developmental expression of *phb* genes (hpf: hours post-fertilization). Expression across samples was normalised using the TMM method (Robinson et al. 2010) on the full set of brittle star genes, and is shown as $log_2$(TPM + 1). **F.** Luciferase-like gene expression during brittle star arm regeneration, showing that most luciferase-like genes are expressed in differentiated arms only: control arms and the latest regeneration time point (hpa: hours post-amputation, see **Figure 4** for staging details). Expression normalisation as in **E**.

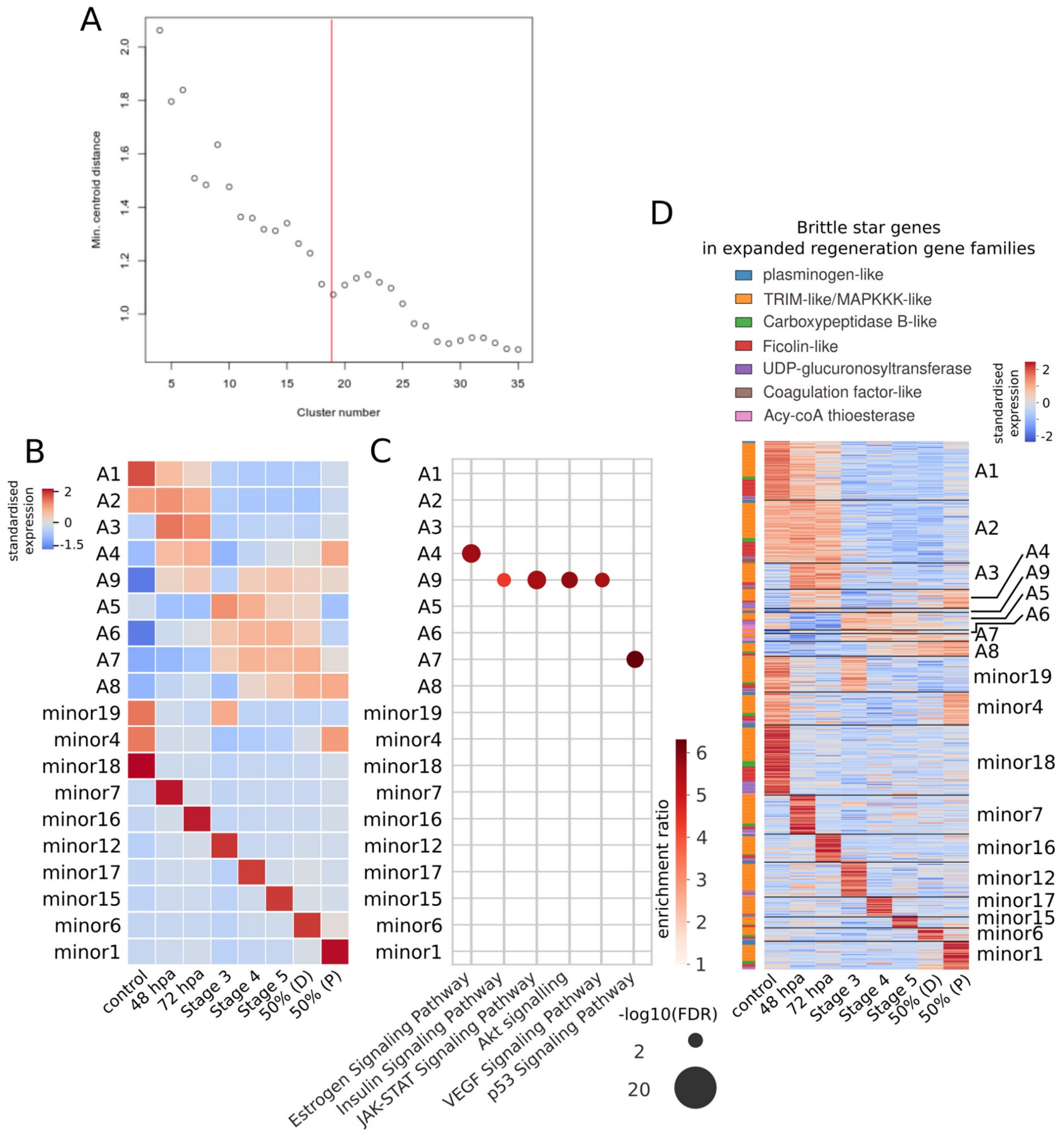


**Figure S6: Clustering of the brittle star arm regeneration time course gene. A**. Optimal number of clusters estimated using the centroid distance. We selected n=19 clusters since further increase of the number of clusters does not result in a significant decrease of the centroid distance. **B.** Normalised expression profiles (expression of the centroid) for each of the n=19 clusters. Clusters with genes expressed over a single regeneration time point (or one regeneration point + control) were defined as minor clusters and not presented in the main text as these typically do not display significant enrichments and may be driven by noisy gene expression. **C.** Signalling pathways enrichment for each co-expression cluster (hypergeometric test, Benjamini-Hochberg adjusted p-values <0.05**, Methods**). **D.** Expression throughout arm regeneration of the brittle star in the expanded gene families annotated with the GO term 'regeneration' (see **Figure 3B, 3C**). Gene family membership (correspondence with **Figure 3C**) are indicated with colours on the left of the expression heatmap, clusters are shown on the right.

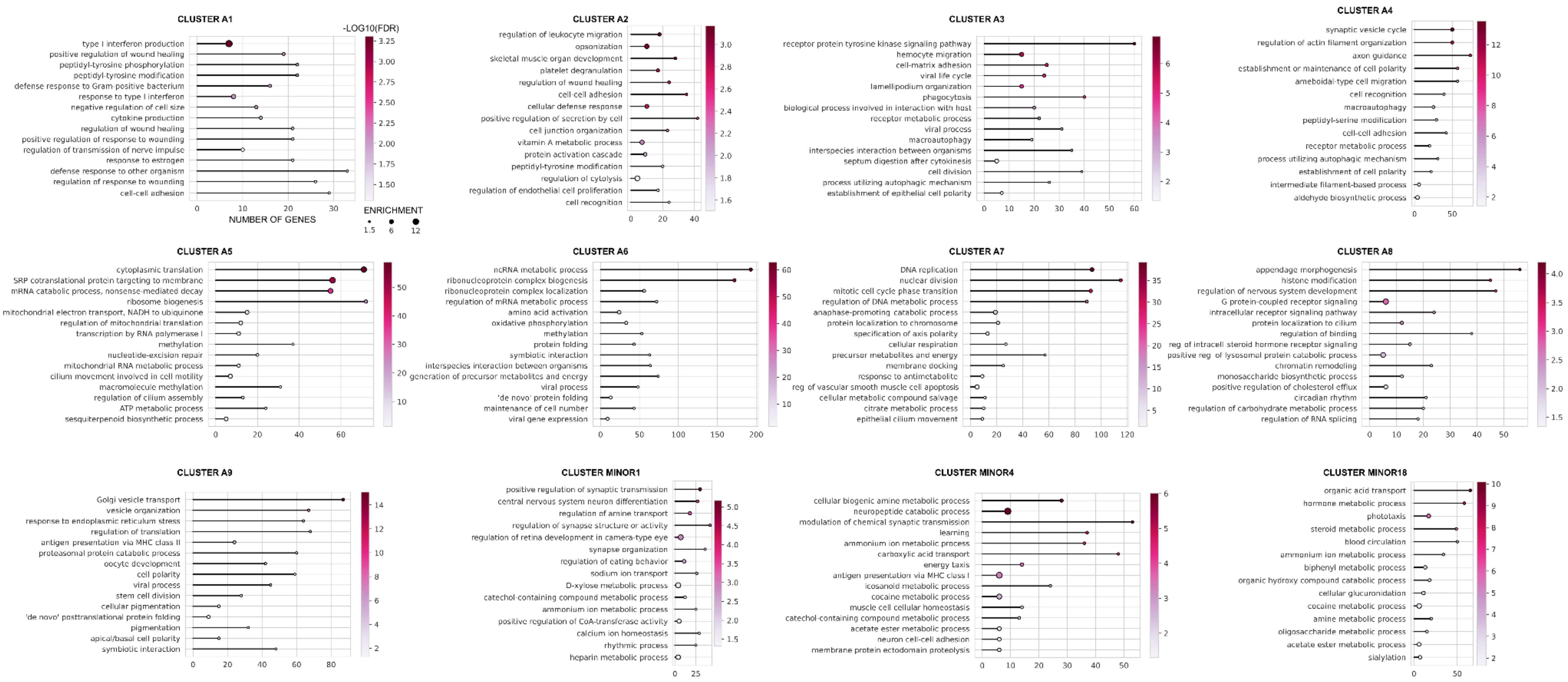


**Figure S7: Gene ontology enrichment results for brittle star arm regeneration co-expression clusters.** GO enrichment tests were performed on each co-expression cluster and summarised using REVIGO (**Methods**). The complete list of enriched terms is presented in **Table S3**.

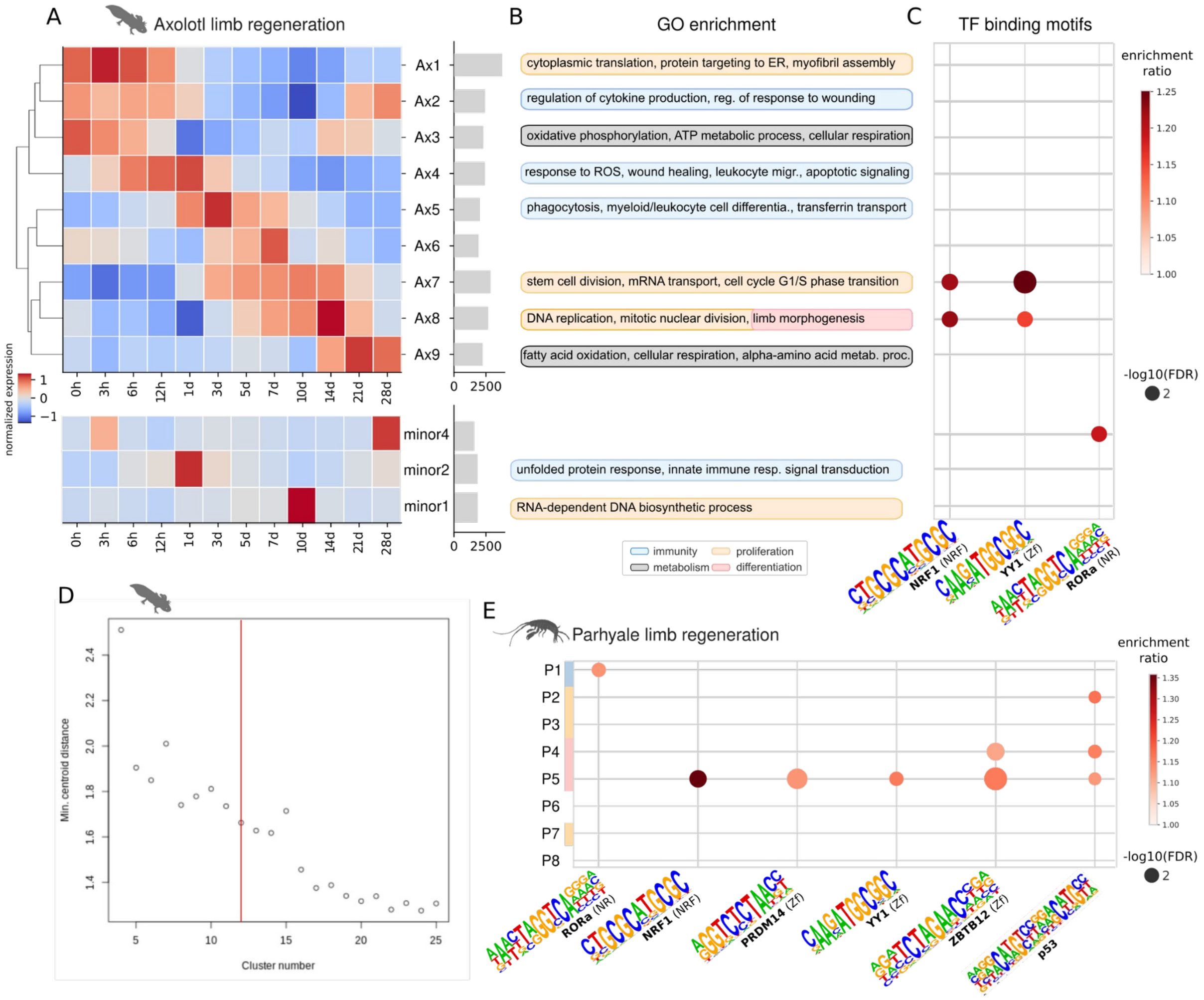


**Figure S8: Clustering and functional enrichments for the axolotl and Parhyale limb regeneration gene expression time series. A**. Normalised expression profiles (expression of the centroid) for each of the n=12 axolotl limb regeneration co-expression clusters. Raw expression data were re-processed from (Stewart et al. 2013) (**Methods**). Barplots on the right indicate the number of genes assigned to each cluster. Clusters with genes expressed over a single regeneration time point were defined as minor clusters and not presented in the main text as they may be driven by noisy gene expression. **B.** Gene ontology enrichment for each co-expression cluster (**Methods**, **Table S5**). **C.** TF binding motifs enriched around the TSS of genes from axolotl co-expression clusters (hypergeometric test adjusted p-value <0.05, **Methods**). Note that only TFBS motifs enriched in brittle star clusters are represented. **D.** Optimal number of clusters estimated using the centroid distance. We selected n=12 clusters since further increase of the number of clusters does not result in a significant decrease of the centroid distance until n=16, which, on the basis of functional enrichment tests, over-clusters the data. **E.** TF binding motifs enriched around the TSS of genes from Parhyale co-expression clusters as in **C**. Parhyale clusters were renamed from (Sinigaglia et al. 2022) as follows: P1 is R4 in the notation of Sinigaglia et al., P2 is R1, P3 is R8, P4 is R2, P5 is R6, P6 is R3, P7 is R5 and P8 is R7.

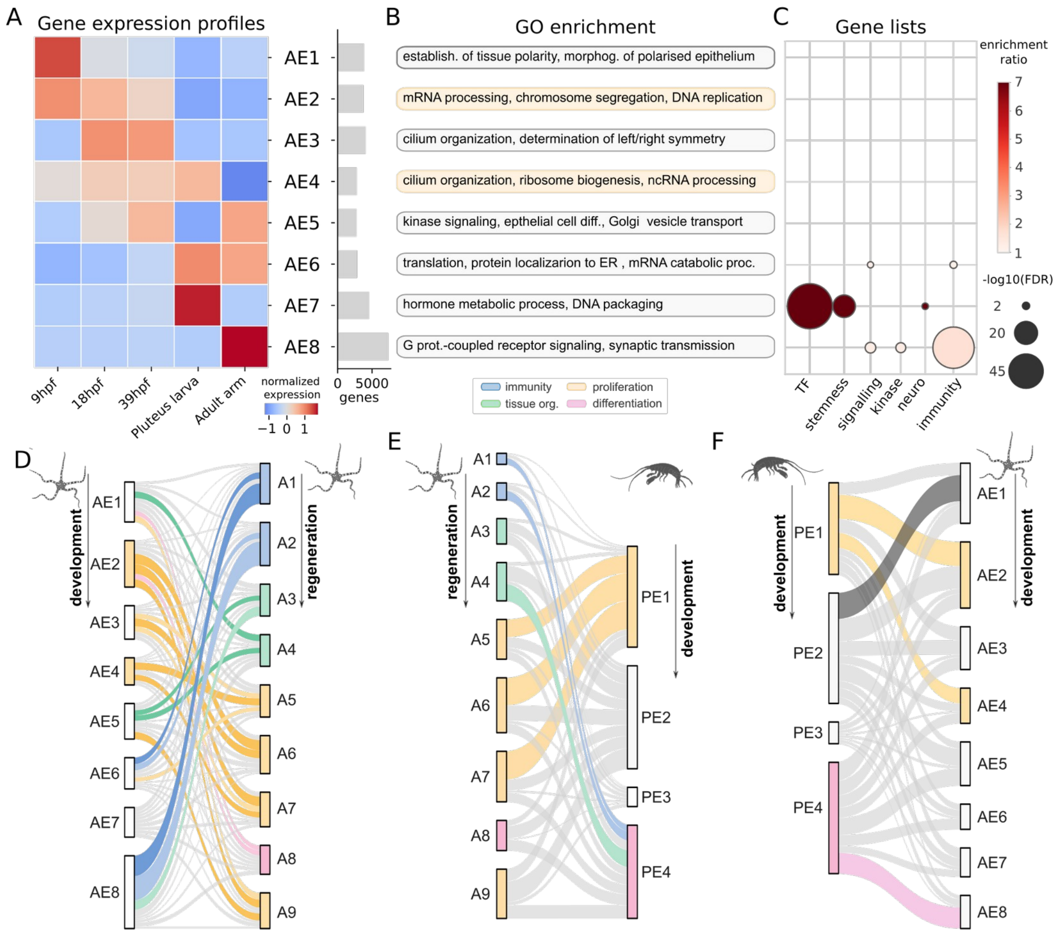


**Figure S9: Comparison of co-expression gene clusters during regeneration and development. A.** Clustering of the brittle star development time series. Normalised expression profiles for each of the n=8 development co-expression clusters. Processing, clustering procedure and representation is as in (**Figure 4**, **Figure S8)**, and based on RNA-seq datasets from (Delroisse et al. 2014; Dylus et al. 2016) (**Table S1**). **B.** Gene ontology enrichment for each co-expression cluster. **C.** Curated gene lists enrichment for each co-expression cluster (hypergeometric test, Benjamini-Hochberg adjusted p-values <0.05). **D.** Comparison of co-expressed gene clusters deployed during embryonic development and appendage regeneration in the brittle star. Note that the embryonic development in brittle star does not produce appendages and is thus less informative than Parhyale development data. Clusters are represented by vertical rectangles whose sizes are proportional to the number of homologous genes in the cluster, and coloured according to enriched GO terms. Genes are linked across clusters, with coloured links indicating significant overlaps (hypergeometric test with the Benjamini-Hochberg correction <0.01, darker shades indicate p-values < $10^{-15}$). **E.** Comparison of co-expressed gene clusters deployed during appendage regeneration in the brittle star and leg development in Parhyale. Clusters in Parhyale (clusters PE1 to PE4) correspond to the clustering reported in (Sinigaglia et al. 2022), but clusters were renamed to follow temporal activation (PE1 corresponds to E2, PE2 to E4, PE3 to E1, PE4 to E3). Coloured links indicating significant overlaps (permutation-based p-values with Benjamini-Hochberg correction <0.05, Methods). **F.** Comparison of co-expressed gene clusters deployed during development in the brittle star and leg development in Parhyale, as in **E**.

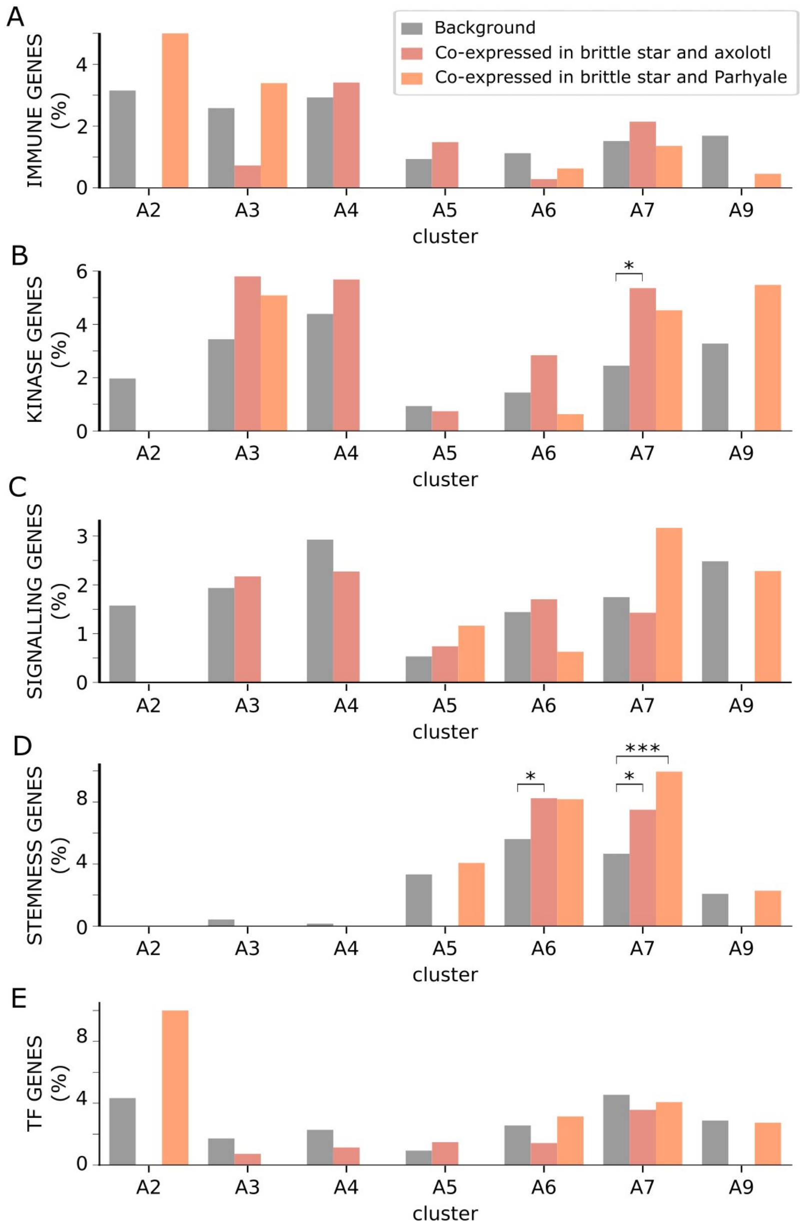


**Figure S10: Gene lists enrichment for genes with a conserved expression profile during appendage regeneration. A-E.** Gene list enrichment tests, as in **Figure 5D**, but sub-divided by cluster and species comparisons (hypergeometric tests, p-values corrected for multiple testing with the BH procedure, * p-values < 0.05, ** p-values < 0.01, *** p-values < 0.001).

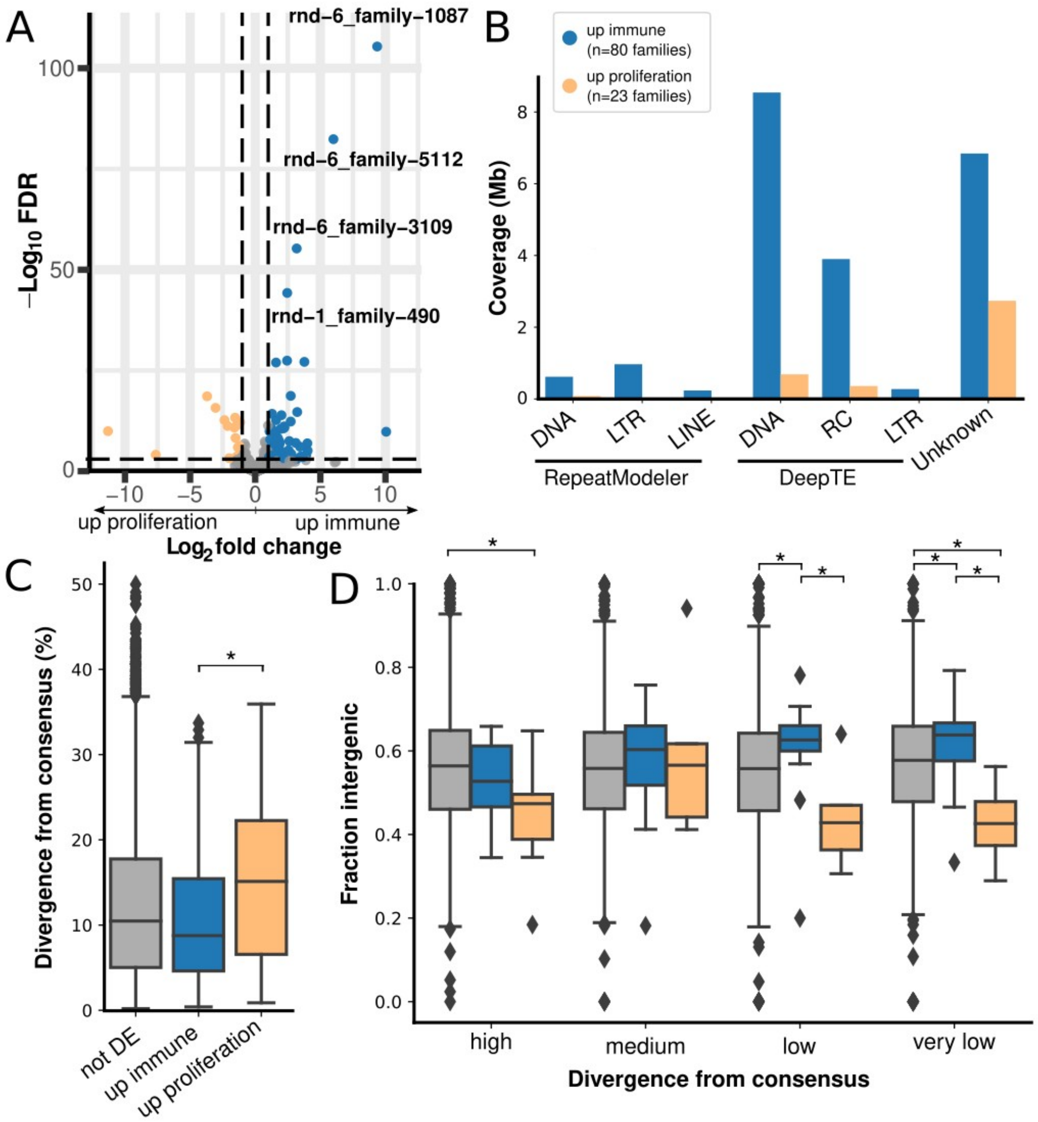


**Figure S11: Differential transcriptional activity of repetitive elements in the immune and proliferative phases of brittle star arm regeneration. A.** Differentially expressed repetitive elements in early regeneration (immune phase: 48 hpa and 72 hpa samples) versus middle regeneration (proliferation: Stage3, Stage4, Stage5 samples). Coloured dots represent repeat families with significant up-expression in immune (blue) or proliferation phases (orange) (absolute log fold change > 1, FDR < 0.001, **Methods**). **B.** Immune up-expressed repeat families (n=80) have a higher genomic coverage than proliferation up repeat families (n=23), regardless of repeat class. Coverage is shown subdivided by repeat class, where classification was performed first using the homology-based approach of RepeatModeler, then with DeepTE for repeats that could not be classified by RepeatModeler (**Methods**). We note that the DeepTE classification has higher false positives than the RepeatModeler classification. **C.** Immune up-expressed repeat families have significantly lower divergence to their consensus (Kimura distance, **Methods**) than proliferation up-expressed repeat families (Mann–Whitney U test, one-sided p-value corrected for multiple testing with the BH procedure, * p-values < 0.05), indicating they are younger repeats with a higher potential to still be active mobilisable transposable elements. **D.** Immune up-expressed repeat families with low divergence from their consensus have significantly higher fraction of intergenic repeat instances, suggesting up-expression is less likely to be a side-effect of host gene transcription. P-values and boxplot colours are as in **C** (grey = no significant differential expression, blue = up in immune, orange = up in proliferation). Repeat families were subdivided in 4 balanced categories based on their divergence to consensus (Kimura distance, d): $d < 5.02$ (very low), $5.02 < d < 10.48$ (low), $10.48 < d < 17.73$ (medium), $17.73 < d$ (high).

#### Supplemental references


Arshinoff BI, Cary GA, Karimi K, Foley S, Agalakov S, Delgado F, Lotay VS, Ku CJ, Pells TJ, Beatman TR, et al. 2022. Echinobase: leveraging an extant model organism database to build a knowledgebase supporting research on the genomics and biology of echinoderms. *Nucleic Acids Res.* 50:D970–D979.

Comte N, Morel B, Hasić D, Guéguen L, Boussau B, Daubin V, Penel S, Scornavacca C, Gouy M, Stamatakis A, et al. 2020. Treerecs: an integrated phylogenetic tool, from sequences to reconciliations. *Bioinformatics* 36:4822–4824.

Davidson PL, Guo H, Swart JS, Massri AJ, Edgar A, Wang L, Berrio A, Devens HR, Koop D, Cisternas P, et al. 2022. Recent reconfiguration of an ancient developmental gene regulatory network in Heliocidaris sea urchins. *Nat Ecol Evol* 6:1907–1920.

Davidson PL, Guo H, Wang L, Berrio A, Zhang H, Chang Y, Soborowski AL, McClay DR, Fan G, Wray GA. 2020. Chromosomal-Level Genome Assembly of the Sea Urchin Lytechinus variegatus Substantially Improves Functional Genomic Analyses. *Genome Biol. Evol.* 12:1080–1086.

Delroisse J, Ullrich-Lüter E, Blaue S, Ortega-Martinez O, Eeckhaut I, Flammang P, Mallefet J. 2017. A puzzling homology: a brittle star using a putative cnidarian-type luciferase for bioluminescence. *Open Biol.* [Internet] 7. Available from: http://dx.doi.org/10.1098/rsob.160300

Delroisse J, Ullrich-Lüter E, Ortega-Martinez O, Dupont S, Arnone M-I, Mallefet J, Flammang P. 2014. High opsin diversity in a non-visual infaunal brittle star. *BMC Genomics* 15:1035.

Dylus DV, Czarkwiani A, Stångberg J, Ortega-Martinez O, Dupont S, Oliveri P. 2016. Large-scale gene expression study in the ophiuroid Amphiura filiformis provides insights into evolution of gene regulatory networks. *Evodevo* 7:2.

Ketchum RN, Davidson PL, Smith EG, Wray GA, Burt JA, Ryan JF, Reitzel AM. 2022. A Chromosome-level Genome Assembly of the Highly Heterozygous Sea Urchin Echinometra sp. EZ Reveals Adaptation in the Regulatory Regions of Stress Response Genes. *Genome Biol. Evol.* [Internet] 14. Available from: http://dx.doi.org/10.1093/gbe/evac144

Kozlov AM, Darriba D, Flouri T, Morel B, Stamatakis A. 2019. RAxML-NG: a fast, scalable and user-friendly tool for maximum likelihood phylogenetic inference. *Bioinformatics* 35:4453–4455.

Lee Y, Kim B, Jung J, Koh B, Jhang SY, Ban C, Chi W-J, Kim S, Yu J. 2022. Chromosome-level genome assembly of Plazaster borealis sheds light on the morphogenesis of multiarmed starfish and its regenerative capacity. *Gigascience* [Internet] 11. Available from: http://dx.doi.org/10.1093/gigascience/giac063

Liu J, Zhou Y, Pu Y, Zhang H. 2023. A chromosome-level genome assembly of a deep-sea starfish (Zoroaster cf. ophiactis). *Sci Data* 10:506.

Long KA, Nossa CW, Sewell MA, Putnam NH, Ryan JF. 2016. Low coverage sequencing of three echinoderm genomes: the brittle star Ophionereis fasciata, the sea star Patiriella regularis, and the sea cucumber Australostichopus mollis. *Gigascience* 5:20.

Marlétaz F, Couloux A, Poulain J, Labadie K, Da Silva C, Mangenot S, Noel B, Poustka AJ, Dru P, Pegueroles C, et al. 2023. Analysis of the P. lividus sea urchin genome highlights contrasting trends of genomic and regulatory evolution in deuterostomes. *Cell Genom* 3:100295.

Mashanov V, Machado DJ, Reid R, Brouwer C, Kofsky J, Janies DA. 2022. Twinkle twinkle brittle star: the draft genome of Ophioderma brevispinum (Echinodermata: Ophiuroidea) as a resource for regeneration research. *BMC Genomics* 23:574.

Robinson MD, McCarthy DJ, Smyth GK. 2010. edgeR: a Bioconductor package for differential expression analysis of digital gene expression data. *Bioinformatics* 26:139–140.

Schiebelhut LM, Puritz JB, Dawson MN. 2018. Decimation by sea star wasting disease and rapid genetic change in a keystone species, *Pisaster ochraceus*. *Proc. Natl. Acad. Sci. U. S. A.* 115:7069–7074.

Sea Urchin Genome Sequencing Consortium, Sodergren E, Weinstock GM, Davidson EH, Cameron RA, Gibbs RA, Angerer RC, Angerer LM, Arnone MI, Burgess DR, et al. 2006. The genome of the sea urchin Strongylocentrotus purpuratus. *Science* 314:941–952.

Sinigaglia C, Almazán A, Lebel M, Sémon M, Gillet B, Hughes S, Edsinger E, Averof M, Paris M. 2022. Distinct gene expression dynamics in developing and regenerating crustacean limbs. *Proc. Natl. Acad. Sci. U. S. A.* 119:e2119297119.

Stewart R, Rascón CA, Tian S, Nie J, Barry C, Chu L-F, Ardalani H, Wagner RJ, Probasco MD, Bolin JM, et al. 2013. Comparative RNA-seq analysis in the unsequenced axolotl: the oncogene burst highlights early gene expression in the blastema. *PLoS Comput. Biol.* 9:e1002936.

Yamazaki A, Morino Y, Urata M, Yamaguchi M, Minokawa T, Furukawa R, Kondo M, Wada H. 2020. pmar1/phb homeobox genes and the evolution of the double-negative gate for endomesoderm specification in echinoderms. *Development* [Internet] 147. Available from: http://dx.doi.org/10.1242/dev.182139

Yan H, Bombarely A, Li S. 2020. DeepTE: a computational method for de novo classification of transposons with convolutional neural network. *Bioinformatics* 36:4269–4275.
